# Serine ADP-ribosylation marks nucleosomes for ALC1-dependent chromatin remodeling

**DOI:** 10.1101/2021.07.06.449314

**Authors:** Jugal Mohapatra, Kyuto Tashiro, Ryan L. Beckner, Jorge Sierra, Jessica A. Kilgore, Noelle S. Williams, Glen Liszczak

## Abstract

Serine ADP-ribosylation (ADPr) is a DNA damage-induced post-translational modification catalyzed by the PARP1/2:HPF1 complex. As the list of PARP1/2:HPF1 substrates continues to expand, there is a need for technologies to prepare mono- and poly-ADP-ribosylated proteins for biochemical interrogation. Here we investigate the unique peptide ADPr activities catalyzed by PARP1 in the absence and presence of HPF1. We then exploit these activities to develop a method that facilitates installation of ADP-ribose polymers onto full-length proteins with precise control over chain length and modification site. A series of semi-synthetic ADP-ribosylated histone proteins are prepared which demonstrate that ADPr at H2BS6 or H3S10 converts nucleosomes into robust substrates for the chromatin remodeler ALC1. Importantly, we found ALC1 selectively remodels ‘activated’ substrates within heterogeneous nucleosome populations and that nucleosome serine ADPr is sufficient to stimulate ALC1 activity in nuclear extracts. Our study identifies a biochemical function for nucleosome serine ADPr and describes a method that is broadly applicable to explore the impact that site-specific serine mono- and poly-ADPr have on protein function.

## Introduction

Protein ADP-ribosylation (ADPr) has been implicated in diverse mammalian cellular signaling pathways(Gupte et al., 2017). In this process, the ADP-ribose moiety from an NAD^+^ co-factor is deposited onto one of several chemically distinct amino acid side chain functionalities(Daniels et al., 2015). In cells, proteins can be modified with a mono-ADP-ribose adduct or variable length ADP-ribose polymers that emanate from specific protein sites, a process henceforth referred to as poly-ADPr. Among the 17-member poly(ADP-ribose) polymerase (PARP) enzyme family, PARP1/2 have emerged as the most extensively studied owing to the success of PARP1/2 inhibitors to treat DNA repair-deficient cancers(Lord and Ashworth, 2017). As the clinical utility of PARP1/2 inhibitors continues to expand, it is critical to understand how PARP1/2-dependent ADPr impacts cellular physiology and disease. In light of intense PARP1/2 substrate identification efforts(Bonfiglio et al., 2017; Larsen et al., 2018; Leidecker et al., 2016), several creative methods have been developed to install serine mono-ADPr onto synthetic peptides for biochemical interrogation(Bonfiglio et al., 2020; Voorneveld et al., 2018; Zhu et al., 2020). However, these technologies have been limited to relatively short peptide constructs. Additionally, no methods exist to reconstitute well-defined ADP-ribose chains at specific sites on isolated proteins for functional analysis. Hence, there is a dearth of mechanistic insight into how specific PARP1/2:HPF1-dependent mono- and poly-ADPr events regulate protein function.

Upon binding to single or double-stranded DNA breaks, PARP1/2 undergo conformational changes that induce the formation of a catalytically competent complex with NAD^+^ and the PARP1/2-interacting protein HPF1(Benjamin and Gill, 1980; Dawicki-McKenna et al., 2015; Gibbs-Seymour et al., 2016; Langelier et al., 2012; Suskiewicz et al., 2020). It has long been appreciated that DNA damage-induced ADPr has a profound effect on chromatin architecture through a variety of proposed mechanisms(Poirier et al., 1982; Ray Chaudhuri and Nussenzweig, 2017; Tulin and Spradling, 2003). Indeed, there are several ATP-dependent chromatin remodeling enzymes that localize to damage sites in an ADPr-dependent manner and contribute to decompaction of higher order chromatin structure, ultimately increasing repair factor accessibility(Ahel et al., 2009; Chou et al., 2010; Luijsterburg et al., 2016; Smeenk et al., 2013). One such chromatin remodeler, ALC1, harbors a macrodomain module that has been shown to specifically interact with tri-ADP-ribose(Singh et al., 2017). This binding event relieves an autoinhibited ALC1 conformation and activates the ATPase domain that powers nucleosome remodeling(Lehmann et al., 2017; Singh et al., 2017). ALC1 activation via ternary complex formation with auto-ADP-ribosylated PARP1 and nucleosomes has been extensively studied(Gottschalk et al., 2009; Gottschalk et al., 2012; Lehmann et al., 2017; Singh et al., 2017), and it has been suggested that other DNA-bound, ADP-ribosylated proteins may contribute to this process. However, it remains unclear which PARP1/2:HPF1 substrates and corresponding modification sites can lead to ALC1 activation, and if any are sufficient to do so in the absence of auto-modified PARP1. Such questions surrounding ALC1 regulation are increasingly important as recent studies show that abrogating ALC1 activity vastly increases the efficacy of PARP inhibitors(Blessing et al., 2020; Verma et al., 2021) and may even be useful for treatment of PARP inhibitor-resistant cancers(Juhasz et al., 2020).

The core histones H2B and H3 are consistently identified as some of the most abundantly modified PARP1/2:HPF1 substrates(Bonfiglio et al., 2017; Huletsky et al., 1989; Larsen et al., 2018). While much effort has been directed towards deciphering the regulatory mechanisms that govern serine ADPr(Bilokapic et al., 2020; Bonfiglio et al., 2017; Bonfiglio et al., 2020; Gibbs-Seymour et al., 2016; Palazzo et al., 2018; Suskiewicz et al., 2020), the functional consequences of specific nucleosome serine ADPr sites remain unclear. We and others have demonstrated that histone H2B serine 6 (H2BS6) and histone H3 serine 10 (H3S10) are the primary PARP1/2:HPF1 target sites in biochemical and cellular systems(Liszczak et al., 2018; Palazzo et al., 2018). Building upon these studies, we sought to determine how mono- and poly-ADPr on H2BS6 and H3S10 contribute to PARP1/2-dependent DNA repair activities such as ATP-dependent chromatin remodeling.

Here we employ an HPLC/MS-based analysis to investigate PARP1-dependent peptide ADPr activity in the absence and presence of HPF1. Reaction analyses guided the development of an approach that combines peptide chemistry, enzymatic catalysis, and protein ligation technologies to generate full-length proteins that bear mono- or poly-ADPr at user-defined serine sites. Key to this method is the separation of two enzyme-based peptide modification steps: 1.) mono-ADPr of unmodified peptides by the PARP1:HPF1 complex, and 2.) ADP-ribose chain elongation from mono-ADP-ribosylated peptides by the uncomplexed PARP1 enzyme. We prepare eight unique, semi-synthetic ADP-ribosylated nucleosomes and demonstrate that histone serine poly-ADPr marks nucleosomes for ALC1-dependent chromatin remodeling, with ALC1 activation levels of up to ∼370-fold observed relative to unmodified nucleosome substrates. Additional data support a model wherein nucleosome serine ADPr is sufficient to initiate ALC1-dependent chromatin structure alterations with a high degree of spatial precision. This study describes a broadly applicable method to install ADP-ribose chains at specific PARP1/2:HPF1 target sites on peptides and proteins and identifies a functional output for nucleosome serine ADPr in the DNA damage response.

## Results

### An HPLC/MS-based approach to analyze peptide ADPr by PARP1:HPF1

While synthetic and enzyme-based methodologies exist to prepare mono-ADP-ribosylated peptide fragments(Bonfiglio et al., 2020; Voorneveld et al., 2018; Zhu et al., 2020), installation of poly-ADP-ribose is synthetically more complex and has not been reported. Therefore, we envisioned an enzyme-based approach that employs the PARP1:HPF1 complex to modify specific serine sites on synthetic peptides with homogenous ADP-ribose polymers. A similar elegant approach was recently reported by the Matic group to prepare mono-ADP-ribosylated peptides, which included H2B and H3 tail constructs(Bonfiglio et al., 2020). However, in that study, a post-reaction poly-ADP-ribose glycohydrolase (PARG) treatment was carried out to reduce any poly-ADP-ribosylated species to the mono-ADP-ribose adduct. Our method is unique in that we developed an RP-HPLC-MS-based assay to simultaneously monitor recombinant PARP1:HPF1 complex activity on a peptide substrate and separate distinct mono- and poly-ADP-ribosylated peptide products (Fig. 1a).

**Fig. 1:**
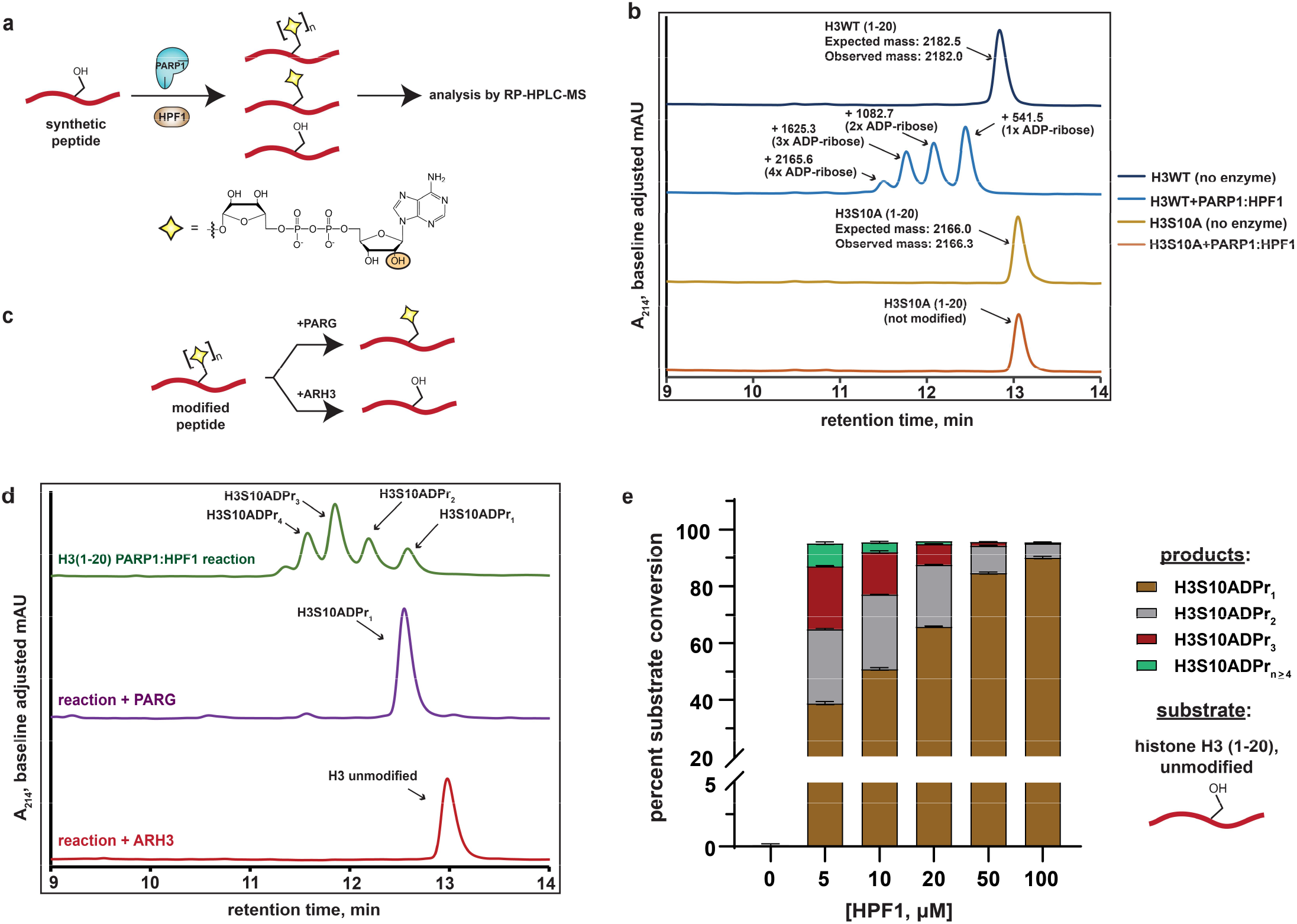
Analysis of serine mono- and poly-ADPr by the PARP1:HPF1 complex on synthetic peptide substrates. **a**, A schematic showing the workflow employed to analyze peptide poly-APDr by the recombinant PARP1:HPF1 complex. Peptide products are separated by polymer length via RP-HPLC. The yellow star represents a serine-linked ADP-ribose modification, ‘n’ represents variable polymer length, and the orange circle indicates the site of linear ADP-ribose polymerization. **b**, RP-HPLC and MS analysis of substrate peptides (histone H3 wild-type or S10A mutant, amino acids 1-20) and corresponding PARP1:HPF1 reaction products (for raw MS data, see Supplementary Fig. 1a). RP-HPLC gradients are from 0-35% Solvent B (2-22 min). **c**, A schematic describing the ADP-ribosylhydrolase-based characterization strategy. Enzymes and their respective reaction products are depicted. **d**, RP-HPLC traces from PARG- or ARH3-treated H3 peptide ADPr reactions that were optimized for ADP-ribose chain elongation. The number of ADP-ribose units was verified by MS analysis. **e**, Product analysis of a PARP1 ADPr reaction in the presence of increasing HPF1 concentrations. Histone H3 substrate peptide starting material and each unique ADP-ribosylated product were quantified via HPLC chromatogram peak integration (see Methods and Supplementary Fig. 1e). The columns represent the percent substrate conversion to each ADP-ribosylated product. Data are represented as mean ± s.d. (n = 3).

We began our study by incubating a synthetic histone H3 peptide (amino acids 1-20) that contains a single known serine target site (H3S10) with the PARP1:HPF1 complex, NAD^+^, and stimulating DNA. Multiple H3 peptide product peaks were observed via chromatography-based reaction analysis. ESI-MS characterization revealed a single, unique mass in each HPLC product peak, which corresponded to an H3 peptide modified with mono-, di-, tri-, or tetra-ADP-ribose (henceforth H3S10ADPr_n_) (Fig. 1b and Supplementary Fig. 1a). Notably, all products are sensitive to the H3S10A mutation, indicating the presence of an ADP-ribose chain that elongates from the S10 site (Fig. 1b). Thus, each individual peptide product corresponding to mono-, di-, tri-, or tetra-ADP-ribosylated H3S10 can be separated via RP-HPLC.

We next treated ADPr reactions with recombinant ADP-ribosylhydrolase enzymes to validate the modification site and chemical identity of modified peptide products (Fig. 1c). Analysis via HPLC-MS demonstrates that PARG(Slade et al., 2011) treatment quantitatively converts all observed ADP-ribosylated H3 peptide products to the mono-ADP-ribosylated species, which is consistent with a single modification site (Fig. 1d). When the serine-specific ADP-ribosylhydrolase 3 (ARH3)(Fontana et al., 2017) enzyme is substituted for PARG, all ADP-ribosylated species are converted to the unmodified H3 peptide, thus confirming a serine-linked modification (Fig. 1d). An established LC-MS/MS analysis protocol(Chen et al., 2018) was used to determine that the peptide-linked ADP-ribose chains were principally linear, with negligible branching (< 0.03%) (Supplementary Fig. 1b).

The workflow and characterization strategies described here were next implemented to install ADP-ribose chains at the known PARP1:HPF1 target site on a synthetic H2B peptide (amino acids 1-16). Despite the presence of two serine residues in the H2B peptide, our mutagenesis and ADP-ribosylhydrolase-based characterizations confirmed H2BS6 as the sole acceptor residue (Supplementary Fig. 1c and d). Notably, while conversion of up to 1 mM (∼20 mg) of unmodified H2B or H3 peptides to the H2BS6ADPr_1_ or H3S10ADPr_1_ products could be routinely achieved, a more scalable approach for peptide poly-ADPr would be required to deploy these molecules in protein ligation reactions and biochemical assays.

Analysis of the PARP2:HPF1 structure suggests that HPF1 binding, while required for serine ADPr, would interfere with the PARP1/2 ADP-ribose chain elongation mechanism(Suskiewicz et al., 2020). This observation is consistent with several recent reports that show HPF1-dependent shortening of PARP1/2-catalyzed ADP-ribose chains in cellular and biochemical assays(Bonfiglio et al., 2020; Gibbs-Seymour et al., 2016; Rudolph et al., 2021). We therefore hypothesized that the concentration of HPF1 in the peptide modification reaction may affect the final distribution of our mono- and poly-ADP-ribosylated peptide products. To explore this, an HPF1 titration from 5 μM to 100 μM was performed in an ADPr reaction containing the H3 peptide. Notably, unmodified peptide starting material and ADP-ribosylated peptide products could be separated via RP-HPLC and quantified by chromatogram peak integration at A_214_ and A_280_, respectively (see Methods for details). Near quantitative conversion (>95%) of the unmodified H3 substrate to ADP-ribosylated products was achieved at HPF1 concentrations as low as 5 μM (Fig. 1e). Interestingly, we observed a gradual increase in mono-ADPr activity and decrease in poly-ADPr activity as HPF1 is titrated into the reaction (Fig. 1e and Supplementary Fig. 1e). In the 5 μM HPF1 reaction, the mono-ADP-ribosylated peptide represents ∼41% of the total product, with the remaining ∼59% comprising a distribution of di- to penta-ADP-ribosylated peptide. In the 100 μM HPF1 reaction, mono-ADP-ribosylated peptide increases to ∼94% of the total product, with di-ADP-ribose representing the remaining ∼6%. This is consistent with a mechanism wherein PARP1:HPF1 complex formation switches PARP1 activity from an ADP-ribose chain elongator to a mono-ADP-ribosyltransferase. Indeed, these experimental data are congruent with the structure-based hypothesis put forth by Suskiewicz, et al. that HPF1 limits PARP1/2 activity to mono-ADPr.

### Synthesis of poly-ADP-ribosylated peptides via two enzymatic steps

Based on the mechanistic interpretation described above, we surmised that PARP1 would display efficient ADP-ribose chain elongation activity on mono-ADP-ribosylated peptides in the absence of HPF1 in our reconstituted system. To investigate this, we employed our purified H3S10ADPr_1_ peptide as a substrate in a PARP1 activity assay that lack HPF1 (Fig. 2a). Importantly, we maintained all reaction conditions, substrate concentrations, and stimulating DNA concentrations described for the PARP1:HPF1 activity assays. Strikingly, incubation of the H3S10ADPr_1_ peptide with PARP1 resulted in robust ADP-ribose chain elongation at all enzyme concentrations tested (0.2, 1, and 5 μM). Nearly 70% conversion of the H3S10ADPr_1_ substrate to poly-ADP-ribosylated products was achieved at 1 μM PARP1 (Fig. 2b and Supplementary Fig. 2a). The di-, tri-, and tetra-ADP-ribosylated species were the most abundant products with yield decreasing precipitously for chains greater than four units in length (Supplementary Fig. 2a). Notably, PARP2 also catalyzes ADP-ribose chain elongation from the H3S10ADPr_1_ substrate and similar polymerization activity was observed with both PARP1 and PARP2 on the H2BS6ADPr_1_ substrate (Fig. 2b and Supplementary Fig. 2b and c).

**Fig. 2:**
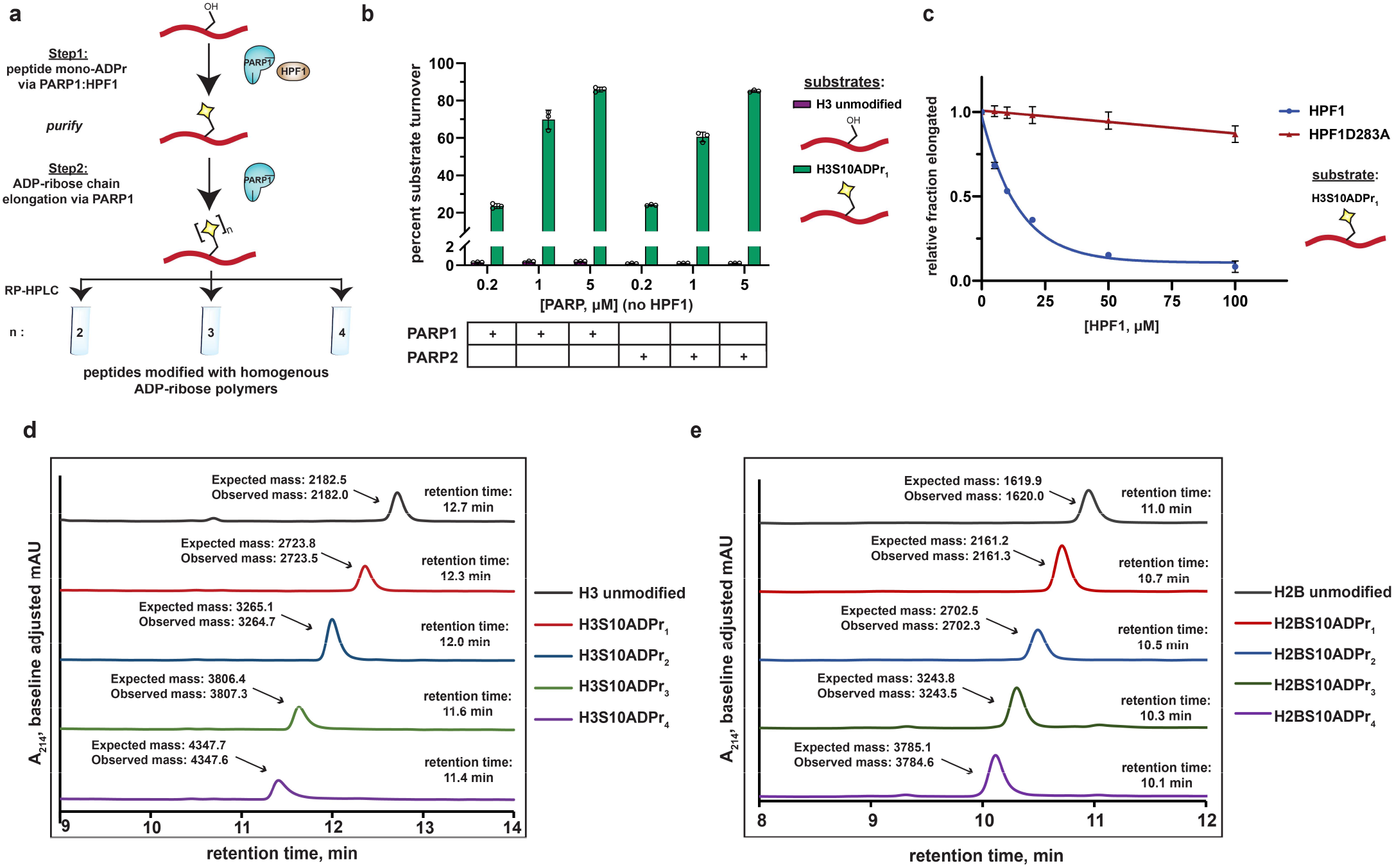
A two-step enzymatic process to prepare poly-ADP-ribosylated peptides with defined ADP-ribose chain lengths. **a**, A schematic showing the two-step enzymatic procedure implemented to synthesize and purify poly-ADP-ribosylated peptides. The mono-ADP-ribosylated peptide product from Step 1 was purified using preparative RP-HPLC prior to use in Step 2. **b**, Substrate turnover analysis of PARP1 and PARP2 ADPr reactions in the absence of HPF1. Purple bars represent total percent turnover of an unmodified H3 peptide to mono- or poly-ADP-ribosylated products. Green bars represent total percent turnover of the H3S10ADPr_1_ peptide to poly-ADP-ribosylated products (for poly-ADP-ribosylated product distribution, see Supplementary Fig. 2a and b) Data are represented as mean ± s.d. (n = 3). **c**, Analysis of PARP1 elongation activity on the H3S10ADPr_1_ peptide substrate in the presence of increasing amounts of HPF1 or HPF1D283A. Fraction elongated represents the fraction of H3S10ADPr_1_ peptide converted to poly-ADP-ribosylated products. Data are normalized to fraction of substrate elongated in the absence of HPF1. Data are represented as mean ± s.d. (n = 3). The curves represent the fit of the data into a non-linear regression model for one-phase exponential decay. **d**, RP-HPLC and MS analysis of mono- and poly-ADP-ribosylated H3 peptides that have been purified to homogeneity via semi-preparative HPLC. **e**, As in **d**, but for H2B (amino acids 1-16) peptides.

To further characterize the inhibitory effect that HPF1 has on PARP1-dependent ADP-ribose chain elongation, we incubated PARP1 with the H3S10ADPr_1_ substrate peptide in the presence of increasing concentrations of HPF1. As expected, HPF1 exhibits dose-dependent inhibition of PARP1-catalyzed ADP-ribose polymerization from the mono-ADP-ribosylated substrate, with 50% inhibition occurring at ∼14 μM HPF1 for 1 μM PARP1. A binding-deficient HPF1 mutant (D283A)(Rudolph et al., 2021; Suskiewicz et al., 2020) is unable to appreciably inhibit ADP-ribose polymerization (Fig. 2c and Supplementary Fig. 2d and e). These data complement our unmodified peptide substrate:HPF1 titration analysis and provide additional evidence that the PARP1:HPF1 complex is a dedicated mono-ADP-ribosyltransferase.

Importantly, by first isolating mono-ADP-ribosylated peptides from a PARP1:HPF1 reaction for use in a PARP1 elongation reaction, each poly-ADP-ribosylated H2BS6 and H3S10 product (up to four ADP-ribose units in length) could now be purified to homogeneity in milligram quantities for downstream applications (Fig. 2d and e). The broad applicability our peptide poly-ADPr strategy was further validated with additional known PARP1:HPF1 target sequences(Bonfiglio et al., 2020) including TMA16 (amino acids 2-19, target residue S9), a fragment of the PARP1 automodification domain (amino acids 501-515, target residue S507), and a secondary histone H3 site (amino acids 21-34; target residue S28). The mono-, di-, tri-, and tetra-ADP-ribosylated species were isolated for each of these peptides (Supplementary Fig. 2f and g). Thus, PARP1 can dependably elongate ADP-ribose chains from peptides that have been ‘primed’ with serine mono-ADP-ribose by PARP1:HPF1. We do note that overall poly-ADP-ribosylated product yields vary depending upon target peptide identity, but all reactions could be optimized to obtain milligram quantities of each unique product (see ‘Methods’ for details).

### ADP-ribosylated H2B and H3 peptides engage the ALC1 macrodomain with equal affinity

Extensive precedent exists demonstrating that chromatin remodeling enzymes are regulated by modifications on the nucleosome substrate(Clapier and Cairns, 2012; Hauk et al., 2010). The Ladurner lab recently reported that the ALC1 macrodomain exhibits high affinity (K_d_ ∼ 10 nM) for free tri-ADP-ribose with little to no binding detectable for free mono- and di-ADP-ribose molecules(Singh et al., 2017). We therefore chose to pursue ALC1 for our initial ADP-ribosylated histone peptide interaction studies. Nine fluorescently-labeled, ADP-ribosylated histone peptides (H2BS6ADPr_1-4_ and H3S10ADPr_1-5_) were prepared for fluorescence polarization-based interaction assays (Supplementary Fig. 3). We note that the ADP-ribose polymerization reaction is more efficient with the H3 peptide and hence longer peptide-conjugated ADP-ribose chains could be isolated relative to H2B. Initial assay development was carried out by titrating a commercially available pan-ADP-ribose detection reagent (an Af1521 macrodomain-Fc region fusion)(Gibson et al., 2017) into each peptide. This reagent exhibits ADPr-dependent binding for all H2B and H3 peptides, with affinity decreasing precipitously for chains less than three ADP-ribose units in length (Fig. 3a, b, and Supplementary Table 1).

**Fig. 3:**
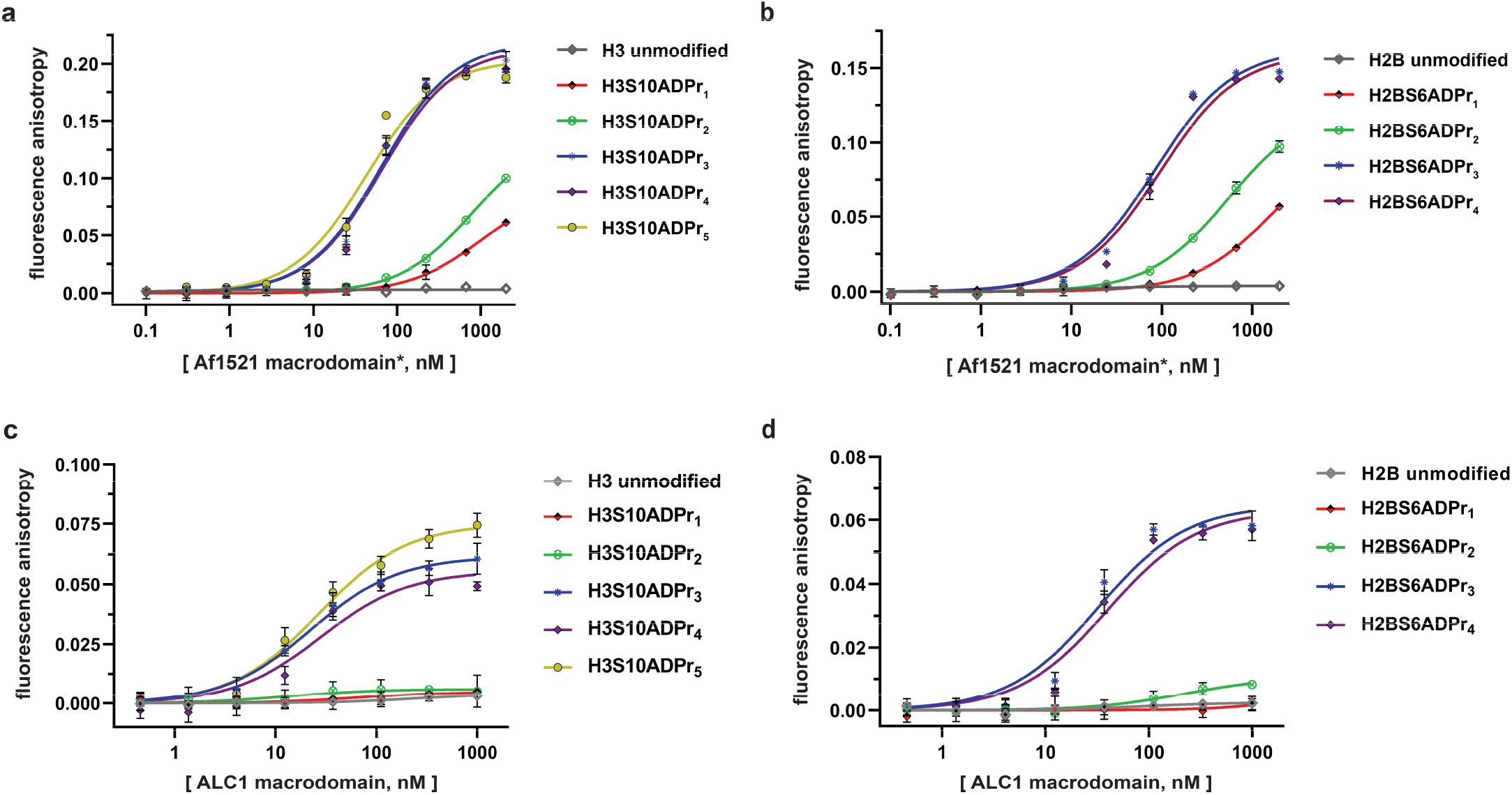
The ALC1 macrodomain engages ADP-ribosylated H2B and H3 peptides with equal affinity. **a**, Fluorescence polarization (FP) assays to evaluate binding affinities of different ADP-ribosylated, fluorescein-labeled H3 (1-20) peptides to the Af1521 macrodomain. Data are represented as mean ± s.d. (n = 3). All curves represent fit of the data into a non-linear regression equation for one-site, specific binding (for K_d, app_ values, see Supplementary Table 1). *The Af1521 macrodomain is from the commercially available pan-ADP-ribose detection reagent. **b**, As in **a**, but with H2B (1-16) peptides. **c**, FP assays as described in **a** to evaluate binding affinities of ADP-ribosylated, fluorescein-labeled H3 (1-20) peptides to the ALC1 macrodomain. **d**, As in **c**, but with H2B (1-16) peptides.

Similar experiments were performed by titrating the ALC1 macrodomain into each fluorescently-labeled histone peptide for apparent dissociation constant (K_d, app_) calculations. Consistent with free ADP-ribose binding preferences(Singh et al., 2017), the mono- and di-ADP-ribosylated H2B and H3 peptides failed to appreciably interact with the ALC1 macrodomain. Contrastingly, all tri-, tetra-, and penta-ADP-ribosylated peptides are high-affinity ligands with K_d, app_ ranging from ∼21-37 nM (Fig. 3c, d, and Supplementary Table 1). Considering the H2BS6ADPr_3-4_ and H3S10ADPr_3-5_ peptides exhibit similar affinities, we concluded that the tri-ADP-ribose modification is likely sufficient for optimal ALC1 macrodomain:peptide engagement. These data also indicate that while the ALC1 macrodomain engages the H2BS6 and H3S10-modified peptides, it does not exhibit sequence-based preference for either site.

### Preparation of full-length, homogenously ADP-ribosylated histone proteins and assembly into nucleosomes

Chromatin remodelers comprise multiple domains that function synergistically to recognize nucleosome substrates and mobilize histone proteins(Bowman and Poirier, 2015). This phenomenon implies that macrodomain-ligand specificity may not represent the sole determinant of ALC1 substrate preference. To address this, we sought to analyze full-length ALC1 remodeling activity in the context of ADP-ribosylated nucleosome substrates. The first step towards reconstituting modified nucleosomes requires preparation of full-length, ADP-ribosylated histones. We generated a series of ADP-ribosylated H2B and H3 peptides with C-terminal thioesters to enable an eventual native chemical ligation reaction to the remainder of the corresponding histone fragment (Fig. 4a). The following six semi-synthetic, full-length histones were prepared: H2BS6ADPr_1_, H2BS6ADPr_3_, H2BS6ADPr_4_, H3S10ADPr_1_, H3S10ADPr_3_, and H3S10ADPr_4_ (Supplementary Fig. 4). The tri- and tetra-ADP-ribosylated H2B and H3 proteins were essential to probe the effect of chain length and nucleosome modification site on ALC1 activation. Mono-ADP-ribosylated histones were prepared to serve as negative controls and to further corroborate ALC1 macrodomain interaction results. All final protein products were characterized via HPLC/MS analysis and determined to be >95% pure, hence validating our workflow to reconstitute homogenously ADP-ribosylated proteins (Fig. 4b and Supplementary Fig. 4).

**Fig. 4:**
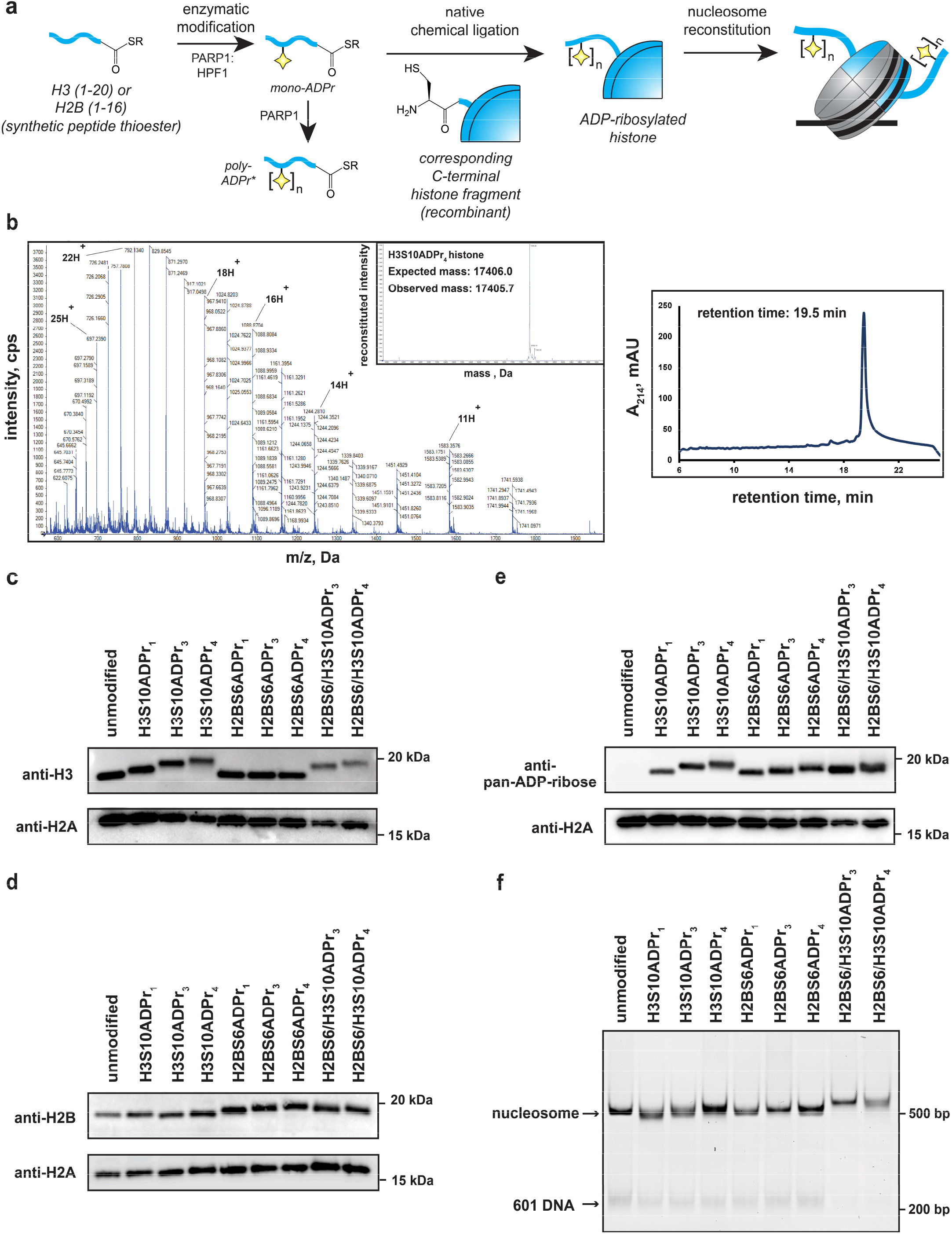
Installation of homogenous ADP-ribose polymers onto reconstituted nucleosomes via a chemoenzymatic strategy. **a**, A schematic depicting the protein semi-synthesis-based strategy to install homogenous ADP-ribose polymers at specific sites on histone proteins. The nucleosome cartoon includes DNA (black line), as well as the histone protein octamer core (grey = recombinant histones, blue = semi-synthetic histone). *The poly-ADP-ribosylated peptides are separated via HPLC to yield homogenous species prior to the ligation reaction. **b**, Representative HPLC/MS characterization of the full-length H3S10ADP_4_ protein. Raw ESI-MS spectra, MS deconvolution, and RP-HPLC chromatogram are shown. RP-HPLC gradients are from 0-80% Solvent B (2-22 min). For additional histone HPLC and MS characterizations, see Supplemental Fig. 4. **c**, Western blot analysis of histone H3 following nucleosome assembly. ADP-ribose-dependent gel migration shifts demonstrate sample homogeneity. **d**, Histone H2B analysis as described in panel **c**. **e**, Pan-ADP-ribose detection western blot analysis of all assembled nucleosomes. **f**, Native gel analysis of assembled nucleosomes. Single nucleosome bands and trace levels of free 601 DNA demonstrate sample homogeneity and assembly efficiency. EtBr = ethidium bromide stain.

Each of the six semi-synthetic ADP-ribosylated histones were combined with the necessary recombinant histones to form stable histone octamer complexes (henceforth labeled as H2BS6ADPr_n_ or H3S10ADPr_n_, depending on the modified histone they possess) via established protocols(Luger et al., 1999). We also prepared an octamer that contains both H2BS6ADPr_3_ and H3S10ADPr_3_ (H2BS6/H3S10ADPr_3_), and another that contains both H2BS6ADPr_4_ and H3S10ADPr_4_ (H2BS6/H3S10ADPr_4_). Following purification via gel filtration chromatography, octamer quality and ADPr stability was determined via SDS-PAGE/western blot analysis. Histone detection via western blotting with H2B and H3 antibodies revealed single, distinct species for each ADP-ribosylated H2B and H3 histone (Fig. 4c and d). We found that ADP-ribose chain length is inversely proportional to histone gel migration distance, suggesting that single migration bands for H2B and H3 are a reliable indicator of modification stability and sample homogeneity. Additionally, all gel species that correspond to ADP-ribosylated histones exhibited strong signal in a pan-ADP-ribose detection blot (Fig. 4e). Next, the eight ADP-ribosylated octamers were assembled into unique nucleosomes using a DNA template that contains the ‘601’ nucleosome positioning sequence and is compatible with a previously reported restriction enzyme accessibility (REA)-based chromatin remodeling assay (see Methods for details)(He et al., 2006). Nucleosome quality was analyzed on a native polyacrylamide TBE gel, which shows a single, distinct nucleosome species for each assembly and only trace levels of free 601 DNA (Fig. 4f). Notably, ADP-ribose has a polymer length-dependent effect on nucleosome gel migration patterns, again indicating sample homogeneity and modification stability. We concluded that all of our site-specifically ADP-ribosylated histones could be efficiently incorporated into nucleosomes for downstream chromatin remodeling experiments.

### Serine ADPr converts nucleosomes into robust ALC1 substrates

Recombinant, full-length ALC1 was isolated to determine chromatin remodeling rate constants with each ADP-ribosylated nucleosome substrate. The DNA from each remodeling reaction was isolated at various time points and remodeling-dependent restriction enzyme cleavage was visualized on a polyacrylamide TBE gel and quantified via densitometry (Fig. 5a and Supplementary Fig. 5a). Consistent with the macrodomain interaction results, ALC1 exhibits relatively low remodeling rate constants (< 3×10^−4^ min^−1^) with unmodified and mono-ADP-ribosylated nucleosome substrates (Fig. 5b and Supplementary Table 2). Contrastingly, robust chromatin remodeling activity is observed with all nucleosomes that contain tri- or tetra-ADP-ribose at the H2B or H3 sites. The H2BS6/H3S10ADPr_4_ nucleosome has the most striking effect on the ALC1 remodeling rate constant, which increases ∼370-fold relative to the unmodified nucleosome. Further rate constant analyses show that ALC1 exhibits modest preference for the H2BS6 modification site and tetra-ADP-ribose polymers (Fig. 5b). Importantly, addition of H2BS6ADPr_4_ or H3S10ADPr_4_ peptide to a reaction containing ALC1 and unmodified nucleosome was unable to appreciably stimulate remodeling activity regardless of peptide concentration (Fig. 5c and Supplementary Fig. 5b and c). Therefore, in addition to disrupting an autoinhibited conformation, the modified histone tail:macrodomain interaction is crucial for presenting the ATPase domain to the nucleosome.

**Fig. 5:**
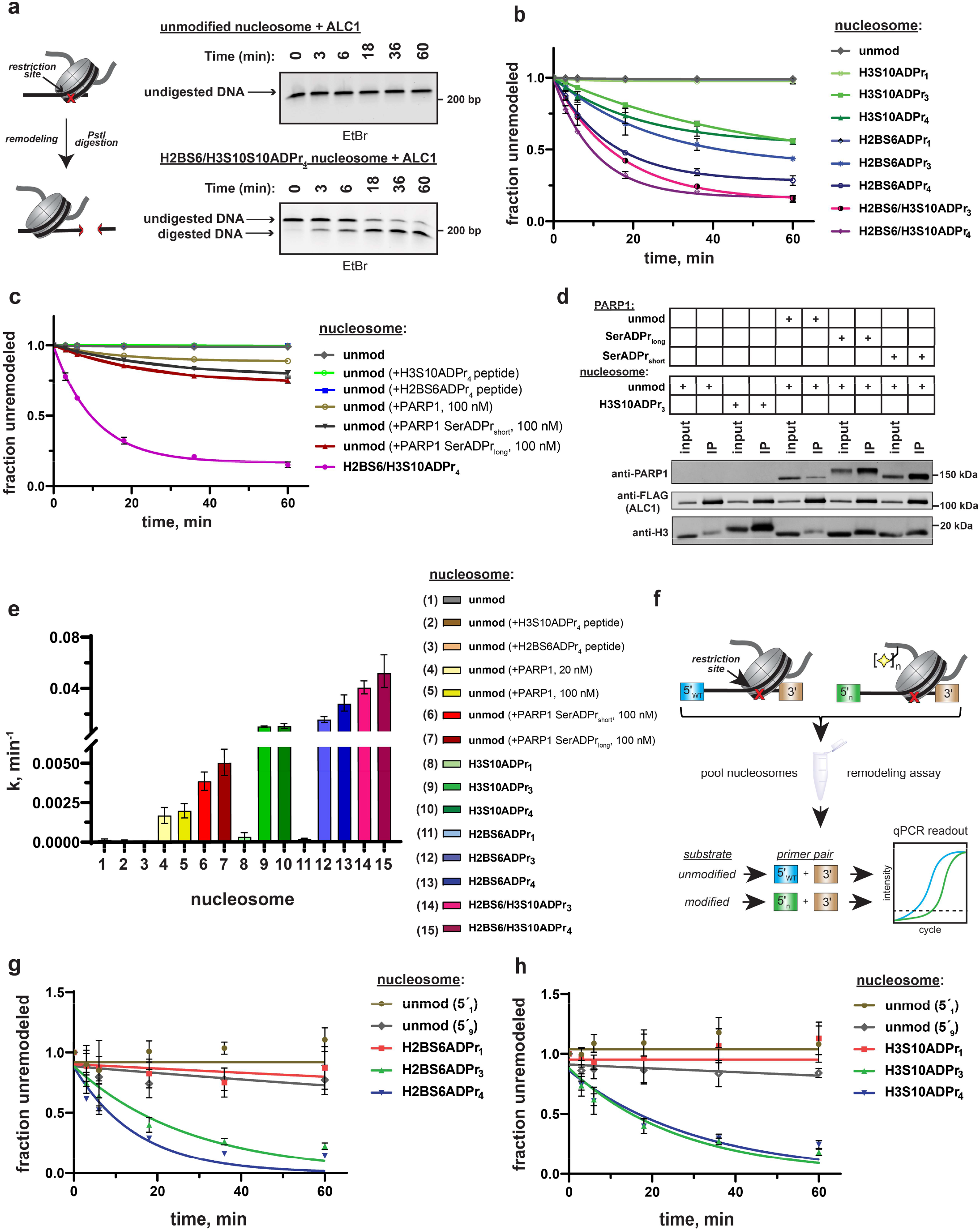
ADPr at H2BS6 and H3S10 convert nucleosomes into robust ALC1 substrates. **a**, Schematic depicting the REA assay for chromatin remodeling and representative TBE gel analyses of recombinant ALC1 activity on unmodified or H2BS6/H3S10ADPr_4_ nucleosomes. **b**, ALC1 nucleosome remodeling assay time-course wherein each reaction comprises ALC1 and the indicated nucleosome (‘unmod’= unmodified). **c**, As in **b**, but each reaction comprises ALC1, unmodified nucleosome (20 nM), and the indicated modified histone peptide or PARP1. Modified histone peptide concentration is equal to the corresponding full-length histone concentration (40 nM). The H2BS6/H3S10ADPr_4_ nucleosome remodeling data is included for direct comparison. **d**, Western blot analysis of a FLAG immunoprecipitation (IP) wherein ALC1 is FLAG-tagged and its association with nucleosomes is analyzed in the presence and absence of unmodified or automodified PARP1. The corresponding input (5%) was loaded alongside the IP (elution) lanes for comparison. **e**, ALC1 remodeling rate constants calculated from data in **b, c** and Supplementary Fig. 5b. Rate constants were determined by fitting data to a non-linear regression model for one phase exponential decay. **f**, Schematic depicting the strategy to prepare heterogenous nucleosome substrate pools and determine ALC1 remodeling activity on specific nucleosomes. **g**, ALC1 nucleosome remodeling assay time-course for each nucleosome in the histone H2B mixed substrate pool. Two unmodified nucleosomes with different 5′ primer sequences (5′_1_ and 5′_9_) were included as internal controls. **h**, As in **g**, but with the histone H3 substrate pool. Data in **b**, **c**, **e**, **g**, and **h** are represented as mean ± s.d. (n=3). Curves in **b**, **c**, **g** and **h** represent data fitting to a linear-regression model for one-phase exponential decay.

We next asked how ALC1 activation by nucleosome serine ADPr compares to activation by auto-ADP-ribosylated PARP1(Gottschalk et al., 2009; Gottschalk et al., 2012; Lehmann et al., 2017; Singh et al., 2017). As previously described, chromatin remodeling reactions were performed on unmodified nucleosome substrates in the presence of NAD^+^ and PARP1(Gottschalk et al., 2009; Gottschalk et al., 2012). In this experimental setup, PARP1 maintains auto-ADPr activity but is unable to modify histones due to absence of HPF1. Quantitative PARP1 auto-ADPr was observed within 5 min of initiating the reaction as judged by altered PARP1 gel migration in SDS-PAGE/western blot analyses (Supplementary Fig. 5d). PARP1 was added to the reaction at equimolar concentrations relative to nucleosome substrates 20 nM to closely mimic ADP-ribose concentrations in our modified nucleosome experiments or 100 nM to ensure optimal ALC1 activation. We found that auto-ADP-ribosylated PARP1 leads to an ∼12-fold increase in ALC1 remodeling rate constant on unmodified nucleosomes (Fig. 5c and Supplementary Table 2). Notably, higher PARP1 concentrations were unable to further stimulate ALC1 remodeling activity (Supplementary Fig. 5e).

In the PARP1 automodification reaction described above, aspartate and glutamate side chains are the primary targets for ADPr as no HPF1 is present. However, in the cellular DNA damage response, it is now well-established that auto-modification occurs primarily on serine residues(Bonfiglio et al., 2017; Palazzo et al., 2018). We therefore performed a PARP1 automodification reaction in the presence of low (5 µM) and high (25 µM) amounts of HPF1. By employing different HPF1 concentrations, a full-length PARP1 construct with relatively short (PARP1-SerADPr_short_) and long (PARP1-SerADPr_long_) serine-linked ADP-ribose chains could be generated. These constructs were purified over a heparin column to remove activating DNA and HPF1, which would otherwise abrogate the nucleosome interaction or induce histone ADPr, respectively. The auto-ADPr linkage identity was then validated via hydroxylamine treatment, which specifically cleaves ADPr from aspartate and glutamate side chains. As expected, the ADP-ribose chains conjugated to PARP1-SerADPr_short_ and PARP1-SerADPr_long_ are resistant to hydroxylamine cleavage (Supplementary Fig. 5f). Immunoprecipitations with Flag-tagged ALC1 revealed that PARP1-SerADPr_short_ and PARP1-SerADPr_long_ are able to induce formation of an ALC1:nucleosome:PARP1 complex (Fig. 5d). We then titrated each construct into an ALC1 remodeling reaction with unmodified nucleosomes and observed optimal remodeling stimulation at 100 nM of automodified PARP1 (Supplementary Fig. 5g). Remodeling rate constant calculations show that PARP1-SerADPr_short_ and PARP1-SerADPr_long_ stimulate ALC1 activity ∼28-fold and ∼36-fold, respectively, when compared to activity in the absence of automodified PARP1 (Fig. 5c and Supplementary Table 2). We stress that while nucleosome serine ADPr is superior to PARP1 auto-ADPr for ALC1 activation in biochemical assays (Fig. 5e), these data do not allow us to conclude that this is the case in the cellular DNA damage response. However, our work does raise interesting new questions about regulatory mechanisms underlying ALC1 activity (see Discussion).

### ALC1 specificity persists within mixed nucleosome pools

To further probe ALC1 nucleosome substrate selectivity, we designed a method to pool unmodified, mono, tri-, and tetra-ADP-ribosylated nucleosomes into a single reaction and analyze nucleosome remodeling activity for each unique substrate simultaneously (Fig. 5f). Similar next-generation sequencing-based approaches have been implemented for rate constant analysis of the ISWI chromatin remodeler family(Dann et al., 2017). If ALC1 activity is dependent upon the ADPr status of target nucleosomes, only the tri- and tetra-ADP-ribosylated species should be efficiently remodeled in this substrate competition-based platform. We again turned to the REA assay but appended a unique 5′ 15-base pair primer binding site to each 601 DNA template (Supplementary Table 3). Importantly, we designed priming sequences with similar primer binding efficiencies and found that DNA sequence alterations in this region of the template do not affect remodeling rates (Supplementary Fig. 5h). In this assay, restriction enzyme-dependent destruction of a given 601 template amplicon is quantified by qPCR to monitor remodeling activity. Thus, unique primer pairs corresponding to each nucleosome can be employed to determine substrate-specific chromatin remodeling rate constants in heterogenous substrate reactions.

We assembled a nucleosome pool comprising equimolar concentrations of H2BS6ADPr_1_, H2BS6ADPr_3_, H2BS6ADPr_4_, and two unmodified nucleosome controls. An additional unmodified nucleosome without the PstI restriction site and a free DNA template with the PstI site were also included as negative and positive digestion controls, respectively. The heterogeneous nucleosome substrate pool was employed in ALC1 remodeling reactions as described above, and DNA from various time points was isolated and analyzed via qPCR. We found that relative remodeling rate constants were consistent with those observed in our single substrate, densitometry-based assays (Fig. 5g and Supplementary Table 3). ALC1 again exhibits modest preference for the H2BS6ADPr_4_ nucleosome relative to the H2BS6ADPr_3_ nucleosome. Remodeling was very slow for the unmodified and H2BS6ADPr_1_ nucleosomes and corresponding rate constants could not be determined in this assay platform. Substrate preferences were also maintained within a similar H3S10-modified substrate pool (Fig. 5h and Supplementary Table 3). Notably, H3 nucleosomes were analyzed as a separate population because they require a higher ALC1 concentration to achieve optimal dynamic range in the qPCR-based assay. The pooled substrate approach demonstrates that ALC1 activity is highly specific for binding-competent nucleosome substrates and target disengagement triggers rapid transition back to an inactive conformation. This mechanism likely minimizes that likelihood that freely diffusing, activated ALC1 is present in the nuclear milieu.

### Nucleosome serine ADPr triggers ALC1-dependent chromatin remodeling in nuclear lysates

It is possible that a poly-anionic chain fused to H2BS6 or H3S10 destabilizes the histone octamer:DNA complex and thereby non-specifically sensitizes nucleosomes to ATP-dependent chromatin remodelers. To examine this concept, we isolated the ATP-dependent chromatin remodeler CHD4 for activity analysis. CHD4 lacks a macrodomain while its ATPase domain shares a high degree of sequence similarity (63%) with ALC1 (Supplementary Fig. 6a), suggesting that the two enzymes may catalyze DNA translocation through similar mechanistic principles. The REA assay revealed that CHD4 remodels unmodified nucleosomes with a rate constant of ∼0.01 min^−1^ and this activity is not appreciably affected by the nucleosome ADPr status (Fig. 6a, b, Supplementary Fig. 6b, and Supplementary Table 2). These data suggest that nucleosome serine ADPr does not simply decrease the energy barrier to DNA translocation but rather serves to specifically stimulate ALC1-dependent chromatin remodeling.

**Fig. 6:**
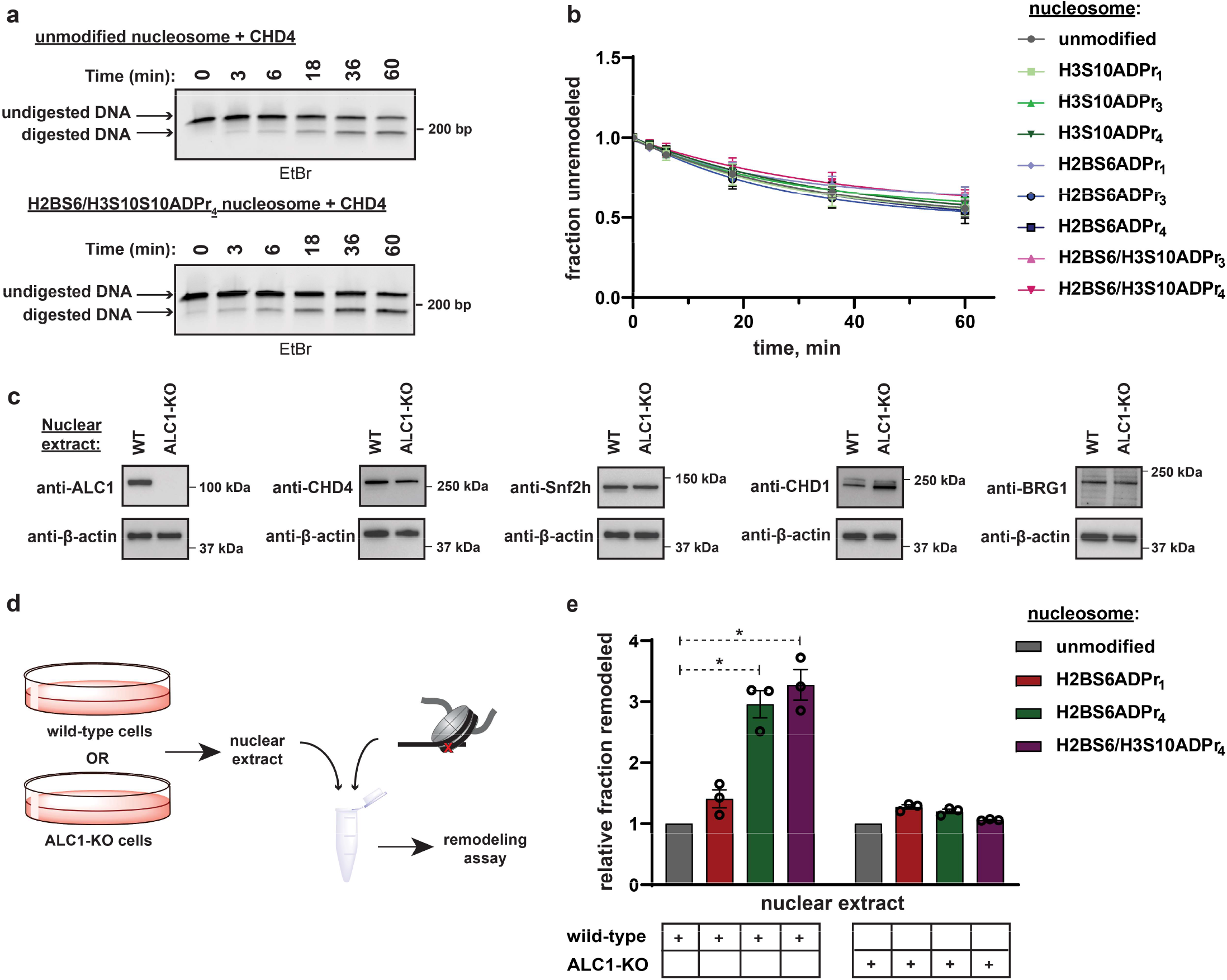
Nucleosome serine ADPr stimulates ALC1-dependent chromatin remodeling activity in nuclear lysates. **a**, Representative TBE gel analysis from a REA assay corresponding to recombinant CHD4 chromatin remodeling activity on unmodified or H2BS6/H3S10ADPr_4_ nucleosomes. **b**, CHD4 nucleosome remodeling assay time-course wherein each reaction comprises CHD4 and the indicated nucleosome substrate. Data are represented as mean ± s.d. (n = 3). Curves represent fit of data into a non-linear regression model for one-phase exponential decay. **c**, Western blot analysis demonstrating the presence of various chromatin remodelers in the wild-type or ALC1 knock-out (KO) HEK293T nuclear extracts. **d**, Schematic depicting the strategy to analyze chromatin remodeling activity in wild-type or ALC1-KO HEK293T nuclear extracts. **e**, Nuclear extract nucleosome remodeling activity assay wherein each reaction comprises the indicated nucleosome substrate and wild-type or ALC1-KO HEK293T cell nuclear extracts. Total remodeling for each ADP-ribosylated nucleosome substrate relative to the unmodified nucleosome substrate in the respective nuclear extract is shown. Data are represented as mean ± s.e.m. (n = 3). * indicates p-value < 0.02, obtained using an unpaired Student’s t-test with Welch’s correction.

To investigate the ability of nucleosome serine ADPr to stimulate ALC1 activity in a more physiological context, mammalian cell nuclear extracts were employed as a source of remodeling activity with the ADP-ribosylated nucleosome substrates. Nuclear extracts were prepared from wild-type or ALC1 knock-out (KO) HEK293T cells and the presence of various endogenous chromatin remodelers was confirmed (Fig. 6c and d). Each extract was then incubated with unmodified, H2BS6ADPr_1_, H2BS6ADPr_4_, or H2BS6/H3S10ADPr_4_ nucleosomes and remodeling activity was determined via the REA assay. The wild-type extract exhibited a ∼3-fold increase in remodeling activity towards the H2BS6ADPr_4_ and H2BS6/H3S10ADPr_4_ nucleosomes when compared to their unmodified counterpart (Fig. 6e). Contrastingly, there was no appreciable increase in activity towards the H2BS6ADPr_1_ nucleosome. Strikingly, the ALC1-KO nuclear extract exhibited similar remodeling activity towards all nucleosomes regardless of their ADPr status (Fig. 6e, and Supplementary Fig. 6c). We also note that no accumulation of additional ADPr events was detected in these lysates throughout the duration of the assay and only minor ADPr hydrolysis from the H2BS6/H3S10ADPr_4_ nucleosome was detected while other modified nucleosome substrates were unaffected (Supplementary Fig. 6d). These results suggest that nucleosome serine ADPr is sufficient to activate ALC1 in the nuclear milieu and that ALC1 is the primary chromatin remodeler responsible for directly manipulating the ADP-ribosylated nucleosomes described here.

## Discussion

Chemical and topological complexities have stymied previous efforts to synthesize poly-ADP-ribosylated proteins. Our investigation of HPF1-dependent and -independent PARP1 activities in peptide serine ADPr reactions guided the development of a multistep chemoenzymatic approach that is broadly applicable for the preparation of poly-ADP-ribosylated peptides and proteins. Through the use of chemically homogenous, ADP-ribosylated histones we were able to define a biochemical role for nucleosome serine ADPr and explore long-standing questions related to DNA damage-induced chromatin remodeling.

Multiple recent reports show that the PARP1/2:HPF1 complex catalyzes the formation of relatively short poly-ADP-ribose chains(Bilokapic et al., 2020; Bonfiglio et al., 2020; Gibbs-Seymour et al., 2016). Our study is unique in that we prepare unmodified and mono-ADP-ribosylated peptide substrates and use HPLC-MS to analyze PARP1 reaction products in the absence and presence of HPF1. This approach demonstrated that HPF1 simultaneously stimulates mono-ADPr activity and blocks ADP-ribose chain elongation on trans-peptide substrates. Our data support PARP2:HPF1 structural implications that mono- and poly-ADPr are mutually exclusive activities(Suskiewicz et al., 2020) and demonstrate that structural dynamics are insufficient to accommodate both catalytic mechanisms. Notably, HPF1 and PARP1/2 undergo DNA damage-induced ADPr, which may serve to disrupt the complex and initiate chain elongation from mono-ADP-ribosylated proteins. This would explain why we and others observe elongation activity in recombinant assays that include relatively high molar ratios of HPF1 to PARP1; ADPr on one or both complex components decreases the effective PARP1:HPF1 concentration as the reaction progresses. It is also likely that high HPF1 concentration is necessary to ensure rapid binding to the pre-formed PARP1:DNA complex(Suskiewicz et al., 2020) for immediate inhibition of elongation activity. Alternatively, we note that the cellular molar ratio of PARP1 to HPF1 (20:1)(Hein et al., 2015) is favorable for a mechanism wherein free PARP1 displaces the PARP1/2:HPF1 complex once mono-ADP-ribose seeding has occurred.

In chromatin remodeling experiments, ALC1 exhibits modest preference for the H2BS6 site and tetra-ADPr despite the observation that all H2BS6ADPr_3,4_ and H3S10ADPr_3,4_ peptides engage the ALC1 macrodomain with equal affinity. It is therefore likely that each histone modification site requires an ideal ADP-ribose chain length that allows the ATPase domain to progress through the DNA translocation cycle while the macrodomain:histone tail interaction is maintained. There are several factors that may explain why nucleosome serine ADPr more efficient than auto-ADP-ribosylated PARP1 for ALC1 activation in our assays: (i) robust ALC1 activation by auto-ADP-ribosylated PARP1 may require a specific modification site and ADP-ribose chain length that is only partially represented on our automodified PARP1 constructs, (ii) the PARP1:nucleosome interaction, while necessary for ALC1 recruitment and activation, may also sterically abrogate DNA translocation activity, and (iii) a direct interaction between ALC1 and ADP-ribosylated nucleosomes may be stronger than the ternary complex that is mediated by automodified PARP1, as evidenced from nucleosome pull-down efficiency in Fig. 5d.

Critical distinctions unique to nucleosome ADPr over other ADP-ribosylated proteins are: (i) the nucleosome-incorporated histones cannot diffuse away from the DNA damage site, and (ii) the stimulatory ADP-ribose chain is not tethered to a DNA-bound protein that may sterically hinder remodeling by ALC1. Therefore, nucleosome ADPr offers a fail-safe mechanism to ensure that robust ALC1-dependent remodeling can persist in the event that automodified PARP1 dissociates from the damage site prior to ALC1 activation. It is also interesting that ALC1 exhibits prolonged retention at DNA damage sites in HPF1-null cells where serine ADPr does not occur(Gibbs-Seymour et al., 2016). This is consistent with our observation that aspartate/glutamate-automodified PARP1 is the least potent activator in biochemical assays. It is plausible that serine ADPr, be it tethered to the nucleosome or PARP1, is critical for ALC1 remodeling activity at DNA damage sites in cells. While our technology has allowed us to separate and characterize ALC1 activation by ADPr on nucleosomes or PARP1 in a reconstituted environment, new approaches will be required to specifically control these parameters and analyze their contributions to ALC1-dependent remodeling at damage sites in cells.

Analyses of remodeling activity in biochemical assays and mammalian cell nuclear extracts show that nucleosome serine ADPr is sufficient to specifically activate ALC1 in the absence of auto-ADP-ribosylated PARP1. We surmise that other PARP1/2-dependent chromatin remodelers are recruited to damage sites via alternative ADPr modification sites or chain lengths, as has been reported for SMARCA5(Smeenk et al., 2013). Additionally, these remodelers may not directly interact with ADP-ribose but are rather recruited by alternative PARP1/2-dependent activities, a phenomenon that has been demonstrated for CHD4(Smith et al., 2018). Thus, our study supports the ‘PAR code’ hypothesis(Aberle et al., 2020) as it pertains to chromatin structure at DNA lesions wherein different ADPr sites and chain lengths may orchestrate spaciotemporal control over unique remodeler activities. Notably, dozens of proteins reportedly exhibit PARP1/2-dependent recruitment to DNA damage sites and have been annotated as ADP-ribose ‘readers’(Ray Chaudhuri and Nussenzweig, 2017; Teloni and Altmeyer, 2016). With full-length ADP-ribosylated proteins, ADPr-mediated activities can now be reconstituted for rigorous biochemical, biophysical, and structural analysis.

Beyond protein recruitment, it will now be possible to explore the direct biophysical effects that H2B and H3 ADPr have on poly-nucleosome array structure and compaction. Our modular chemoenzymatic approach can also be expanded to other PARP1/2:HPF1 substrate proteins, wherein one would expect to find ADPr exerts its effects via unique regulatory mechanisms that are tailored to the target protein. As demonstrated here, critical aspects of PARP biological function can be unveiled by reconstituting ADP-ribosylated proteins and related signaling pathway components. A greater understanding of PARP-regulated biological processes, including ALC1 activation, may lead to identification of new biomarkers and therapeutic strategies for PARP inhibitor-sensitive diseases.

### Technological limitations

The method described here is currently limited to installation of ADP-ribose units ∼4-5 linear units in length. Exceedingly large-scale reactions would be required to prepare peptides modified with longer ADP-ribose chains. Therefore, this method is ideal to study signal transduction events that are mediated by relatively short ADP-ribose chains. Our strategy also requires that a peptide of interest be a substrate for the PARP1:HPF1 complex. Alternative ADP-ribosyltransferases will be required to install ADPr on proteins that are not endogenous targets of this complex using the chemoenzymatic approach presented here. Regarding accessibility of modification sites that are not proximal to the protein amino-terminus: as proof of feasibility, the H3S28 peptide construct (amino acids 21-34) was prepared with an N-terminal thiazolidine and a C-terminal bis(2-sulfanylethyl)amido (SEA) group. This peptide can be easily activated for N-terminal (via SEA to thioester conversion) or C-terminal ligations (via thiazolidine to N-terminal cysteine conversion) and such a synthetic strategy will enable sequential protein ligations in the future. Lastly, our method is still susceptible to restraints that exist throughout the field of protein chemistry. This means that alternative protein ligation technologies will be required to install modification onto full-length proteins that are not amenable to protein folding.

## Acknowledgements

We thank Dr. W. Lee Kraus, Dr. Deepak Nijhawan, Dr. Steven McKnight, Dr. Benjamin Tu and members of the Liszczak laboratory for insightful discussions. We thank Dr. Andrew Lemoff and the UT Southwestern Proteomics Core for technical assistance. We also thank members of the laboratories of Dr. Ping Mu and Dr. Michael Roth for help with reagents. This work was supported by grants from the Welch Foundation (I-2039 to G.L.), the Cancer Prevention Research Institute of Texas (RR180051 to G.L.), and the American Cancer Society (UTSW-IRG-17-174-13). G.L. is the Virginia Murchison Linthicum Scholar in Medical Research.

## Author contributions

G.L., R.B., J.M., and K.T., conceived the study and designed experiments. G.L., R.B., J.M., J.S., and K.T. carried out molecular cloning, protein purification and characterization, peptide synthesis and protein ligation chemistries. G.L., J.M., R.B. and K.T. performed peptide interaction and ADPr assays. N.S.W., J.K. and J.M. performed LC-MS/MS analysis. J.M. and G.L. performed all chromatin remodeling and cellular experiments. All authors analyzed data. G.L. and J.M. wrote the manuscript and prepared figures with input from all authors.

## Quantification and statistical analysis

Details related to replicates, error, and curve fitting are described in respective figure legends. In Fig. 6e, the difference of means of two samples was statistically significant with p-value < 0.02, obtained using an unpaired Student’s t-test with Welch’s correction.

## Contact for reagent and resource sharing

Further information and requests for reagents may be directed to and will be fulfilled by the Lead Contact, Dr. Glen Liszczak.

## Data availability

Representative HPLC chromatograms, LC-MS characterizations and gel images are included in Supplementary Information. Complete raw data (in triplicate) for all quantitative experiments are included in the excel spreadsheets of Supplementary Dataset. Additional data will be provided upon request.

## METHODS

### Molecular cloning, protein expression, and protein purification

#### General protocols

All PCR amplification steps described here were performed using the Phusion High-Fidelity DNA Polymerase (NEB) according to the manufacturer’s protocols. All DNA oligonucleotides were synthesized by Sigma-Aldrich (Milwaukee, WI) or Integrated DNA Technologies (Coralville, IA). All plasmids used in this study were sequence verified by GENEWIZ (South Plainfield, NJ) or EurofinsGenomics (Louisville, KY). All cloning was carried out using Mach1 E. coli cells (ThermoFisher) and protein expression in E. coli was carried out in Rosetta2 cells (Sigma-Aldrich).

#### PARP1/PARP2 Expression and Purification

The full-length PARP1 gene was purchased from GE Healthcare and subcloned into a pACEBac1 plasmid bearing an N-terminal 6xHis-tag via a Gibson Assembly (NEB). The PARP2 expression plasmid (C-terminal FLAG-6xHis-tag) is available on Addgene (plasmid #: 111574). PARP1 and PARP2 proteins were produced in Sf9 cells (ThermoFisher) using a baculovirus expression system. Corresponding plasmids were transformed into DH10Bac cells (ThermoFisher) and bacmids were isolated via manufacturer’s protocols (ThermoFisher). All subsequent Sf9 cell and baculovirus manipulations were performed in a sterile biosafety cabinet. Cellfectin^TM^ II (ThermoFisher) was employed to transfect 10 μg of bacmid into 1×10^6^ attached Sf9 cells following manufacturer’s protocols (ThermoFisher). P1 virus was harvested 3 days post-transfection. 1 mL of P1 virus was then used to infect 20 mL of Sf9 cells grown in suspension at 1.5×10^6^ cells per mL, which were maintained in a dark orbital shaker at 27 °C. Cells were centrifuged and supernatant (P2 virus) was collected once cell viability dropped to 50%, as measured by trypan blue staining. P3 virus was generated by infecting 50 mL of Sf9 cells at 1.5×10^6^ cells per mL with 0.5 mL of P2 virus. P3 virus was harvested once cells reached 50% viability. Protein production was achieved by treating 2 L of Sf9 cells at 2.0×10^6^ cells per mL with 20 mL of P3 virus for 48 h.

For PARP1, cells were harvested by centrifugation and disrupted via sonication in a lysis buffer containing 50 mM Tris, pH 7.5, 1 M NaCl, 1 mM MgCl_2_, 5 mM beta-mercaptoethanol (β-ME), and protease inhibitor cocktail (Roche). Soluble lysate was isolated via centrifugation at 100,000 RCF for 60 minutes at 4 °C. The target protein was captured on Ni-NTA resin that was pre-equilibrated in lysis buffer. Following 1-hour batch binding, resin was washed with 50 column volumes (CV) of lysis buffer supplemented with 25 mM imidazole and eluted in a buffer containing 50 mM Tris, pH 7.0, 100 mM NaCl, 1.5 mM MgCl_2_, and 5 mM β-ME. Target protein was then loaded onto a HiTrap Heparin (GE Healthcare) column pre-equilibrated in a low salt buffer (50 mM Tris, pH 7.0, 150 mM NaCl, 1 mM EDTA, 1 mM TCEP) and elution was achieved via an isocratic salt gradient to a high salt buffer (50 mM Tris, pH 7.0, 1 M NaCl, 1 mM EDTA, 1 mM TCEP). Fractions containing the target protein were concentrated to 2 mL using an Amicon Ultra Centrifugal filter (Millipore; 30 kDa molecular weight cut-off (MWCO)) and injected into a gel filtration column (HiLoad 16/60 Superdex 200; GE Healthcare) that had been pre-equilibrated with a buffer containing 50 mM Tris, pH 7.5, 150 mM NaCl, 10% glycerol, and 1 mM TCEP. Pure fractions (as judged by SDS-PAGE) were pooled and concentrated to 100 μM, flash frozen in single-use aliquots, and stored at −80 °C.

For PARP2, cells were harvested by centrifugation and disrupted via sonication in a lysis buffer containing 20 mM Tris, pH 7.9, 500 mM NaCl, 4 mM MgCl_2_, 0.4mM EDTA, 20% glycerol, 2mM DTT, 0.4mM PMSF, and protease inhibitor cocktail (Roche). Soluble lysate was isolated via centrifugation at 100,000 RCF for 60 minutes at 4 °C. The supernatant was carefully removed without disturbing the top layer and an equal volume of dilution buffer containing 20 mM Tris, pH 7.9, 10% glycerol, 0.02% NP-40 and protease inhibitor cocktail (Roche) was added to it. The target protein was captured on anti-FLAG M2 magnetic resin that was pre-equilibrated with dilution buffer. Following a 60 min batch binding, resin was washed with 50 CV of wash buffer containing 20 mM Tris, pH 7.9, 150 mM NaCl, 2 mM MgCl_2_, 0.2 mM EDTA, 15% glycerol, 0.01% NP-40, 0.2 mM PMSF, 1 mM DTT and protease inhibitor cocktail (Roche), and eluted in the wash buffer supplemented with FLAG peptide at a concentration of 0.25 mg/mL. Pure protein was concentrated using an Amicon Ultra Centrifugal filter (Millipore; 30 kDa MWCO) to around 55 μM, as determined by BSA standards in SDS-PAGE, flash frozen in single-use aliquots, and stored at −80 °C.

#### HPF1 (and HPF1D283A mutant)

A pET30 plasmid harboring the 6xHis-SUMO-FLAG-HPF1 protein (addgene plasmid #: 111577), encoding amino acids 27-346, was transformed into Rosetta2 (DE3) cells and inoculated into 6 L of Luria Broth (Miller). Cells were grown in a shaker at 37 °C up to an OD_600_ of 0.6 and protein expression was induced with 0.5 mM IPTG at 18 °C for 16 h. Cells were harvested by centrifugation and disrupted via sonication in a lysis buffer containing 50 mM Tris, pH 7.5, 500 mM NaCl, 5 mM β-ME and 1 mM PMSF. Soluble lysate was isolated via centrifugation at 40,000 RCF for 40 minutes at 4 °C. Target protein was captured on Ni-NTA resin that was pre-equilibrated in lysis buffer. Following 1 h batch binding at 4 °C, resin was washed with 50 CV of lysis buffer supplemented with 25 mM imidazole and protein was eluted in lysis buffer supplemented with 300 mM imidazole. The elution was dialyzed into a buffer containing 50 mM Tris, pH 7.5, 200 mM NaCl, and 5 mM TCEP for 16 h at 4 °C in the presence of the Ulp1 protease to cleave the SUMO tag. The dialysate was then incubated with Ni-NTA resin pre-washed with the dialysis buffer for 1 h at 4 °C to capture the cleaved SUMO tag and the Ulp1, and the flow-through containing the target protein was collected. The flow-through was concentrated to 2 mL using an Amicon Ultra Centrifugal filter (Millipore; 30 kDa MWCO) and injected into a gel filtration column (HiLoad 16/60 Superdex 200) that had been pre-equilibrated with a buffer containing 50 mM Tris, pH 7.5, 200 mM NaCl, 10% glycerol, and 2 mM TCEP. Pure fractions (as judged by SDS-PAGE) were concentrated to around 600 μM, flash frozen in single-use aliquots, and stored at −80 °C. The HPF1D283A bacterial expression plasmid was generated via inverse PCR from the parent pET30 plasmid containing the HPF1 construct and transformed into Rosetta2 (DE3) cells. It was purified in the same way as described for HPF1.

#### ARH3

A pET30 plasmid harboring the 6xHis-SUMO-ARH3 protein (addgene plasmid #: 111578) was transformed into Rosetta2 (DE3) cells and inoculated into 6 L of Luria Broth (Miller). Protein expression was induced with 0.5 mM IPTG at a cell OD_600_ of 0.6. Expression was carried out at 18 °C for 16 h. Cells were harvested by centrifugation and protein was purified using Ni-NTA resin followed by reverse nickel and size-exclusion chromatography (SEC) in a manner similar to that described for HPF1. Pure fractions from the SEC (as judged by SDS-PAGE) were concentrated to around 600 μM, flash frozen in single-use aliquots, and stored at −80 °C.

#### PARG

A PARG gene fragment encoding amino acids 448-976 was synthesized by Integrated DNA Technologies and cloned into a modified pET30 vector via Gibson Assembly to produce an E. coli expression plasmid for the 6xHis-SUMO-PARG construct. The plasmid was transformed into Rosetta2 (DE3) cells and inoculated into 2 L of Luria Broth (Miller). Protein expression was induced with 0.5 mM IPTG at a cell OD_600_ of 0.6, and carried out at 18 °C for 16 h. Cells were harvested by centrifugation and protein was purified using Ni-NTA resin followed by reverse nickel and size-exclusion chromatography (SEC) in a manner similar to that described for HPF1. Pure fractions from the SEC (as judged by SDS-PAGE) were concentrated to around 300 μM, flash frozen in single-use aliquots, and stored at −80 °C.

#### ALC1 macrodomain

The full-length ALC1 gene was synthesized by Twist Biosciences. A fragment encoding amino acids 636-878, corresponding to the macrodomain(Singh et al., 2017), was cloned into a modified pET30 vector via Gibson Assembly to produce an E. coli expression plasmid for the 6xHis-SUMO-ALC1macrodomain construct. The plasmid was transformed into Rosetta2 (DE3) cells and inoculated into 6 L of Luria Broth (Miller). Protein expression was induced with 0.5 mM IPTG at a cell OD_600_ of 0.6. Expression was carried out at 18 °C for 16 h. Cells were harvested by centrifugation and disrupted via sonication in a lysis buffer containing 50 mM Tris, pH 7.5, 500 mM NaCl, 5 mM β-ME and 1 mM PMSF. Soluble lysate was isolated via centrifugation at 40,000 RCF for 30 minutes at 4 °C. Target protein was captured on Ni-NTA resin that was pre-equilibrated in lysis buffer. Following 1 h batch binding, resin was washed with 50 CV of lysis buffer supplemented with 25mM imidazole, and then 2 CV of lysis buffer supplemented with 80 mM imidazole, and target protein was eluted in lysis buffer supplemented with 300 mM imidazole. The elution was dialyzed into a buffer containing 50 mM Tris, pH 7.5, 200 mM NaCl, and 5 mM TCEP for 16 h at 4 °C in the presence of Ulp1 to cleave the SUMO tag. The dialysate was then incubated with Ni-NTA resin pre-washed with the dialysis buffer for 1 h at 4 °C to capture the cleaved SUMO tag and the Ulp1, and the flow-through containing the target protein was collected. The flow-through was concentrated to 2 mL using an Amicon Ultra Centrifugal filter (Millipore; 30 kDa MWCO) and injected into a gel filtration column (HiLoad 16/60 Superdex 200) that had been pre-equilibrated with a buffer containing 40 mM Tris, pH 7.5, 200 mM NaCl, 10% glycerol, and 2 mM TCEP. Pure fractions (as judged by SDS-PAGE) were concentrated to around 400 μM, flash frozen in single-use aliquots, and stored at −80 °C.

#### ALC1

The full-length ALC1 gene was cloned into a modified pACEBac1 vector via Gibson Assembly to produce the 6xHis-1xFLAG-ALC1 DNA construct. Bacmid and baculovirus preparation was performed as described for PARP1/2. Protein expression was achieved by treating 2 L of Sf9 cells at 2.0×10^6^ cells per mL with 20 mL of P3 virus for 48 h. Cells were harvested by centrifugation and target protein was purified using anti-FLAG M2 magnetic resin in a procedure similar to that described for PARP2. Pure protein was concentrated using an Amicon Ultra Centrifugal filter (Millipore; 30 kDa MWCO) to around 20 μM, as determined by BSA standards in SDS-PAGE, flash frozen in single-use aliquots, and stored at −80 °C.

#### CHD4

The full-length CHD4 gene was purchased from Horizon Discovery and cloned into a modified pACEBac1 vector via Gibson Assembly to produce the CHD4-1xFLAG DNA construct. Bacmid and baculovirus preparation was performed as described for PARP1/2. Protein production was achieved by treating 2 L of Sf9 cells at 2.0×10^6^ cells per mL with 20 mL of P3 virus for 48 h. Cells were harvested by centrifugation and target protein was purified using anti-FLAG M2 magnetic resin in a procedure similar to that described for PARP2. Pure protein was concentrated using an Amicon Ultra Centrifugal filter (Millipore; 30 kDa MWCO) to around 20 μM, as determined by BSA standards in SDS-PAGE, flash frozen in single-use aliquots, and stored at −80 °C.

#### Auto-ADP-ribosylated PARP1 (serine-linked)

The PARP1 purified by the above method was incubated in auto-ADP-ribosylation reactions with NAD^+^ and activating DNA. An HPF1 was titration experiment was employed to identify two concentrations at which relatively short or long serine-linked ADP-ribose chains could be installed on PARP1. Reactions (5 mL) included 2 μM of purified recombinant PARP1, 5 μM of activating DNA, and 250 μM of NAD^+^ and were incubated in a buffer containing 50 mM Tris (pH 7.5), 20 mM NaCl, 2 mM MgCl_2_, 1 mM TCEP with either 5 μM or 25 μM HPF1 for 30 min at 30 °C. The reaction with 5 μM HPF1 yielded PARP1 automodified with long serine-linked ADP-ribose chains and that with 25 μM HPF1 yielded PARP1 automodified with short serine-linked ADP-ribose chains. After completion of the automodification reaction, the sample was injected onto a 5 mL Cytiva HiTrap Heparin column (GE) that was pre-equilibrated with low salt buffer (150 mM NaCl, 50 mM Tris pH 7.5, 2 mM βMe, 1 mM MgCl_2_). The column was washed with 5 CV of low salt buffer and the protein was eluted using a gradient from the low salt buffer to a high salt buffer (1 M NaCl, 50 mM Tris pH 7.5, 2 mM βMe, 1 mM MgCl_2_) over 20 CV at a flow rate of 3 mL/min. Fractions were analysed on SDS-PAGE and those containing pure protein were pooled, concentrated to around 20 μM, supplemented with 10% glycerol, flash-frozen into single-use aliquots and stored in −80 °C.

#### Core histones (H2A, H2B, H3, H4)

Identical purification protocols were employed for each full-length histone. Expression plasmids were transformed into Rosetta2 (DE3) cells and inoculated into 1 L of Luria Broth (Miller). Protein expression was induced with 0.5 mM IPTG at a cell OD_600_ of 0.6. Expression was carried out at 37 °C for 3 h. Cells were harvested by centrifugation and disrupted via sonication in a lysis buffer containing 40 mM Tris, pH 7.5, 0.3 M NaCl, 1 mM EDTA, 5 mM β-ME, and 1 mM PMSF. Following centrifugation at 20,000 RCF for 30 minutes at 4 °C, the inclusion body pellet was then washed with lysis buffer supplemented with 1% Triton X-100 and centrifuged at 20,000 RCF for 15 min. This wash was repeated two more times with the final wash being performed in the absence of Triton X-100. Next, recombinant histone protein was extracted from the insoluble pellet in a buffer containing 50 mM Tris, 7.5, 300 mM NaCl, 6 M guanidine hydrochloride, and 5 mM β-ME for 1 hour at 25 °C and centrifuged at 20,000 RCF for 30 min. The soluble extract was then centrifuged at 100,000 RCF, injected onto a preparative C18 RP-HPLC column equilibrated in Solvent A (0.1% TFA in water) and eluted via an isocratic gradient 20–80% Solvent B (90% acetonitrile, 0.1% TFA in water) over a period of 30 min. Pure fractions (as determined by LC–MS) were lyophilized and stored at −80 °C until use in histone octamer assembly.

#### H2B (amino acids 17-125) and H3 (amino acids 21-135)

Identical protocols were employed for each truncated 6xHis-ketosteroid isomerase-SUMO-tagged histone. The ketosteroid isomerase tag (synthesized by IDT and incorporated into histone expression plasmids via Gibson Assembly) rapidly shuttles truncated histones to E. coli inclusion bodies to protect them from degradation and increase yield. Truncated histones were expressed and extracted as described for full-length histone constructs. Following extraction, the histones were immobilized on Ni-affinity resin in extraction buffer, washed with 50 mM Tris, pH 7.5, 300 mM NaCl, 6 M guanidine hydrochloride, 20 mM imidazole, and 5 mM β-ME, and eluted in wash buffer supplemented with 300 mM imidazole. The eluted protein was dialyzed for 16 h at 4 °C into dialysis buffer (50 mM Tris, pH 7.5, 300 mM NaCl, 6 M urea, and 5 mM β-ME). Following dialysis, the sample was diluted three-fold with dilution buffer (50 mM Tris, pH 7.5, 300 mM NaCl, and 5 mM β-ME) in the presence of Ulp1 to cleave the ketosteroid isomerase-SUMO tag. This target proteins were then purified via preparative RP-HPLC and stored as described for full-length histone constructs.

#### 601 DNA preparation

The 200 bp template used to assemble all nucleosomes is shown below with the 601 sequence in bold, the PstI site in yellow, and the overhangs underlined: 5’-<u>GGCCGCTCTAGAACTAGTGGATCCGATATCGCTGTTCACCGCGTG</u>**ACAGGATGTATATAT CTGACACGTGCCTGGAGACTAGGGAGTAATCCCCTTGGCGGTTAAAACGCGGGGGACAGCGCGTACGTGCGTTTAAGCGGTGCTAGAGCTGTCTACGACCAATTGAGCGGCTGCAGCACCG GGATTCTCCAG**<u>CATCAGAG</u>-3’

The 601 sequence was purchased from IDT and incorporated into a pET30a plasmid via Gibson Assembly. DNA was amplified from the parent plasmid using Phusion polymerase and the primers shown in the Supplementary Table 4. The PCR product was purified using QIAquick Spin Columns (Qiagen) following manufacturer’s protocols. Following elution, an ethanol precipitation step was performed and DNA was resuspended to 1 μg/μL in water for use in nucleosome assembly.

To insert unique 5¢ primer-binding sites for the nucleosome competition remodeling assays, primers bearing unique 5¢ 15 bp overhangs were employed in the protocol described above. Primer sequences are shown in the Supplementary Table 4. The final template design is outlined below with the 601 sequence in bold, the PstI site in yellow, the unique 5¢ primer-binding site in teal, and the universal 3¢ primer-binding site in gray:

5’-nnnnnnnnnnnnnnnAGTGGATCCGATATCGCTGTTCACCGCGTG**ACAGGATGTATATATCTG ACACGTGCCTGGAGACTAGGGAGTAATCCCCTTGGCGGTTAAAACGCGGGGGACAGCGCGTACGTGCGTTTAAGCGGTGCTAGAGCTGTCTACGACCAATTGAGCGGCTGCAGCACCGGGA TTCTCCAG**CATCAGAG-3’

### Peptide synthesis

#### General protocols

All fluorenylmethyloxycarbonyl (Fmoc)-protected amino acids were purchased from Oakwood Chemical or Combi-Blocks. Peptide synthesis resins (Trityl-OH ChemMatrix and Rink Amide ChemMatrix) were purchased from Biotage. All analytical reversed-phase HPLC (RP-HPLC) was performed on an Agilent 1260 series instrument equipped with a quaternary pump and an XBridge Peptide C18 column (5 μm, 4 × 150 mm; Waters) at a flow rate of 1 mL/min. Similarly, semi-preparative scale purifications were performed employing a XBridge Peptide C18 semi-preparative column (5 μm, 10 mm × 250 mm, Waters) at a flow rate of 4 mL/min. Preparative RP-HPLC was performed on an Agilent 1260 series instrument equipped with a preparatory pump and a XBridge Peptide C18 preparatory column (10 µM; 19 × 250 mm, Waters) at a flow rate of 20 mL/min. All instruments were equipped with a variable wavelength UV-detector. All RP–HPLC steps were performed using 0.1% (trifluoroacetic acid, TFA, Oakwood Chemical) in H_2_O (Solvent A) and 90% acetonitrile (Sigma-Aldrich), 0.1% TFA in H_2_O (Solvent B) as mobile phases. For LC/MS analysis, 0.1% formic acid (Sigma-Aldrich) was substituted for TFA in mobile phases. Gradients and run times are described in the characterization section for each molecule. Mass analysis was carried out for each product on an LC/MSD (Agilent Technologies) equipped with a 300SB-C18 column (3.5 µM; 4.6 × 100 mm, Agilent Technologies) or a X500B QTOF (Sciex).

#### Preparation of amidated peptides

Sequence of H3 (1-20)-CONH_2_: **ARTKQTARKSTGGKAPRKQL**-CONH_2_

Sequence of H3S10A (1-20)-CONH_2_: **ARTKQTARKATGGKAPRKQL**-CONH_2_

Sequence of H2B (1-16)-CONH_2_: **PEPAKSAPAPKKGSKK**-CONH_2_

Sequence of H2BS6A (1-16)-CONH_2_: **PEPAKAAPAPKKGSKK**-CONH_2_

Sequence of PARP1 (501-515)-CONH_2_: **AALSKKSKGQVKEEG**-CONH_2_

Sequence of PARP1S507A (501-515)-CONH_2_: **AALSKKAKGQVKEEG**-CONH_2_

Sequence of TMA16 (2-19)-CONH_2_: **PKAPKGKSAGREKKVIHP**-CONH_2_

Sequence of TMA16S9A (2-19)-CONH_2_: **PKAPKGKAAGREKKVIHP**-CONH_2_

The above amidated peptides were synthesized via solid-phase peptide synthesis on a CEM Discover Microwave Peptide Synthesizer (Matthews, NC) using the Fmoc-protection strategy on Rink Amide-ChemMatrix resin (0.5 mmol/g). For coupling reactions, amino acids (5 eq) were activated with N,N′-diisopropylcarbodiimide (DIC, 5 eq, Oakwood Chemical)/Oxyma (5 eq, Oakwood Chemical) and heated to 90 °C for 2 min while bubbling with nitrogen gas in N,N-dimethylformamide (DMF, Oakwood Chemical). Fmoc deprotection was carried out with 20% piperidine (Sigma-Aldrich) in DMF supplemented with 0.1 M 1-hydroxybenzotriazole hydrate (HOBt, Oakwood Chemical) at 90 °C for 1 minute while bubbling with nitrogen gas. The H3 Cleavage from the resin was performed with 92.5% TFA, 2.5% triisopropylsilane (TIS, Sigma-Aldrich), 2.5% 1,2-ethanedithiol (EDT, Sigma-Aldrich), and 2.5% H_2_O for 2 h at 25 °C. The crude peptide was then precipitated by the addition of a 10-fold volume of cold ether and centrifuged at 4,000 RCF for 10 min at 4 °C. The pellet was resuspended in Solvent A and purified via preparative RP-HPLC using a linear gradient from 0-30% Solvent B over 30 minutes. Fractions were analyzed on analytical RP-HPLC and ESI-MS and those containing pure product (>95%) were pooled, lyophilized, and stored at −80 °C.

#### Fluorescein-labeled H3.1 (1-20)-CONH_2_, H2B (1-16)-CONH_2_

Peptides were synthesized as described for the amidated species. Prior to cleavage, 5(6)-carboxyfluorescein (3 eq, Sigma-Aldrich) was activated with PyAOP (3 eq, Oakwood Chemical) and N,N-diisopropylethylamine (DIPEA, 6 eq, Sigma-Aldrich) and coupled to the deprotected α-amine on resin for 30 minutes at 25 °C in DMF while bubbling with nitrogen gas. Resin was washed with DMF and treated with 20% piperidine in DMF prior to cleavage with 92.5% TFA, 2.5% TIS, 2.5% EDT, and 2.5% H_2_O for 2 h at 25 °C. The crude peptide was then precipitated by the addition of a 10-fold volume of cold ether and centrifuged at 4,000 RCF for 10 min at 4 °C. The pellet was resuspended in Solvent A and purified via preparative RP-HPLC using a linear gradient from 0-50% Solvent B over 40 minutes. Fractions were analyzed on analytical RP-HPLC and ESI-MS and those containing pure product (>95%) were pooled, lyophilized, and stored at −80 °C.

#### Synthesis of H3.1 (1-20) -NHNH_2_, H2B (1-16) - NHNH_2_

Sequence of H3.1 (1-20) -NHNH_2_: **ARTKQTARKSTGGKAPRKQL-NHNH_2_**

Sequence of H2B (1-16) -NHNH_2_: **PEPAKSAPAPKKGSKK-NHNH_2_**

H3 (1-20) and H2B (1-16) containing C-terminal hydrazide were synthesized similarly to the amidated peptides described above with the following modifications. ChemMatrix Trityl-OH PEG resin (0.49 mmol/g) was washed with dichloromethane (DCM, Oakwood Chemical) and reacted with 5% (v/v) thionyl chloride (Sigma-Aldrich) in DCM for 90 minutes at 25 °C. Resin was washed with DCM and this step was repeated to ensure efficient resin chlorination. Next, the resin was washed with DCM, DMF, and 5% (v/v) DIPEA in DMF. The resin was reacted with 9-fluorenylmethyl carbazate (Combi-Blocks) in the presence of DIPEA (20 eq) in DMF for 2 hr at RT. The resin was washed with DMF and the 9-fluorenylmethyl carbazate coupling step was repeated to ensure complete loading. The resin was washed with DMF and 5% (v/v) anhydrous methanol (Sigma-Aldrich) in DMF. For coupling reactions, amino acids (5 eq) were activated with DIC (5 eq) and Oxyma (5 eq) and heated to 50 °C for 10 min while bubbling with nitrogen gas in DMF. Fmoc deprotection was carried out with 20% piperidine in DMF supplemented with 0.1 M HOBt at 60 °C for 4 minutes while bubbling with nitrogen gas. Cleavage and purification were performed as described for amidated peptides.

For peptide thioesterification, purified peptides containing C-terminal hydrazide were dissolved in a de-gassed buffer of 6 M guanidine hydrochloride and 0.1 M sodium phosphate, pH 3.0. The reaction was initiated by adding sodium nitrite (15 eq, Sigma-Aldrich) at −15 °C 10 minutes. The pH was monitored and maintained at 3.0 throughout the reaction. Immediately following this reaction, MESNa (75 eq, Sigma-Aldrich) and TCEP (final concentration of 20 mM, GoldBio) were added and the pH was adjusted to 7.0. The mixture was incubated at 25 °C for additional 30 min and monitored by RP-HPLC and ESI-MS analyses. Once quantitative conversion was complete, the peptide was purified via preparative RP-HPLC with a linear gradient of 0-30% Solvent B over 30 min. Pure fractions were characterized as described for amidated peptides, pooled, lyophilized, and stored at −80 °C.

#### Synthesis of H3 (21-34)-SEA, H3 (21-34, S28A)-SEA

Sequence of H3 (21-34)-SEA: Thz-**TKAARKSAPATGG-SEA**

Sequence of H3S28A (21-34)-SEA: Thz-**TKAARKAAPATGG-SEA**

where,

Thz = thiazolidine, and SEA = bis(2-sulfanylethyl)amido group

H3 (amino acids 21-34) and the corresponding S28A mutant peptides containing N-terminal thiazolidine and C-terminal SEA were synthesized similarly to the amidated peptides described above with the following modifications. SEA resin (0.16 mmol/g; Iris Biotech) was weighed out, washed with DMF and bubbled in nitrogen for 15 min to swell the resin. Fmoc-glycine (5 eq) and HATU (1-Bis(dimethylamino)methylene-1H-1,2,3-triazolo [4,5-b]pyridinium 3-oxide hexafluorophosphate; 5 eq), and DIPEA (15 eq) were mixed in DMF and the resin was bubbled in this mixture for 1 h. This step was repeated with fresh reagents to ensure complete loading. The resin was then washed with DMF and bubbled in acetic anhydride:DIPEA:DMF (10:5:85) for 20 min for acetyl capping. For coupling reactions, amino acids (5 eq) were activated with DIC (5 eq) and Oxyma (5 eq) and heated to 50 °C for 10 min while bubbling with nitrogen gas in DMF. Fmoc deprotection was carried out with 20% piperidine in DMF supplemented with 0.1 M HOBt at 60 °C for 4 minutes while bubbling with nitrogen gas. Cleavage and purification were performed as described for amidated peptides. The use of thiazolidine offers a way to keep the thiol of the N-terminal cysteine protected while performing native chemical ligation on the C-terminus of the peptide.

### Recombinant PARP1:HPF1 complex ADPr activity assays and analysis

#### General protocols

To analyze PARP1:HPF1 ADPr activity on synthetic peptide substrates, 1 μM PARP1 (or 2 μM for H2B peptides), 10 μM HPF1, 2 mM NAD^+^ (Sigma-Aldrich), and 1 μM stimulating DNA (or 2 μM for H2B peptides; see Supplementary Table 4 for stimulating DNA sequence information) were combined into the ADPr reaction buffer (50 mM Tris, pH 7.5, 20 mM NaCl, 2 mM MgCl_2_, 5 mM TCEP) at a final volume of 25 μL (or 50 μL for H2B peptides). All substrate peptides were initially analyzed at a concentration of 180 μM (40 μM for H2B peptides). The reaction was then incubated at 30 °C for 25 min and quenched via addition of Solvent A to a final volume of 120 μL. Reactions were then centrifuged at 20,000 RCF for 5 min and 100 μL of the supernatant was injected onto an analytical C18 column for product analysis via RP-HPLC. An elution gradient of 0-35% Solvent B over 20 min was employed to separate the poly-ADP-ribosylated peptide products. Individual peaks corresponding to products with mono-, di-, tri-, tetra-, or penta-ADP-ribose were collected and analyzed by ESI-MS.

#### Fluorescent peptide ADPr

Reaction volumes were scaled to 1 mL to obtain sufficient amounts of each purified product for fluorescence polarization assays. For purification via semi-preparative RP-HPLC, an elution gradient of 5-20% Solvent B over 40 min was employed to optimize separation of the poly-ADP-ribosylated peptide products. Peaks corresponding to products with mono-, di-, tri-, tetra-, or penta-ADP-ribose were collected separately, analyzed by ESI-MS, and the pure fractions were pooled, lyophilized, and stored at −80 °C. The reaction and purified peptides were kept wrapped in aluminum foil whenever possible. We note that 5(6)-carboxyfluorescein causes peak splitting in HPLC characterization corresponding to individual fluorescein isomers. This phenomenon was unique to fluorescein-labeled peptides and peak resolution varied based on ADP-ribose chain length.

#### HPF1 titration analysis on unmodified peptide substrates

For the HPF1 titration experiments described in Fig. 1e, reactions were performed as described above in the presence of 0, 5, 10, 20, 50 or 100 μM HPF1. Histone peptide starting material was quantified via integration of the corresponding HPLC peak at A_214_. Peak area was converted to peptide concentration via a standardization curve that was generated using known quantities of substrate peptide. ADP-ribosylated peptide products were quantified via integration of corresponding HPLC peaks at A_280_. Peak areas were then converted to peptide concentrations via a standardization curve that was generated using known quantities of ADP-ribosylated peptides. Standardization curves were generated for the mono- and di-ADP-ribosylated products. We note that the peptide HPLC A_280_ signal is dependent upon the ADP-ribose moiety and no A_280_ signal is present for any unmodified peptides used in this study. Therefore, there is a linear relationship between product extinction coefficient at 280 nm and the number of ADP-ribose units that are attached to the peptide. This linear increase in extinction coefficient was extrapolated to quantify all products with chain lengths greater than or equal to di-ADP-ribose. All reactions and standardization curve samples were run on the same C18 column and HPLC instrument using identical mobile phase gradients. All reactions were performed in triplicate and error bars represent standard deviations.

The following formula was used to calculate percent conversion to each product in a given reaction:

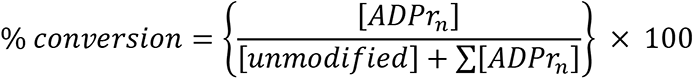

In the above formula:

[ADPr_n_] represents the concentration of an individual product modified with mono-, di-, tri-, tetra-, or penta-ADP-ribose

[unmodified] represents the concentration of the unmodified peptide starting material

Σ[ADPr_n_] represents the sum total concentration of all detectable ADP-ribosylated products

#### Optimized peptide mono-ADPr preparation

Optimal yield of mono-ADP-ribosylated peptides was achieved via a reaction of 1 μM PARP1 (or 2 μM PARP1 for H2B peptides), 20 μM HPF1, 5 μM of PARG, 10 mM NAD^+^, 0.5 mM unmodified substrate peptide, and 3 μM stimulating DNA (or 6 μM for H2B peptides) in ADPr reaction buffer. Reactions were incubated at 30 °C for 30 min and quenched via addition of 6 M guanidine hydrochloride and 0.1 M sodium phosphate. Purification was carried out on a preparative RP-HPLC C18 column and characterization by ESI-MS and glycohydrolase treatment was performed. We have scaled to as high as 15 mL reaction volume and 2 mM substrate peptide. Percent conversion of peptide starting material drop precipitously at higher substrate peptide concentrations. Importantly, 5 μM of PARG is included throughout this reaction to cleave all poly-ADP-ribosylated products back to mono-ADP-ribose. We also noticed that PARG enhances percent conversion at higher peptide concentrations and is necessary for quantitative conversion under the conditions described here. We suspect this is because PARG reverses PARP1 auto-poly-ADPr that accumulates throughout the reaction. Notably, auto-ADPr abrogates the PARP1:DNA interaction and inactivates the enzyme(Kim et al., 2004). For mono-ADPr of 0.5 mM of PARP1 (501-515) or TMA16 (2-19) peptide, 2 μM PARP1, 20 μM HPF1, 8 μM activating DNA, 2 μM PARG, and 10 mM NAD^+^ was used. For mono-ADPr of 0.5 mM of H3 (21-34) SEA peptide, 2.5 μM PARP1, 25 μM HPF1, 10 μM activating DNA, 2 μM PARG, and 10 mM NAD^+^ was used.

### Recombinant PARP1/2 ADPr polymerization activity assays and analysis

#### General protocols

To analyze PARP1 and PARP2 ADPr activity on peptide substrates, 2 mM NAD^+^ and 1 μM stimulating DNA were combined into the ADPr reaction buffer in the presence of 0.2, 1, or 5 μM PARP1 or PARP2, in a 25 μL reaction (2 μM PARP1/2, 2 μM DNA and 50 μL reaction volume for H2B peptides). All unmodified and mono-ADP-ribosylated substrate peptides were analyzed at a concentration of 180 μM (40 μM for H2B peptides). The reaction was incubated at 30 °C for 25 min, quenched via addition of 95 μL (70 μL in case of H2B peptide reactions) of Solvent A. It was then centrifuged at 20,000 RCF for 5 min and 100 μL of the supernatant was injected into an analytical C18 column for product analysis via RP-HPLC. An elution gradient of 0-35% Solvent B over 20 min was employed to optimize separation of the poly-ADP-ribosylated peptide products. Percent substrate turnover was calculated by integrating the peaks for the starting material and each product on the RP-HPLC A_280_ trace, normalizing them depending on their number of ADP-ribose moieties, and calculating ratio of total product to total peptide amounts for each reaction. All reactions were performed in triplicate and error bars represent standard deviations. Individual peaks corresponding to products with unique ADP-ribose chain lengths were collected and analyzed by ESI-MS.

#### HPF1 titration analysis on mono-ADP-ribosylated peptide substrates

To analyze HPF1-dependent inhibition of PARP1/2 elongation activity, elongation reactions were performed as described above in the presence of 0, 5, 10, 20, 50 or 100 μM of HPF1 or HPF1D283A, 1 μM of PARP1 or PARP2, and 180 μM of mono-ADP-ribosylated H3 peptide. Percent conversion of the mono-ADP-ribosylated peptide substrates to poly-ADP-ribosylated peptide products was calculated by integrating the peaks for the starting material and each product on the RP-HPLC A_280_ trace. Peak areas were again converted to molar concentrations (as described in HPF1 titration analysis on unmodified peptide substrates).

The following formula was used to calculate fraction elongated in a given reaction:

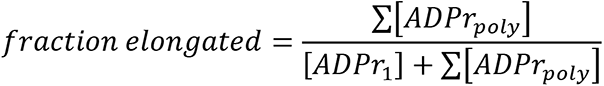

In the above formula:

[ADPr_1_] represents the concentration of the mono-ADP-ribosylated peptide starting material

Σ[ADPr_poly_] represents the sum total concentration of all detectable poly-ADP-ribosylated peptide products (products modified with di-, tri-, tetra-, or penta-ADP-ribose*)

*in Fig. 2b and c, the data is represented as the sum total of all poly-ADP-ribosylated peptide products. For distribution of product species, see Supplementary Fig. 2a, b and e. The relative fraction elongated at any concentration of HPF1 was calculated as:

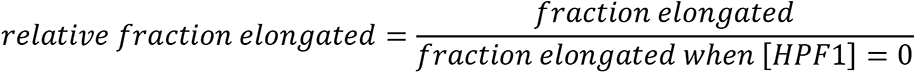

#### Optimized peptide poly-ADPr preparation

When optimal yield of poly-ADP-ribosylated peptides is desired, a reaction of 1 μM PARP1 (2 μM PARP1 for H2B peptides), 10 mM NAD^+^, 500 μM mono-ADP-ribosylated substrate peptide, and 3 μM stimulating DNA (6 μM for H2B peptides) in ADPr reaction buffer is employed. Reactions are incubated at 30 °C for 30 min and quenched via addition of 6 M guanidine hydrochloride and 0.1 M sodium phosphate. Purification is carried out on a preparative RP-HPLC C18 column and characterization by ESI-MS and glycohydrolase treatment is performed. We have scaled to as high as 14 mL reaction volume and 1 mM mono-ADP-ribosylated substrate peptide. For poly-ADPr of 0.5 mM of mono-ADP-ribosylated PARP1 (501-515) peptide, 5 μM PARP1, 25 μM activating DNA, and 10 mM NAD^+^ was used. For poly-ADPr of TMA16 (2-19) peptide, 2 μM PARP1, 8 μM activating DNA, and 10 mM NAD^+^ was used. For poly-ADPr of 0.5 mM of H3 (21-34) SEA peptide, 5 μM PARP1, 40 μM activating DNA, and 10 mM NAD^+^ was used.

### Glycohydrolase activity assays

For histone H3 peptide analysis, ADPr reactions containing 1 μM PARP1, 10 μM HPF1, 2 mM NAD^+^, and 1 μM stimulating DNA, and 125 μM unmodified H3 peptide were combined into the ADPr reaction buffer at a final volume of 75 μL. Following a 25 min incubation at 30 °C, the reaction was quenched with 10 μM Olaparib (Selleckchem) and 25 μL was removed for pre-glycohydrolase treatment analysis. ARH3 or PARG was then added to a final concentration of 3 μM or 1 μM, respectively, and incubated at 37 °C for 2 h. Pre- and post-glycohydrolase-treated samples were then analyzed via analytical RP-HPLC on a C18 column using an elution gradient of 0-35% Solvent B over 20 min. Product identities were verified by ESI-MS.

We note that for H2B, glycohydrolase analysis was performed after the PARP1 elongation reaction from the mono-ADP-ribosylated peptide. This is because only very low levels of poly-ADP-ribosylated products could be generated in the PARP1:HPF1 reaction. Elongation was much more efficient from mono-ADP-ribosylated H2B peptides in reactions that lacked HPF1. For H2B product glycohydrolase analysis, peptide ADPr reactions containing 2 μM PARP1, 2 mM NAD^+^, 180 μM mono ADP-ribosylated substrate peptide, and 2 μM stimulating DNA were combined into the ADPr reaction buffer at a final volume of 150 μL. Following a 25 min incubation at 30 °C, the reaction was quenched with 10 μM Olaparib and 50 μL was removed for pre-glycohydrolase treatment analysis. ARH3 or PARG was then added to a final concentration of 3 μM or 1 μM, respectively, to 50 μL of the elongated reaction and incubated at 37 °C for 2 h. Pre- and post-glycohydrolase-treated samples were then analyzed via analytical RP-HPLC on a C18 column using an elution gradient of 0-35% Solvent B over 20 min. Product identities were verified by ESI-MS.

### LC-MS/MS analysis of PAR chains on a peptide substrate

To analyze branching of PAR chains installed on peptide substrates using our technology, we utilized the LC-MS/MS-based approach outlined by Chen, et al(Chen et al., 2018). PAR chains from a purified tetra-ADP-ribosylated H2B (1-16) peptide were subjected to treatment with Alkaline Phosphatase (Sigma-Aldrich) and Phosphodiesterase I (Sigma-Aldrich). ADP-ribosylated peptide (80 μM) was incubated with ∼8 units of Phosphodiesterase I and ∼300 units of alkaline phosphatase at 30 °C overnight in a 0.5 mL reaction in a buffer containing 50 mM Tris (pH 7.5), 20 mM NaCl, 2 mM MgCl_2_, and 5 mM TCEP. The reaction products were then desalted and deproteinized via RP-HPLC (C18 column) and lyophilized. The lyophilized powder containing a mixture of the digestion products was then resuspended in water to a final concentration of 20 μg/mL and 12 μL was injected onto Phenomenex Synergi Polar-RP column (150 x 2 mm, 4 μm packing) and analyzed by LC-MS/MS using a Sciex QTRAP® 6500+ mass spectrometer coupled to a Shimadzu Nexera X2 UPLC. The chromatographic conditions were as follows: Solvent A: dH_2_0 + 0.2% acetic acid, Solvent B: acetonitrile + 0.2% acetic acid; flow rate: 0.46 mL/min; 0-2 min 1% B, 2-2.5 min gradient to 78% B, 2.5-3 min 78% B, 3-3.1 min gradient to 80% B, 3.1-3.5 min 80% B, 3.5-3.6 min gradient to 95% B, 3.6-6 min 95% B, 6.5 min gradient to 1% B, 6.5-7.5 min 1% B. Analytes were detected with the mass spectrometer in MRM (multiple reaction monitoring) mode by following the precursor to fragment ion transitions as follows: Adenosine 268 → 136 (2.27 min retention time), ribosyl-adenosine 400 → 268 and 136 (3.08 min retention time), diribosyl-adenosine 532.18 → 400, 268 and 136 (3.5 min retention time). Peaks were integrated and peak areas were determined by AB Sciex Analyst 1.71 with HotFix 1 software.

### PARP1 pull-down assays

#### Immunoprecipitations with PARP1 and HPF1

A 100 μL solution of 5 μM FLAG-HPF1 (or HPF1D283A), 1 μM PARP1, 10 μM Olaparib, and 10 μM stimulating DNA in Pull-Down Buffer (50 mM Tris, pH 7.5, 50 mM NaCl, 2 mM MgCl_2_, 0.1% Triton X-100, 1 mM DTT) was incubated for 25 min at 25 °C. This solution was then centrifuged at 20,000 RCF for 10 minutes and the supernatant was added to 10 μL of Anti-FLAG M2 magnetic resin (MilliporeSigma; pre-equilibrated in Pull-Down Buffer), after keeping aside 30 μL from the reaction as an input control for SDS-PAGE gel analysis. Resin was incubated on an end-over-end rotator at 4 °C for 30 min, washed for 3 times for 1 min each with 0.5 mL of Pull-Down Buffer, and eluted via incubation in 2X SDS loading dye at 95 °C for 5 min. Samples were analyzed on 10% SDS PAGE Bis-Tris gel and imaged via Coomassie Brilliant blue staining on a BioRad ChemiDoc.

#### Immunoprecipitations with PARP1, nucleosomes, and ALC1

A 50 μL solution of 100 nM FLAG-ALC1, 50 nM unmodified or H3S10ADPr_3_ nucleosomes, and 100 nM PARP1 (unmodified, PARP1 SerADPr_long_, or PARP1 SerADPr_short_) in IP buffer (100 mM KCl, 25 mM HEPES pH 7.9, 2 mM MgCl_2_, 5% glycerol, 0.1% NP-40, 1 mM DTT) was incubated at 30 °C for 15 min. Binding reactions were then added to anti-FLAG M2 magnetic resin (MilliporeSigma; pre-equilibrated in IP Buffer) after keeping aside 5 μL as an input control for western blot analysis. Resin was incubated on an end-over-end rotator at 4 °C for 1 h, washed for 3 times for 1 min each with 0.5 mL of IP Buffer (with very gentle vortexing), and eluted via incubation in 2X SDS loading dye at 95 °C for 5 min. Samples were run on 10% SDS PAGE Bis-Tris gels, analyzed via western blot and imaged on a BioRad ChemiDoc.

### Fluorescence polarization-based peptide interaction assays

Each fluorescently-labeled H2B (unmodified, mono-, di-, tri-, and tetra-ADP-ribose at H2BS6) and H3 peptide (unmodified, mono-, di-, tri-, tetra-, or penta-ADP-ribose at H3S10) was diluted to 2 nM in a buffer containing 25 mM Tris, pH 7.5, 100 mM NaCl, 2 mM MgCl_2_, 0.001% Triton X100, and 1 mM DTT. Note, H3 elongates more efficiently than H2B and so the penta-ADP-ribosylated species could be isolated for this peptide. Peptide concentration was calculated via fluorescein extinction coefficient (A_480_ = 70,000). To analyze peptide:pan-ADP-ribose detection reagent (Af1521 macrodomain fused to rabbit Fc tag, MilliporeSigma) interaction, the pan-ADP-ribose detection reagent was titrated into each peptide to final concentrations ranging from 0-2000 nM (points represent 3x dilutions starting from 2000 nM; a higher concentration of 4000 nM was also included for mono- and di-ADP-ribosylated peptides). To analyze peptide:ALC1-macrodomain interaction, the ALC1-macrodomain was titrated into each peptide to final concentrations ranging from 0-3000 nM (points represent 3x dilutions starting from 3000 nM). Reactions were added to a black, flat-bottom 96-well plate (Corning Costar) and analyzed on a BioTek Cytation 5 imager equipped with a Green FP filter set (excitation: 485 nm, emission: 528 nm). Polarization values were converted to anisotropy using the following formula: r=(2P/(3-P)) (Lakowicz, 2006). Following background subtraction and normalization, data was then processed in GraphPad Prism using a non-linear regression analysis to obtain K_d, app_ values for each peptide:protein interaction. Error bars represent standard deviation value from three biological replicates.

### Assembly of full-length, ADP-ribosylated histones

Native chemical ligation reactions were performed by combining modified histone peptides bearing C-terminal MESNa moieties (H2BS6ADPr_1_, H2BS6ADPr_3_, H2BS6ADPr_4_, H3S10ADPr_1_, H3S10ADPr_3_, or H3S10ADPr_4_) with their corresponding recombinant C-terminal histone fragments (H2BA17C 17-125 or H3A21C 21-135). A typical reaction included 1 mM histone thioester peptide, 0.5 mM recombinant histone fragment, 20 mM TCEP, and 150 mM 2,2,2-trifluoroethanethiol (Sigma-Aldrich) in a degassed buffer of 6 M guanidine hydrochloride and 0.1 M sodium phosphate at pH 7.0. Reactions were incubated at 37 °C for 16 h and progress was monitored via RP-HPLC and ESI-MS analysis. Full-length histone products were purified on a semi-preparative C18 RP-HPLC column using a gradient from 10-80% Solvent B over 40 minutes. Fractions were analyzed via analytical C18 RP-HPLC and ESI-MS and those greater than 95% pure were pooled, lyophilized, and stored at −80 °C until use. We note that all H2B and H3 histones have an alanine to cysteine mutation at the respective ligation junction (H2BA17C and H3A21C). We have since optimized desulfurization protocols to convert this cysteine back to the native alanine residue without affecting the ADP-ribose moiety. Desulfurization will be employed in future applications of this method.

### Preparation of histone octamers

Octamers and nucleosomes were prepared as previously described (Luger et al., 1999) with several modifications. Lyophilized recombinant and semi-synthetic histones were dissolved in a buffer containing 6 M guanidine hydrochloride, 20 mM Tris, pH 7.6, and 5 mM DTT at 4 °C. H2A, H2B, H3, and H4 were combined at a ratio of 1.2:1.2:1.0:1.0, respectively, and diluted to a final concentration of 1 mg/mL of total histone. The histone mixture was then injected into a Slide-A-Lyzer MINI dialysis cassette (3.5 kDa MWCO, ThermoFisher) and dialyzed at 4 °C into Octamer Refolding Buffer (10 mM Tris, pH 7.6, 2 M NaCl, 1 mM EDTA, and 1 mM DTT) for 20 h. The cassette was placed into fresh Octamer Refolding Buffer at the 4 h and 16 h time-points during the dialysis. Next, the histone octamer solution was purified via gel filtration (Superdex 200 Increase 10/300 GL; GE Healthcare) that had been pre-equilibrated with Octamer Refolding Buffer. Injection volume did not exceed 0.5 mL to ensure efficient separation of histone octamers from sub-octamer species. Fractions containing the octamer complex (as judged by FPLC elution chromatogram and SDS–PAGE gel electrophoresis) were concentrated to 50 μM as quantified by A_280_ for unmodified nucleosomes (extinction coefficient = 44,700) or A_260_ for ADP-ribosylated nucleosomes (extinction coefficient = 13,500 x total ADP-ribose units), diluted two-fold with glycerol, and stored at a final concentration of 25 μM at −20 °C prior to nucleosome assembly. The following unique octamers were assembled for nucleosome preparation: unmodified, H2BS6ADPr_1_, H2BS6ADPr_3_, H2BS6ADPr_4_, H3S10ADPr_1_, H3S10ADPr_3_, H3S10ADPr_4_, H2BS6/H3S10ADPr_3_, H2BS6/H3S10ADPr_4_.

### Nucleosome assembly and characterization

Nucleosomes were assembled by combining 150 pmol histone octamer with 180 pmol 601 DNA in 75 μL of a buffer containing 2 M KCl, 10 mM Tris, pH 7.5, 0.1 mM EDTA, 1 mM DTT at 4 °C. The mixture was then injected into a Slide-A-Lyzer MINI dialysis button (3.5 kDa MWCO, ThermoFisher) and dialyzed against a buffer of 10 mM Tris, pH 7.0, 1.4 M KCl, 0.1 mM EDTA, 1 mM DTT at 4 °C for 1 h. Next, 350 mL of Nucleosome End Buffer (10 mM Tris, pH 7.5, 10 mM KCl, 0.1 mM EDTA, 1 mM DTT) was added at a rate of 1 mL/min. After 12 h, the cassette was dialyzed against Nucleosome End Buffer for 4 h with a fresh buffer exchange at the 2 h time-point. Following dialysis, precipitation was removed via centrifugation at 20,000 RCF for 10 min at 4 °C and A_260_ of the supernatant was measured to calculate nucleosome concentration. Note that for individual remodeling experiments, all nucleosomes were assembled on an identical 601-containing 200 bp DNA template. For competition remodeling experiments, each nucleosome was assembled on the same 601-containing 200 bp DNA template except the 15 bp at the 5¢-end were replaced with a unique priming sequence.

Nucleosome quality was analyzed by running the nucleosome on native PAGE on a 5% TBE gel in 0.5X TBE buffer (BioRad) that was run for 60 min at 150 volts. For gel loading, 10 pmol of nucleosome was diluted into 20 μL of Nucleosome End Buffer supplemented with 12% sucrose. Gels were stained with ethidium bromide and imaged on a BioRad ChemiDoc and the nucleosome band migrates around 500 bp. We noted that nucleosome migration is affected by ADP-ribose chain length. If any free 601 DNA was observed on the TBE gel, then a PstI (NEB) restriction digestion was performed to check if the free DNA was present in the nucleosome dialysate or was an artifact of the gel run. To further verify stability of ADPr throughout the histone octamer and nucleosome assembly, 2.5 pmol of nucleosome were run on a 12% SDS-PAGE gel and immuno-blot analysis was performed to detect ADP-ribose, H2B, and H3. ADP-ribosylated H2B and H3 proteins exhibit distinct migration profiles relative to the unmodified species, confirming that they are homogenously modified.

### Western blot protocol

SDS-PAGE gels were transferred to PVDF membranes at 100 volts for 1 h at 4 °C using a wet transfer protocol in Towbin Buffer (25 mM Tris, 192 mM glycine, 20% methanol, pH 8.3). Blots were then blocked for 1 h at 25 °C with 5% non-fat dry milk (BioRad) in TBST (50 mM Tris, pH 7.5, 150 mM NaCl, 0.1% Tween20) prior to incubation with primary antibodies for 12 h at 4 °C. Following primary antibody binding, blots were washed 3 times for 5 min each with TBST and then incubated with the appropriate fluorescent or HRP-conjugated secondary antibody for 1 h at 25 °C. Blots were then washed 3 times for a total of 15 min with TBST and imaged on a BioRad ChemiDoc. All antibodies used in this study and corresponding dilutions can be found in Supplementary Table 5.

### Restriction enzyme accessibility-based nucleosome remodeling assay

REA assays and analysis were performed as previously described (Dann et al., 2017) with several modifications. Nucleosome remodeling reactions (25 μL) were carried out in REA Buffer (12 mM HEPES, pH 7.9, 4 mM Tris, pH 7.5, 60 mM KCl, 10 mM MgCl_2_, 10% glycerol, and 0.02% NP-40) including 1 μL of PstI (NEB, at 100,000 U/mL), 2 mM ATP (Sigma-Aldrich), and final concentrations of 4 nM ALC1 or 10 nM CHD4 and 20 nM of the desired nucleosome substrate. The reaction was incubated for 5 min prior to addition of chromatin remodeler to ensure that any trace amount of free DNA from the nucleosome assembly was digested prior to initiating the reaction. This is required to ensure that free DNA digestion can be assigned as background activity and is not interpreted as enzyme-dependent nucleosome remodeling in data processing. To each reaction, 37.5 μL of Quench Buffer (20 mM Tris, pH 7.5, 70 mM EDTA, pH 8, 2% SDS, 10% glycerol) was added at time points of 0, 3, 6, 18, 36, and 60 min. Samples were then deproteinized with 30 U/mL proteinase K (NEB) for 1 h at 37 °C. DNA purification was performed using the Qiagen PCR purification kit following manufacturer’s protocols. Purple Gel Loading Dye (6X, NEB) was added to a final concentration of 1X to the quenched reaction and samples were loaded onto a 5% TBE gel and run for 60 min at 150 volts in 0.5X TBE Buffer (BioRad). Gels were stained with ethidium bromide and imaged on a BioRad ChemiDoc. Gel densitometry measurements were performed using ImageJ. For each lane, the total densitometry signal was calculated by adding the densitometry values corresponding to the PstI-digested species (lower band) and undigested species (upper band). The fraction unremodeled value for each lane was then calculated using the following formula:

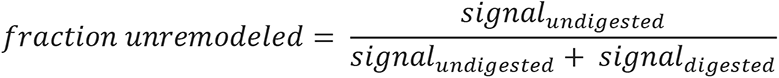

For each chromatin remodeling reaction, activity at the zero time point was considered background activity (described above) and that value of fraction unremodeled was denoted as the reference for normalizing values from other time points in the corresponding reaction. Data was performed in biological triplicate and fit into one-phase exponential decay equation in GraphPad Prism to obtain the remodeling plots and corresponding k values (Supplementary Table 2), where k denotes the rate constant for the exponential decay. For calculation of k values, plateau was constrained to zero and k>0.

To probe ALC1 activation by freely diffusing ADP-ribosylated peptides, the H3S10ADPr_4_ or H2BS6ADPr_4_ (amino acids 1-20 or 1-16, respectively) peptides were added to the REA Buffer at 10, 40, or 200 nM. Remodeling reactions were then carried out as described for 1 h on unmodified nucleosomes (Supplementary Fig. 5c). Control reactions were also set up with the same concentrations of unmodified versions of the corresponding peptides. A full time-course (0, 3, 6, 18, 36, 60 min) was performed at the 40 nM peptide concentration. This concentration was selected for complete analysis (Fig. 5c) because the final ADP-ribose concentration is equivalent to that of the modified nucleosome assays (20 nM nucleosome x 2 ADP-ribosylated histone tails per nucleosome). All reactions and controls were performed in triplicate.

To probe ALC1 activation by Asp-/Glu-auto-ADP-ribosylated PARP1, we added 5, 20, 50, 100, or 200 nM PARP1 and 2 mM NAD^+^ to the remodeling assays. Remodeling reactions were then carried out as described for 1 h on unmodified nucleosomes (Supplementary Fig. 5e). A full time-course (0, 3, 6, 18, 36, 60 min) was performed at 20 and 100 nM PARP1 concentration in triplicate (Supplementary Fig. 5b). These concentrations were selected for complete analysis because the stimulatory effect of ADP-ribosylated PARP1 on nucleosome remodeling by ALC1 plateaued at around 50 nM. Western blots were performed using the PARP1 antibody and the pan-ADPr-detection reagent at time-points 0, 3, 6, 18, 36, and 60 min to quantify conversion of PARP1 in the reaction to the auto-ADP-ribosylated species. Similar titrations were carried out for serine-linked auto-ADP-ribosylated PARP1 species and the stimulatory effect seemed to plateau around 100nM, thus the full time-course (0, 3, 6, 18, 36, 60 min) remodeling experiments were performed at 100nM serine auto-modified PARP1 concentration (Supplementary Fig. 5g).

### Hydroxylamine treatment

A fresh stock solution of 3.3 M hydroxylamine was prepared in 10 mM Tris in water and adjusted to pH ∼6 using filtered 5 M KOH. Automodified PARP1 constructs (1 μM) were added to a buffer containing 50 mM Tris (pH 7.5), 20 mM NaCl and 2 mM MgCl_2_. Hydroxylamine was added to this solution to a final concentration of 0.8 M and the solution was incubated at room temperature for 1 h. The reaction was quenched using 0.3% HCl. SDS-PAGE loading dye was added to the samples at a final concentration of 1X and boiled before being run on an SDS-PAGE gel. The bands on the gels were visualized via silver-stain.

### Nucleosome remodeling competition assay

Two nucleosome substrate pools were prepared for the competition assays, with each substrate pool containing seven unique species. The first pool (H2B Pool) included H2BS6ADPr_1_, H2BS6ADPr_3_, and H2BS6ADPr_4_ nucleosomes, each of which contained a unique 5′ priming site as outlined in the ‘601 DNA preparation’ section above. The second pool (H3 Pool) included H3S10ADPr_1_, H3S10ADPr_3_, and H3S10ADPr_4_ nucleosomes, each of which contained a unique 5′ priming site. Each pool also included two unmodified nucleosomes assembled on unique templates to serve as internal reproducibility controls. Free DNA templates with the PstI site and an unmodified nucleosome without the PstI site were also included as internal controls for PstI activity and data normalization, respectively. To ensure that PCR amplification artifacts do not influence cycle threshold determination, we selected primer:template pairs with similar primer efficiencies (Supplementary Fig. 5h).

Each nucleosome substrate pool was prepared by combining equal volumes of each nucleosome (stock solutions = 250 nM) or free DNA species (stock solutions = 250 nM). Therefore, the total species concentration in each assembled substrate pool is 250 nM (∼36 nM per species). The final total nucleosome species concentration used in remodeling assays was 20 nM. ALC1 was used at a concentration of 4 nM for the H2B substrate pool and 8 nM for the H3 substrate pool. Remodeling assays were carried out, quenched at six different time points (0, 3, 6, 18, 36, 60 min) and DNA was isolated as described in the ‘Restriction enzyme accessibility-based nucleosome remodeling assay’ section. Real-time PCR was then performed with each unique primer pair according to manufacturer’s protocols (iTaq Universal SYBR Green Supermix, BioRad) to quantify undigested (that is, unremodeled) template for each unique species at every time point. Fold-decrease in template quantity from t = 0 to t = x was calculated by determining the ΔΔCt for a species of interest relative to the unmodified nucleosome lacking the PstI site. Note: the template lacking PstI site cannot be digested and thus serves as an internal control for ΔΔCt calculation. Fold-decrease in template quantity was then converted to fraction unremodeled. Each competition assay was performed in triplicate and data points take into account an average of three independent amplifications for each primer pair (see Supplementary Dataset for primer pair:substrate combinations). The data was processed in GraphPad Prism and fit into a one-phase exponential decay equation with plateau constrained to zero and k>0 to obtain the remodeling plots and corresponding k values (Figure 5g, h, and Supplementary Table 3).

### Mammalian cell culture

HEK293T cells (ATCC) were culture in high-glucose DMEM (MilliporeSigma) supplemented with 10% Fetal Bovine Serum (Gibco), 100 units/mL of penicillin (Sigma), and 100 μg/mL of streptomycin (Sigma). Cells were maintained at 37°C and 5% CO_2_ and passaged/frozen down according to manufacturer’s protocols (ATCC). Plasmid transfection was accomplished with Lipofectamine 2000 according to manufacturer’s protocols (Invitrogen).

#### Generation of ALC1 knockout cell lines

CRISPR-Cas9 plasmids (pSpCas9(BB)-2A-Puro (PX459) v2.0; Addgene plasmid #: 62988) targeting the ALC1 gene (for gRNA targeting sequences, see Supplementary Table 4) were transfected into HEK293T cells. Targeting sequences were obtained using the Genetic Perturbation Platform (Broad Institute). After 24 h, 2 μg/mL of puromycin (Sigma) was added to growth medium and cells were selected for 48 h. Puromycin was then removed, dead cells were washed away, and the adhering live cells were left to recover for 24 h prior to dilution for single colony selection. Clones were screened via western blot for ALC1 and those with no detectable ALC1 were frozen down and stored in liquid nitrogen.

### Nuclear lysate preparation

Nuclear lysate was prepared as previously described (Carey et al., 2009) with some modifications. The cells were dounced with a B-type pestle (Kontle Glass Co) until they were lysed. Lysis was confirmed by staining with Trypan Blue dye and visualizing under a microscope. The cell number was estimated using a hemocytometer and the volumes of the different buffers were added depending on that. The nuclear lysate was homogenized using pestle B until it was properly resuspended in Buffer C. The crude nuclear lysate was dialyzed into Buffer D in a dialysis tubing (FisherScientific, 6-8 kDa MWCO) at 4 °C, and dialysis was stopped at first signs of precipitation (around 3-4h).

### Nuclear lysate nucleosome remodeling assay

Nucleosome remodeling reactions (25 μL) were carried out in REA Buffer with 1 μL of PstI (NEB, at 100,000 U/mL), 2 mM ATP, and 8 μL of nuclear lysate derived from either wild-type or ALC1 knockout HEK293T cells and 20 nM of the desired nucleosome substrate. The reactions were carried out at 30°C and quenched with Quench Buffer at 0 min and 60 min time points. The 601 DNA was isolated as described in ‘Restriction enzyme accessibility-based nucleosome remodeling assay’ section and analyzed on a 5% TBE gel. Western blot analyses of the ADPr profile of each nucleosome employed in this assay were carried out after incubation with or without either nuclear lysate under identical reaction conditions.

## Supplementary Information

**Supplementary figures 1-6** and **Supplementary Tables 1-5** and (along with accompanying legends) are provided as a separate document.

**Supplementary Dataset** contains all peak integration values from ADP-ribosylation assays, fluorescence polarization values from peptide-macrodomain interaction assays, densitometry values from single-substrate chromatin remodeling assays, and cycle threshold (C_t_) values from multi-substrate chromatin remodeling assays reported in this study.

## References

Aberle, L., Kruger, A., Reber, J.M., Lippmann, M., Hufnagel, M., Schmalz, M., Trussina, I., Schlesiger, S., Zubel, T., Schutz, K., et al. (2020). PARP1 catalytic variants reveal branching and chain length-specific functions of poly(ADP-ribose) in cellular physiology and stress response. Nucleic Acids Res 48, 10015–10033.

Ahel, D., Horejsi, Z., Wiechens, N., Polo, S.E., Garcia-Wilson, E., Ahel, I., Flynn, H., Skehel, M., West, S.C., Jackson, S.P., et al. (2009). Poly(ADP-ribose)-dependent regulation of DNA repair by the chromatin remodeling enzyme ALC1. Science 325, 1240–1243.

Benjamin, R.C., and Gill, D.M. (1980). Poly(ADP-ribose) synthesis in vitro programmed by damaged DNA. A comparison of DNA molecules containing different types of strand breaks. J Biol Chem 255, 10502–10508.

Bilokapic, S., Suskiewicz, M.J., Ahel, I., and Halic, M. (2020). Bridging of DNA breaks activates PARP2-HPF1 to modify chromatin. Nature 585, 609–613.

Blessing, C., Mandemaker, I.K., Gonzalez-Leal, C., Preisser, J., Schomburg, A., and Ladurner, A.G. (2020). The Oncogenic Helicase ALC1 Regulates PARP Inhibitor Potency by Trapping PARP2 at DNA Breaks. Mol Cell 80, 862–875 e866.

Bonfiglio, J.J., Fontana, P., Zhang, Q., Colby, T., Gibbs-Seymour, I., Atanassov, I., Bartlett, E., Zaja, R., Ahel, I., and Matic, I. (2017). Serine ADP-Ribosylation Depends on HPF1. Mol Cell 65, 932–940 e936.

Bonfiglio, J.J., Leidecker, O., Dauben, H., Longarini, E.J., Colby, T., San Segundo-Acosta, P., Perez, K.A., and Matic, I. (2020). An HPF1/PARP1-Based Chemical Biology Strategy for Exploring ADP-Ribosylation. Cell 183, 1086–1102 e1023.

Bowman, G.D., and Poirier, M.G. (2015). Post-translational modifications of histones that influence nucleosome dynamics. Chem Rev 115, 2274–2295.

Carey, M.F., Peterson, C.L., and Smale, S.T. (2009). Dignam and Roeder nuclear extract preparation. Cold Spring Harb Protoc 2009, pdb prot5330.

Chen, Q., Kassab, M.A., Dantzer, F., and Yu, X. (2018). PARP2 mediates branched poly ADP-ribosylation in response to DNA damage. Nat Commun 9, 3233.

Chou, D.M., Adamson, B., Dephoure, N.E., Tan, X., Nottke, A.C., Hurov, K.E., Gygi, S.P., Colaiacovo, M.P., and Elledge, S.J. (2010). A chromatin localization screen reveals poly (ADP ribose)-regulated recruitment of the repressive polycomb and NuRD complexes to sites of DNA damage. Proc Natl Acad Sci U S A 107, 18475–18480.

Clapier, C.R., and Cairns, B.R. (2012). Regulation of ISWI involves inhibitory modules antagonized by nucleosomal epitopes. Nature 492, 280–284.

Daniels, C.M., Ong, S.E., and Leung, A.K. (2015). The Promise of Proteomics for the Study of ADP-Ribosylation. Mol Cell 58, 911–924.

Dann, G.P., Liszczak, G.P., Bagert, J.D., Muller, M.M., Nguyen, U.T.T., Wojcik, F., Brown, Z.Z., Bos, J., Panchenko, T., Pihl, R., et al. (2017). ISWI chromatin remodellers sense nucleosome modifications to determine substrate preference. Nature 548, 607–611.

Dawicki-McKenna, J.M., Langelier, M.F., DeNizio, J.E., Riccio, A.A., Cao, C.D., Karch, K.R., McCauley, M., Steffen, J.D., Black, B.E., and Pascal, J.M. (2015). PARP-1 Activation Requires Local Unfolding of an Autoinhibitory Domain. Mol Cell 60, 755–768.

Fontana, P., Bonfiglio, J.J., Palazzo, L., Bartlett, E., Matic, I., and Ahel, I. (2017). Serine ADP-ribosylation reversal by the hydrolase ARH3. Elife 6.

Gibbs-Seymour, I., Fontana, P., Rack, J.G.M., and Ahel, I. (2016). HPF1/C4orf27 Is a PARP-1-Interacting Protein that Regulates PARP-1 ADP-Ribosylation Activity. Mol Cell 62, 432–442.

Gibson, B.A., Conrad, L.B., Huang, D., and Kraus, W.L. (2017). Generation and Characterization of Recombinant Antibody-like ADP-Ribose Binding Proteins. Biochemistry 56, 6305–6316.

Gottschalk, A.J., Timinszky, G., Kong, S.E., Jin, J., Cai, Y., Swanson, S.K., Washburn, M.P., Florens, L., Ladurner, A.G., Conaway, J.W., et al. (2009). Poly(ADP-ribosyl)ation directs recruitment and activation of an ATP-dependent chromatin remodeler. Proc Natl Acad Sci U S A 106, 13770–13774.

Gottschalk, A.J., Trivedi, R.D., Conaway, J.W., and Conaway, R.C. (2012). Activation of the SNF2 family ATPase ALC1 by poly(ADP-ribose) in a stable ALC1.PARP1.nucleosome intermediate. J Biol Chem 287, 43527–43532.

Gupte, R., Liu, Z., and Kraus, W.L. (2017). PARPs and ADP-ribosylation: recent advances linking molecular functions to biological outcomes. Genes Dev 31, 101–126.

Hauk, G., McKnight, J.N., Nodelman, I.M., and Bowman, G.D. (2010). The chromodomains of the Chd1 chromatin remodeler regulate DNA access to the ATPase motor. Mol Cell 39, 711–723.

He, X., Fan, H.Y., Narlikar, G.J., and Kingston, R.E. (2006). Human ACF1 alters the remodeling strategy of SNF2h. J Biol Chem 281, 28636–28647.

Hein, M.Y., Hubner, N.C., Poser, I., Cox, J., Nagaraj, N., Toyoda, Y., Gak, I.A., Weisswange, I., Mansfeld, J., Buchholz, F., et al. (2015). A human interactome in three quantitative dimensions organized by stoichiometries and abundances. Cell 163, 712–723.

Huletsky, A., de Murcia, G., Muller, S., Hengartner, M., Menard, L., Lamarre, D., and Poirier, G.G. (1989). The effect of poly(ADP-ribosyl)ation on native and H1-depleted chromatin. A role of poly(ADP-ribosyl)ation on core nucleosome structure. J Biol Chem 264, 8878–8886.

Juhasz, S., Smith, R., Schauer, T., Spekhardt, D., Mamar, H., Zentout, S., Chapuis, C., Huet, S., and Timinszky, G. (2020). The chromatin remodeler ALC1 underlies resistance to PARP inhibitor treatment. Sci Adv 6.

Kim, M.Y., Mauro, S., Gevry, N., Lis, J.T., and Kraus, W.L. (2004). NAD+-dependent modulation of chromatin structure and transcription by nucleosome binding properties of PARP-1. Cell 119, 803–814.

Lakowicz, J.R. (2006). Principles of Fluorescence Spectroscopy (New York, New York: Springer Science+Business Media, LLC).

Langelier, M.F., Planck, J.L., Roy, S., and Pascal, J.M. (2012). Structural basis for DNA damage-dependent poly(ADP-ribosyl)ation by human PARP-1. Science 336, 728–732.

Larsen, S.C., Hendriks, I.A., Lyon, D., Jensen, L.J., and Nielsen, M.L. (2018). Systems-wide Analysis of Serine ADP-Ribosylation Reveals Widespread Occurrence and Site-Specific Overlap with Phosphorylation. Cell Rep 24, 2493–2505 e2494.

Lehmann, L.C., Hewitt, G., Aibara, S., Leitner, A., Marklund, E., Maslen, S.L., Maturi, V., Chen, Y., van der Spoel, D., Skehel, J.M., et al. (2017). Mechanistic Insights into Autoinhibition of the Oncogenic Chromatin Remodeler ALC1. Mol Cell 68, 847–859 e847.

Leidecker, O., Bonfiglio, J.J., Colby, T., Zhang, Q., Atanassov, I., Zaja, R., Palazzo, L., Stockum, A., Ahel, I., and Matic, I. (2016). Serine is a new target residue for endogenous ADP-ribosylation on histones. Nat Chem Biol 12, 998–1000.

Liszczak, G., Diehl, K.L., Dann, G.P., and Muir, T.W. (2018). Acetylation blocks DNA damage-induced chromatin ADP-ribosylation. Nat Chem Biol 14, 837–840.

Lord, C.J., and Ashworth, A. (2017). PARP inhibitors: Synthetic lethality in the clinic. Science 355, 1152–1158.

Luger, K., Rechsteiner, T.J., and Richmond, T.J. (1999). Preparation of nucleosome core particle from recombinant histones. Methods Enzymol 304, 3–19.

Luijsterburg, M.S., de Krijger, I., Wiegant, W.W., Shah, R.G., Smeenk, G., de Groot, A.J.L., Pines, A., Vertegaal, A.C.O., Jacobs, J.J.L., Shah, G.M., et al. (2016). PARP1 Links CHD2-Mediated Chromatin Expansion and H3.3 Deposition to DNA Repair by Non-homologous End-Joining. Mol Cell 61, 547–562.

Palazzo, L., Leidecker, O., Prokhorova, E., Dauben, H., Matic, I., and Ahel, I. (2018). Serine is the major residue for ADP-ribosylation upon DNA damage. Elife 7.

Poirier, G.G., de Murcia, G., Jongstra-Bilen, J., Niedergang, C., and Mandel, P. (1982). Poly(ADP-ribosyl)ation of polynucleosomes causes relaxation of chromatin structure. Proc Natl Acad Sci U S A 79, 3423–3427.

Ray Chaudhuri, A., and Nussenzweig, A. (2017). The multifaceted roles of PARP1 in DNA repair and chromatin remodelling. Nat Rev Mol Cell Biol 18, 610–621.

Rudolph, J., Roberts, G., Muthurajan, U.M., and Luger, K. (2021). HPF1 and nucleosomes mediate a dramatic switch in activity of PARP1 from polymerase to hydrolase. Elife 10.

Singh, H.R., Nardozza, A.P., Moller, I.R., Knobloch, G., Kistemaker, H.A.V., Hassler, M., Harrer, N., Blessing, C., Eustermann, S., Kotthoff, C., et al. (2017). A Poly-ADP-Ribose Trigger Releases the Auto-Inhibition of a Chromatin Remodeling Oncogene. Mol Cell 68, 860–871 e867.

Slade, D., Dunstan, M.S., Barkauskaite, E., Weston, R., Lafite, P., Dixon, N., Ahel, M., Leys, D., and Ahel, I. (2011). The structure and catalytic mechanism of a poly(ADP-ribose) glycohydrolase. Nature 477, 616–620.

Smeenk, G., Wiegant, W.W., Marteijn, J.A., Luijsterburg, M.S., Sroczynski, N., Costelloe, T., Romeijn, R.J., Pastink, A., Mailand, N., Vermeulen, W., et al. (2013). Poly(ADP-ribosyl)ation links the chromatin remodeler SMARCA5/SNF2H to RNF168-dependent DNA damage signaling. J Cell Sci 126, 889–903.

Smith, R., Sellou, H., Chapuis, C., Huet, S., and Timinszky, G. (2018). CHD3 and CHD4 recruitment and chromatin remodeling activity at DNA breaks is promoted by early poly(ADP-ribose)-dependent chromatin relaxation. Nucleic Acids Res 46, 6087–6098.

Suskiewicz, M.J., Zobel, F., Ogden, T.E.H., Fontana, P., Ariza, A., Yang, J.C., Zhu, K., Bracken, L., Hawthorne, W.J., Ahel, D., et al. (2020). HPF1 completes the PARP active site for DNA damage-induced ADP-ribosylation. Nature 579, 598–602.

Teloni, F., and Altmeyer, M. (2016). Readers of poly(ADP-ribose): designed to be fit for purpose. Nucleic Acids Res 44, 993–1006.

Tulin, A., and Spradling, A. (2003). Chromatin loosening by poly(ADP)-ribose polymerase (PARP) at Drosophila puff loci. Science 299, 560–562.

Verma, P., Zhou, Y., Cao, Z., Deraska, P.V., Deb, M., Arai, E., Li, W., Shao, Y., Puentes, L., Li, Y., et al. (2021). ALC1 links chromatin accessibility to PARP inhibitor response in homologous recombination-deficient cells. Nat Cell Biol 23, 160–171.

Voorneveld, J., Rack, J.G.M., Ahel, I., Overkleeft, H.S., van der Marel, G.A., and Filippov, D.V. (2018). Synthetic alpha- and beta-Ser-ADP-ribosylated Peptides Reveal alpha-Ser-ADPr as the Native Epimer. Org Lett 20, 4140–4143.

Zhu, A., Li, X., Bai, L., Zhu, G., Guo, Y., Lin, J., Cui, Y., Tian, G., Zhang, L., Wang, J., et al. (2020). Biomimetic alpha-selective ribosylation enables two-step modular synthesis of biologically important ADP-ribosylated peptides. Nat Commun 11, 5600.

