## Supplemental Information for "Serine ADP-ribosylation marks nucleosomes for ALC1-dependent chromatin remodeling"

**a**

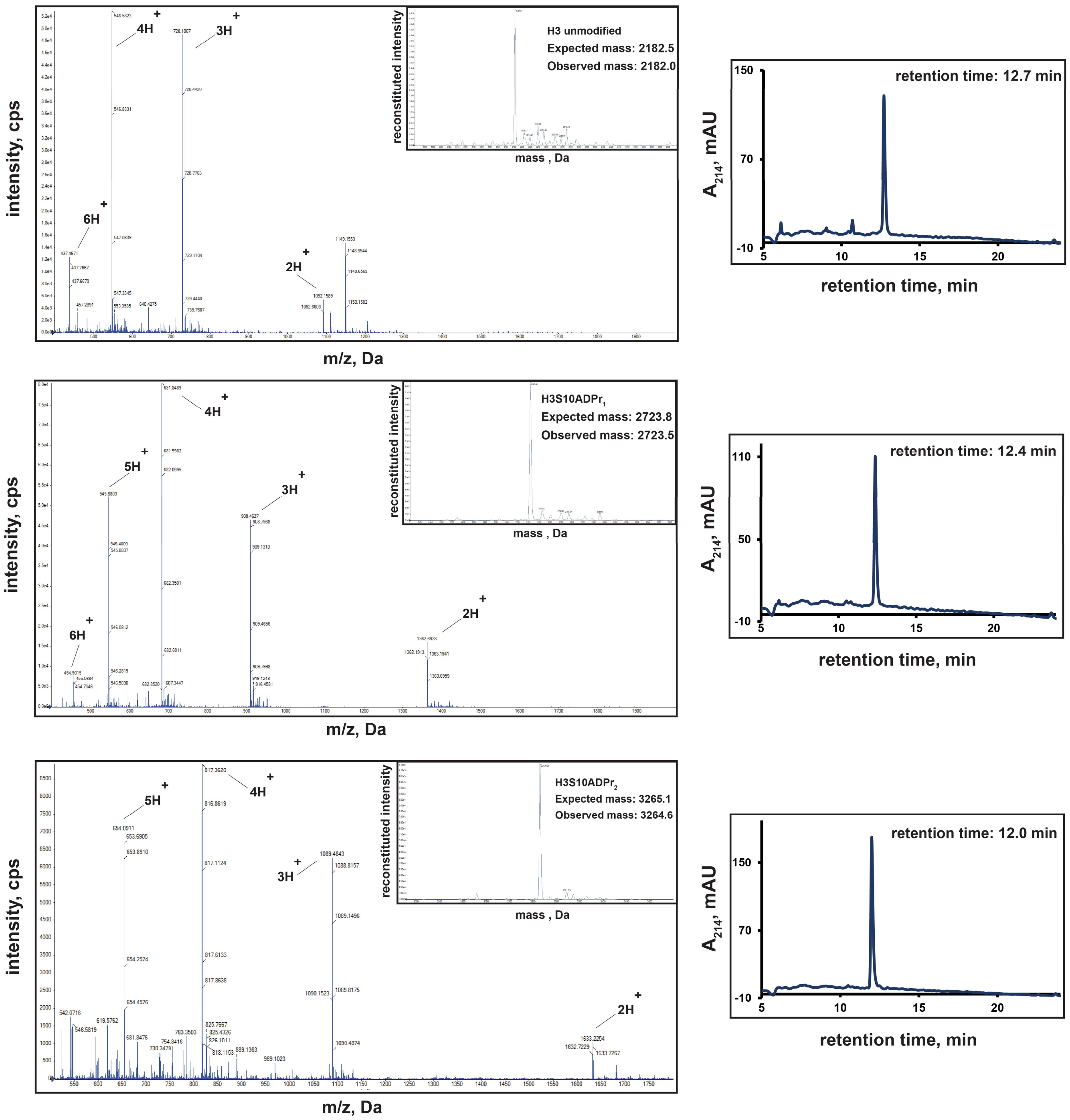
H3 unmodified
Expected mass: 2182.5
Observed mass: 2182.0
reconstituted intensity
mass , Da
intensity, cps
m/z, Da
6H+
4H+
3H+
2H+
retention time: 12.7 min
A214, mAU
retention time, min
H3S10ADPr1
Expected mass: 2723.8
Observed mass: 2723.5
5H+
retention time: 12.4 min
H3S10ADPr2
Expected mass: 3265.1
Observed mass: 3264.6
retention time: 12.0 min

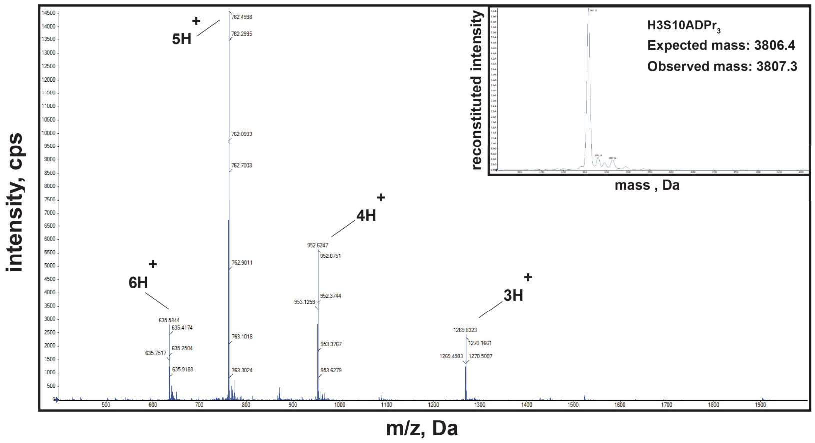
intensity, cps
m/z, Da
5H +
4H +
6H +
3H +
reconstituted intensity
mass , Da
H3S10ADPr$_3$
Expected mass: 3806.4
Observed mass: 3807.3

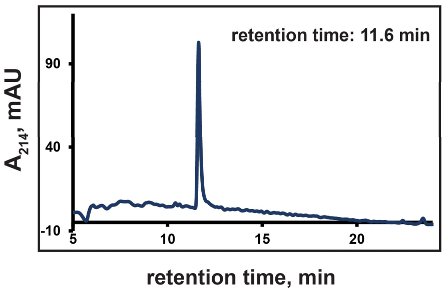
retention time: 11.6 min
A$_{214}$, mAU
retention time, min

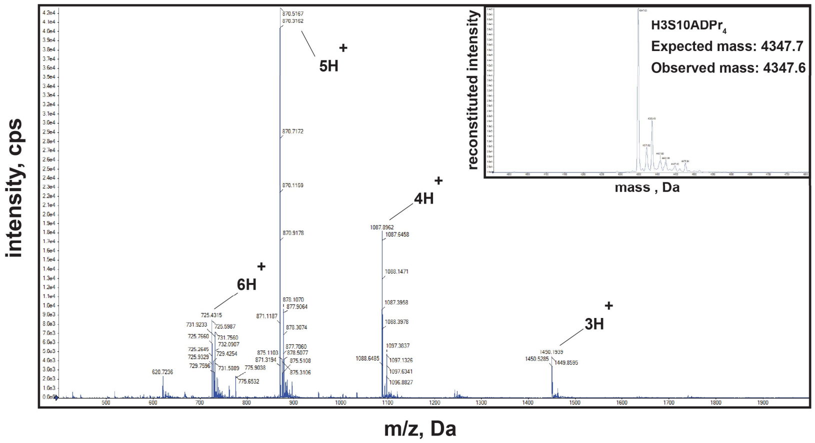
intensity, cps
m/z, Da
5H +
4H +
6H +
3H +
reconstituted intensity
mass , Da
H3S10ADPr$_4$
Expected mass: 4347.7
Observed mass: 4347.6

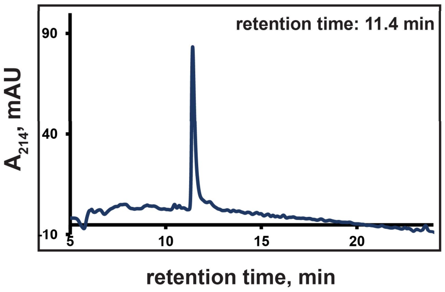
retention time: 11.4 min
A$_{214}$, mAU
retention time, min

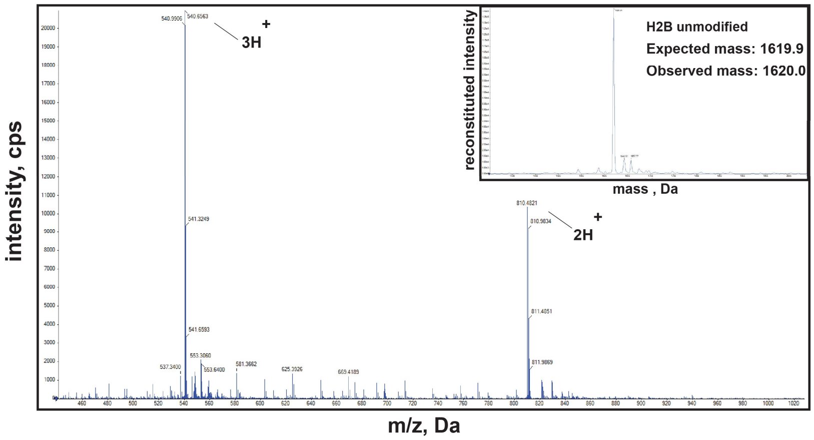
intensity, cps
m/z, Da
3H +
2H +
reconstituted intensity
mass , Da
H2B unmodified
Expected mass: 1619.9
Observed mass: 1620.0

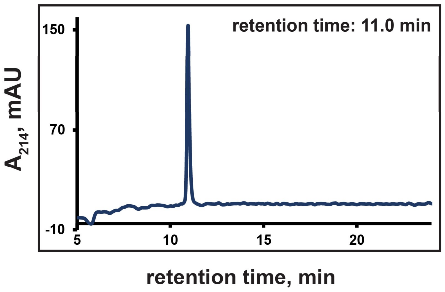
retention time: 11.0 min
A$_{214}$, mAU
retention time, min

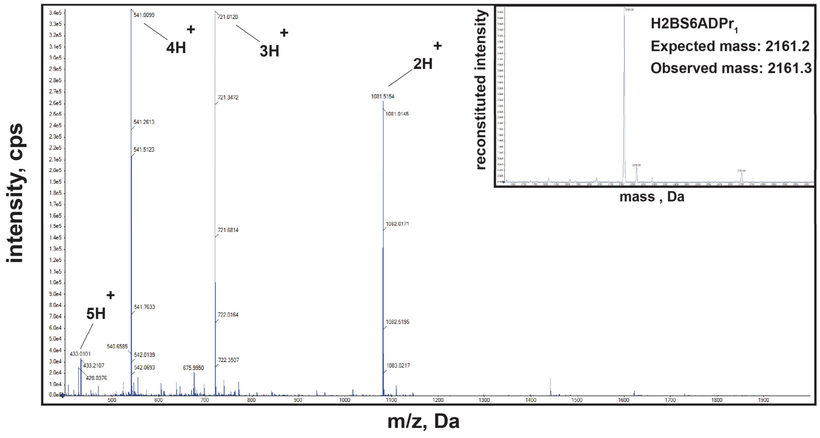
intensity, cps
m/z, Da
4H +
3H +
2H +
5H +
reconstituted intensity
mass , Da
H2BS6ADPr$_1$
Expected mass: 2161.2
Observed mass: 2161.3

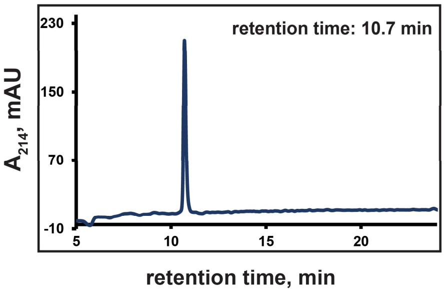
retention time: 10.7 min
A$_{214}$, mAU
retention time, min

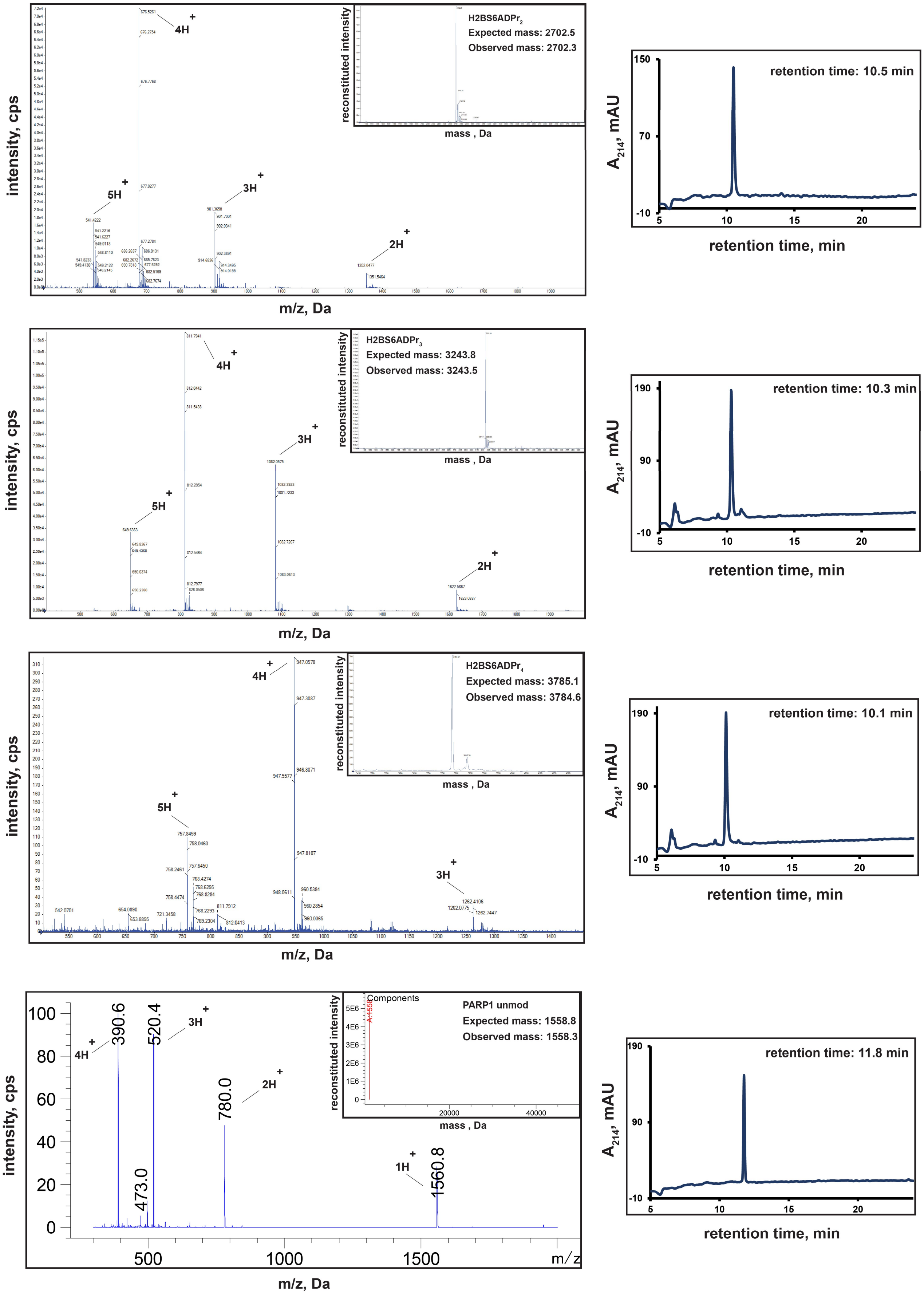
H2BS6ADPr2
Expected mass: 2702.5
Observed mass: 2702.3
reconstituted intensity
mass , Da
intensity, cps
m/z, Da
5H+
4H+
3H+
2H+
retention time: 10.5 min
A214, mAU
retention time, min
H2BS6ADPr3
Expected mass: 3243.8
Observed mass: 3243.5
retention time: 10.3 min
H2BS6ADPr4
Expected mass: 3785.1
Observed mass: 3784.6
retention time: 10.1 min
Components
PARP1 unmod
Expected mass: 1558.8
Observed mass: 1558.3
390.6
520.4
780.0
473.0
1560.8
1H+
retention time: 11.8 min

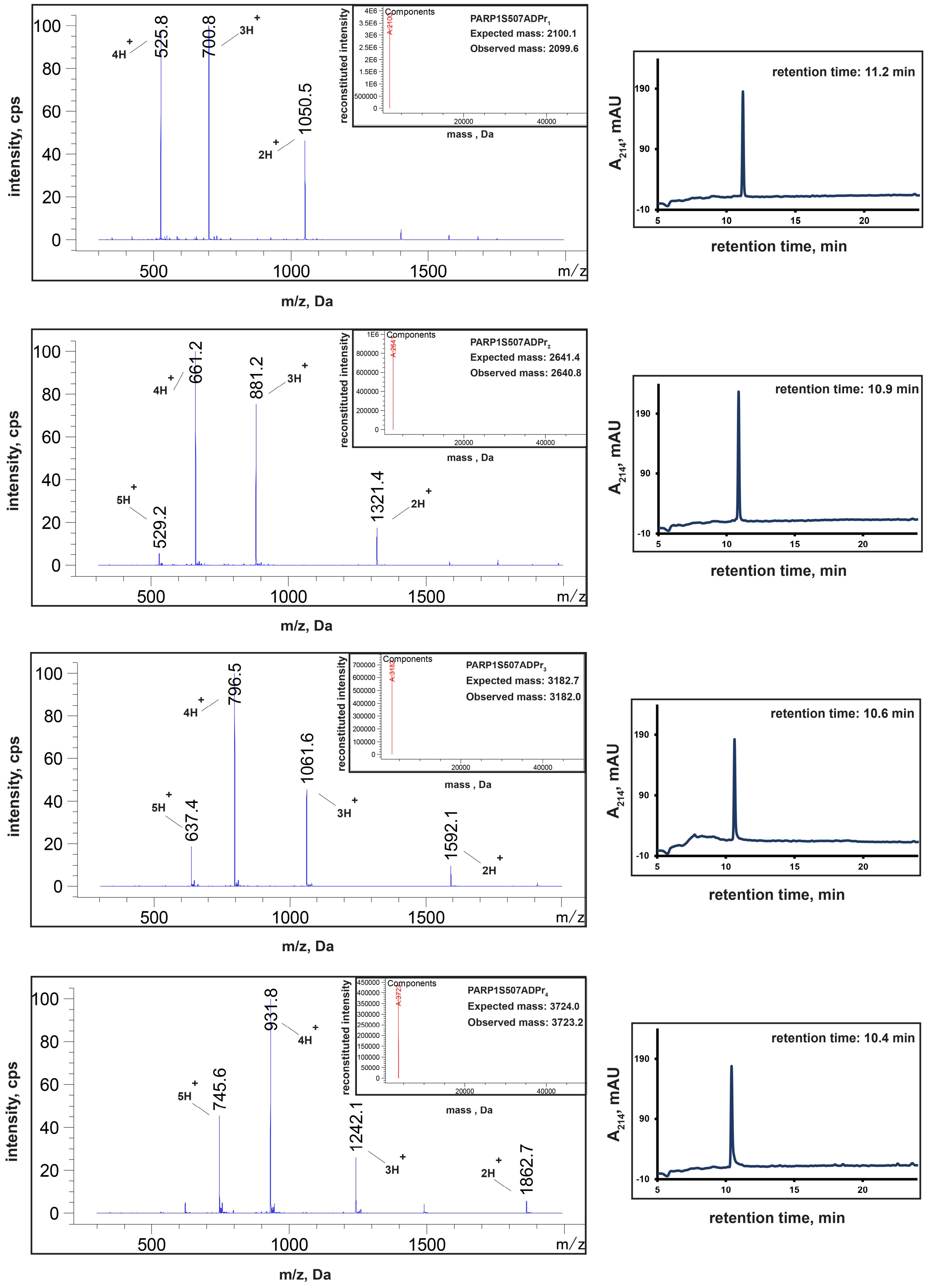
PARP1S507ADPr1
Expected mass: 2100.1
Observed mass: 2099.6
Components
reconstituted intensity
mass , Da
4H+ 525.8
3H+ 700.8
2H+ 1050.5
intensity, cps
m/z, Da
retention time: 11.2 min
A214, mAU
retention time, min
PARP1S507ADPr2
Expected mass: 2641.4
Observed mass: 2640.8
5H+ 529.2
4H+ 661.2
3H+ 881.2
2H+ 1321.4
retention time: 10.9 min
PARP1S507ADPr3
Expected mass: 3182.7
Observed mass: 3182.0
5H+ 637.4
4H+ 796.5
3H+ 1061.6
2H+ 1592.1
retention time: 10.6 min
PARP1S507ADPr4
Expected mass: 3724.0
Observed mass: 3723.2
5H+ 745.6
4H+ 931.8
3H+ 1242.1
2H+ 1862.7
retention time: 10.4 min

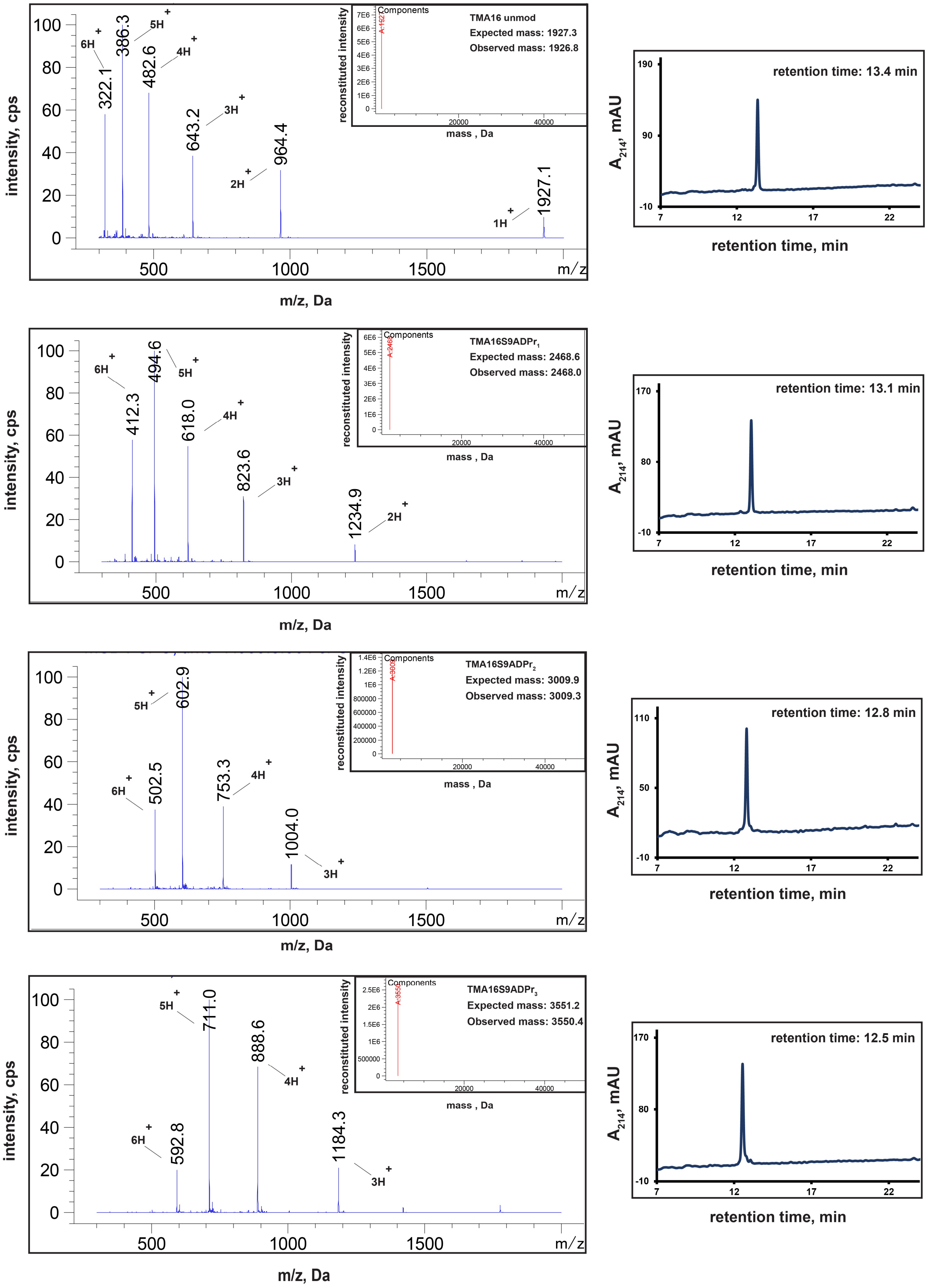
TMA16 unmod
Expected mass: 1927.3
Observed mass: 1926.8
retention time: 13.4 min
TMA16S9ADPr1
Expected mass: 2468.6
Observed mass: 2468.0
retention time: 13.1 min
TMA16S9ADPr2
Expected mass: 3009.9
Observed mass: 3009.3
retention time: 12.8 min
TMA16S9ADPr3
Expected mass: 3551.2
Observed mass: 3550.4
retention time: 12.5 min
Components
reconstituted intensity
mass , Da
intensity, cps
m/z, Da
A214, mAU
retention time, min

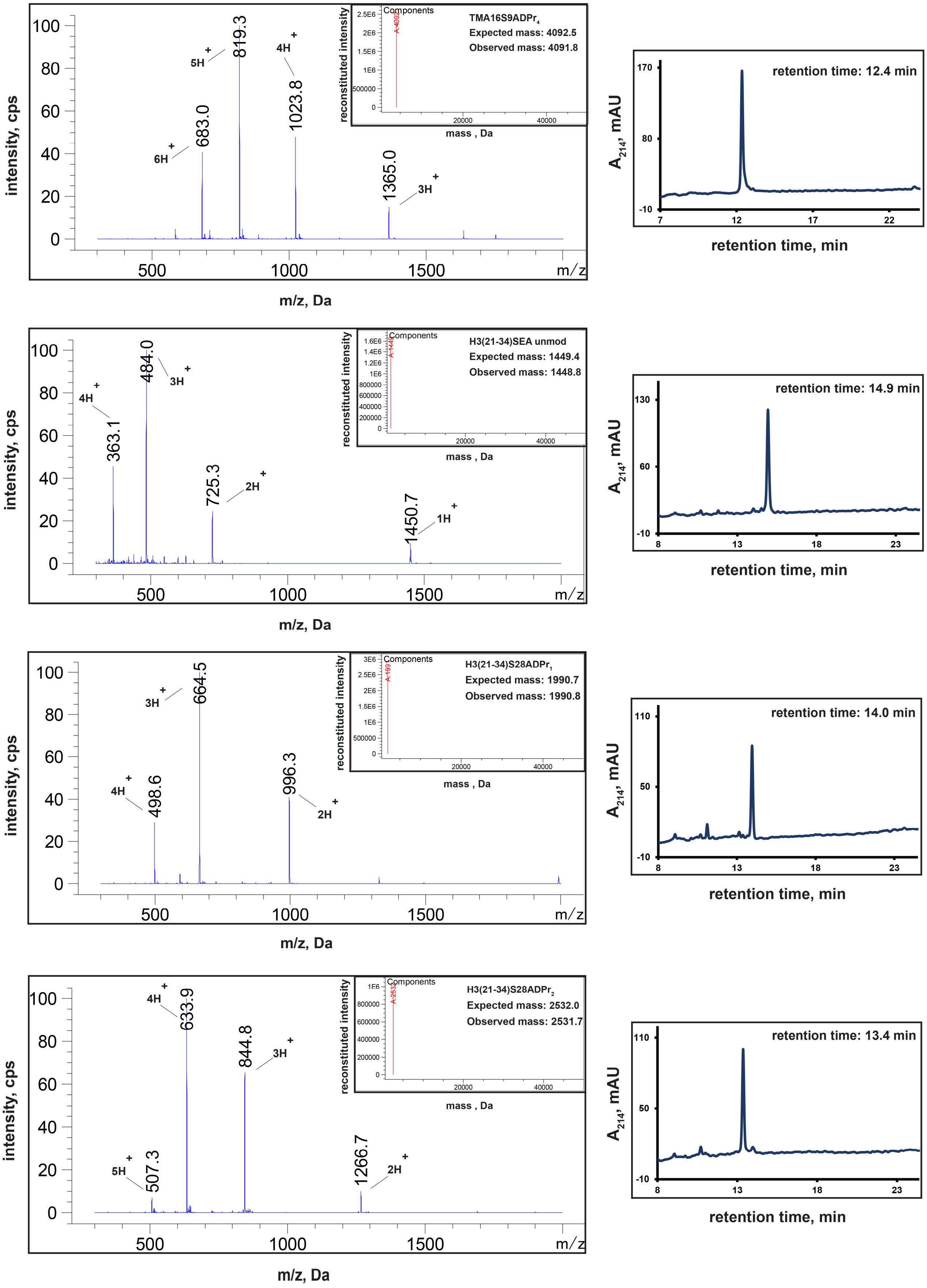
TMA16S9ADPr$_4$
Expected mass: 4092.5
Observed mass: 4091.8
Components
reconstituted intensity
mass , Da
intensity, cps
m/z, Da
6H+ 683.0
5H+ 819.3
4H+ 1023.8
1365.0 3H+
retention time: 12.4 min
$A_{214}$, mAU
retention time, min
H3(21-34)SEA unmod
Expected mass: 1449.4
Observed mass: 1448.8
4H+ 363.1
484.0 3H+
725.3 2H+
1450.7 1H+
retention time: 14.9 min
H3(21-34)S28ADPr$_1$
Expected mass: 1990.7
Observed mass: 1990.8
4H+ 498.6
3H+ 664.5
996.3 2H+
retention time: 14.0 min
H3(21-34)S28ADPr$_2$
Expected mass: 2532.0
Observed mass: 2531.7
5H+ 507.3
4H+ 633.9
844.8 3H+
1266.7 2H+
retention time: 13.4 min

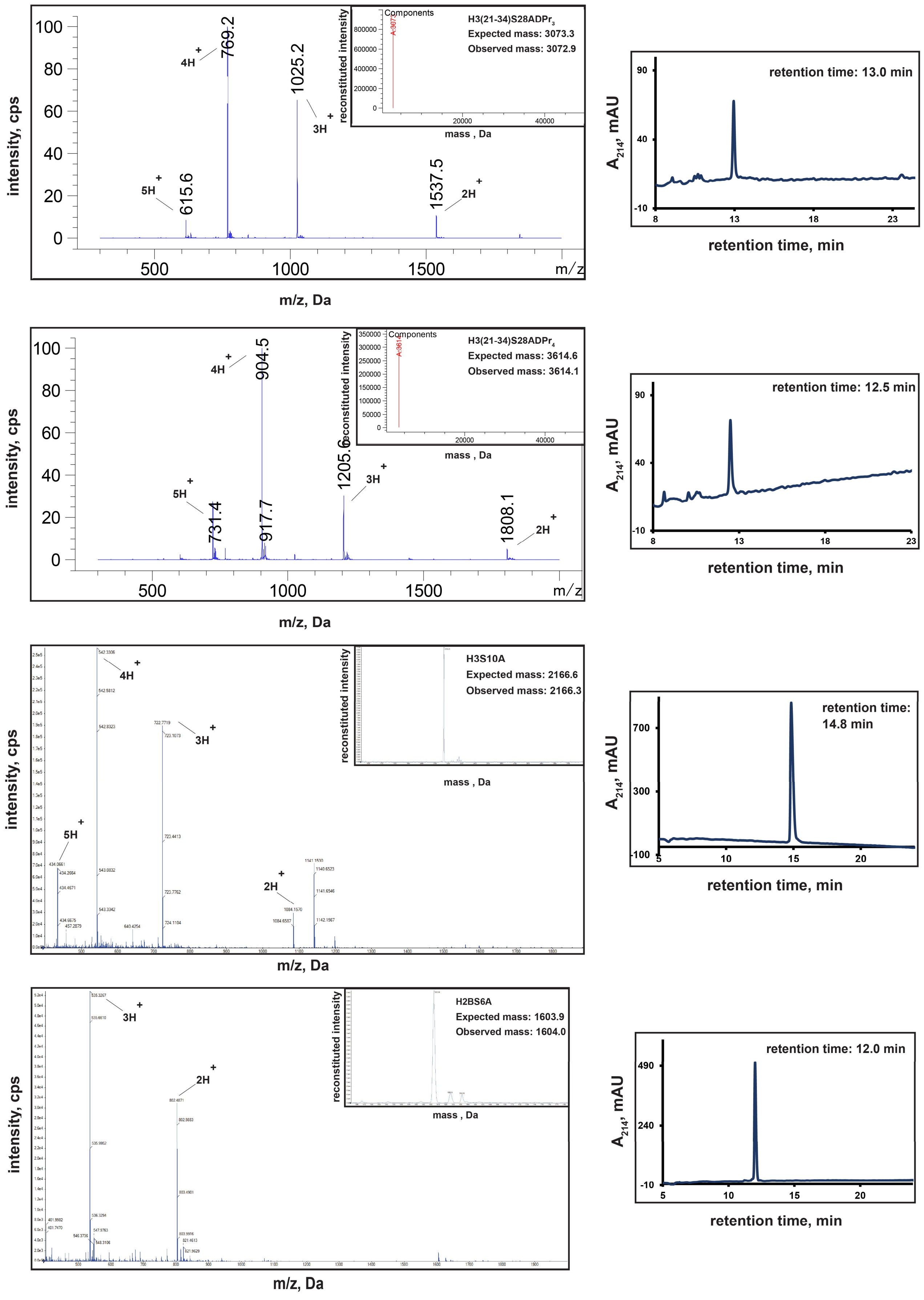
intensity, cps
m/z, Da
reconstituted intensity
mass , Da
H3(21-34)S28ADPr$_3$
Expected mass: 3073.3
Observed mass: 3072.9
retention time: 13.0 min
H3(21-34)S28ADPr$_4$
Expected mass: 3614.6
Observed mass: 3614.1
retention time: 12.5 min
H3S10A
Expected mass: 2166.6
Observed mass: 2166.3
retention time: 14.8 min
H2BS6A
Expected mass: 1603.9
Observed mass: 1604.0
retention time: 12.0 min
A$_{214}$, mAU
retention time, min

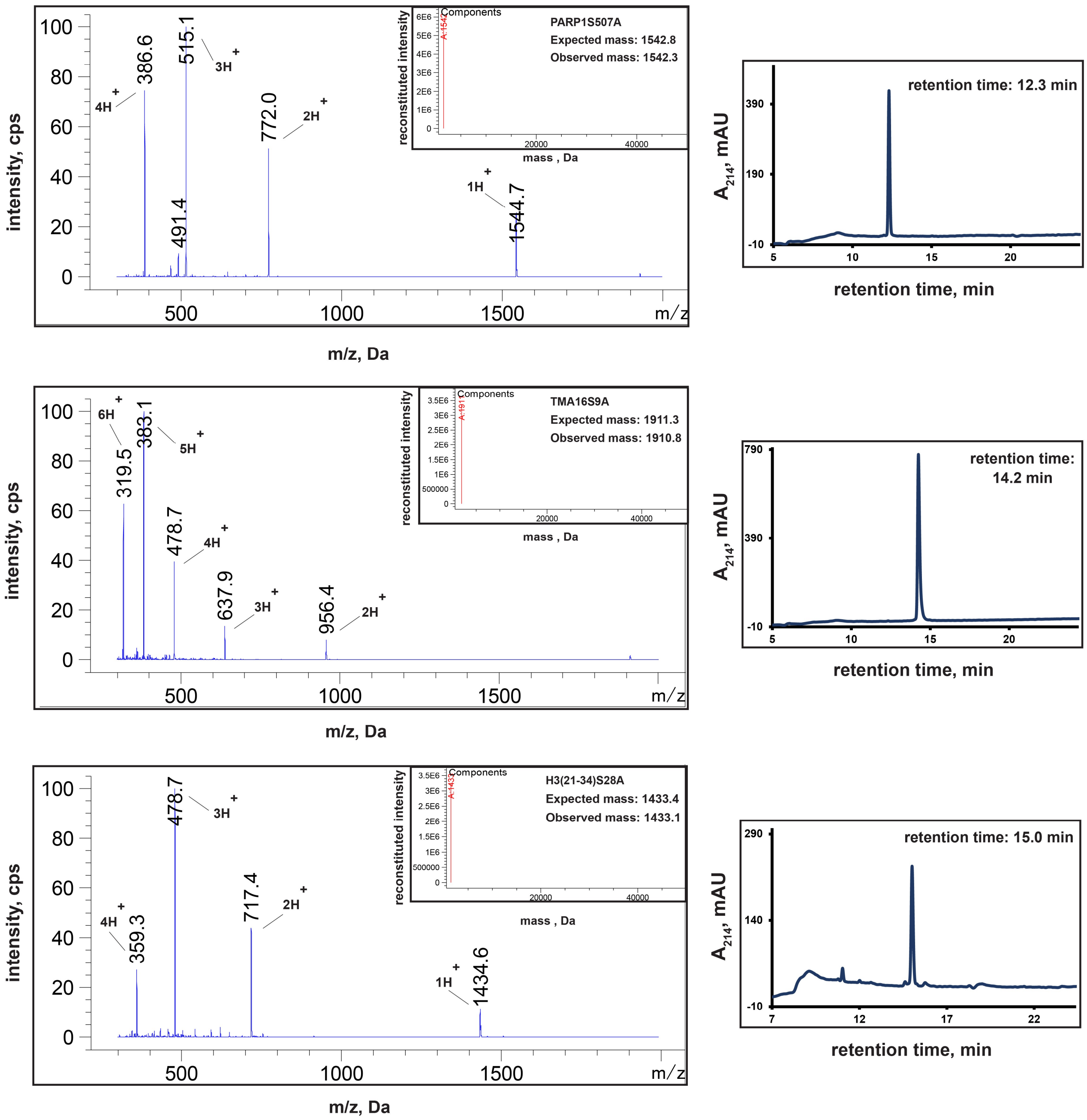
PARP1S507A
Expected mass: 1542.8
Observed mass: 1542.3
Components
reconstituted intensity
mass , Da
intensity, cps
m/z, Da
386.6
515.1
772.0
491.4
1544.7
4H+
3H+
2H+
1H+
retention time: 12.3 min
$A_{214}$, mAU
retention time, min
TMA16S9A
Expected mass: 1911.3
Observed mass: 1910.8
319.5
383.1
478.7
637.9
956.4
6H+
5H+
4H+
3H+
2H+
retention time: 14.2 min
H3(21-34)S28A
Expected mass: 1433.4
Observed mass: 1433.1
359.3
478.7
717.4
1434.6
4H+
3H+
2H+
1H+
retention time: 15.0 min

**b**

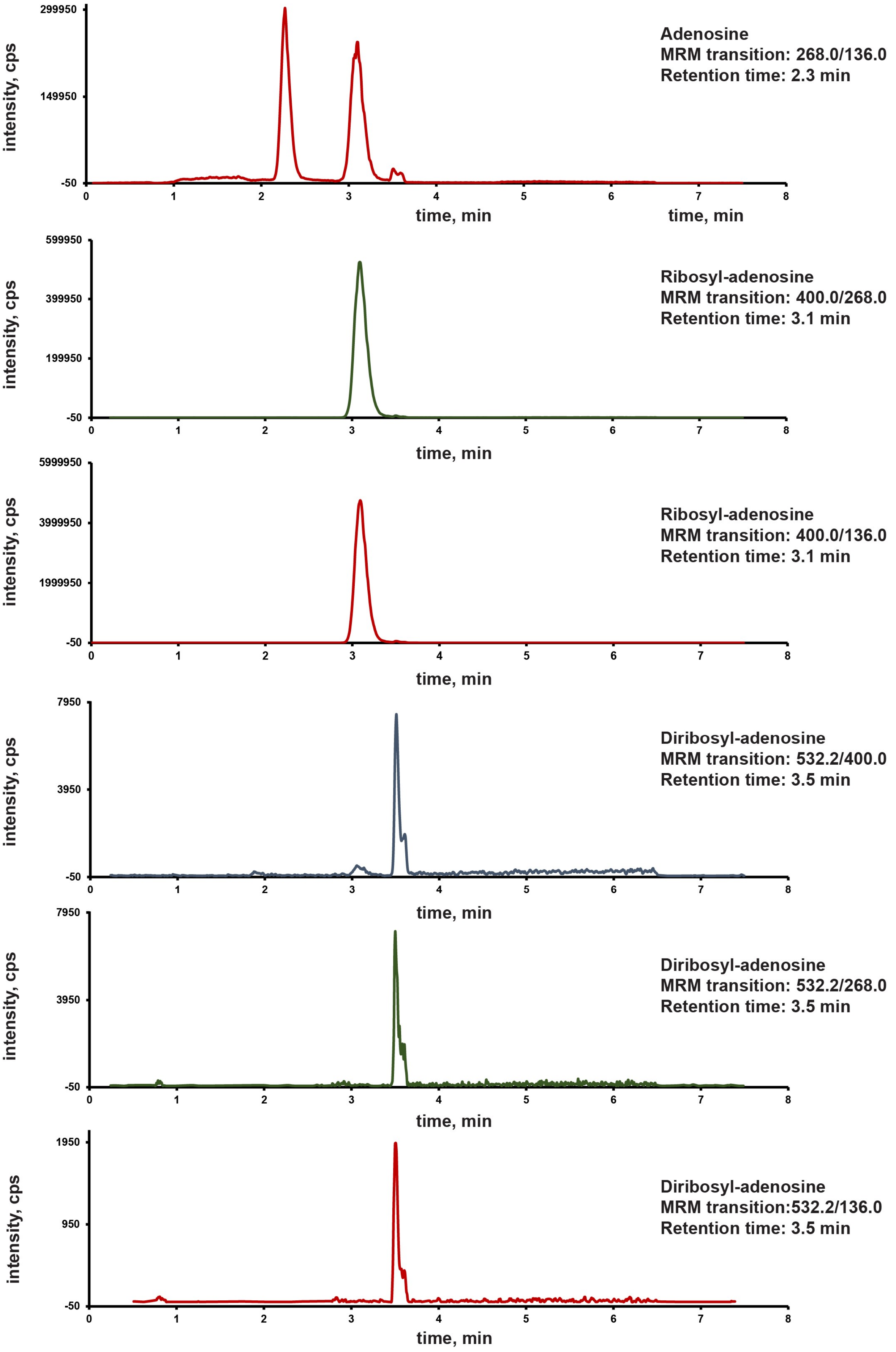

299950
149950
-50
0
1
2
3
4
5
6
7
8
intensity, cps
time, min
time, min
Adenosine
MRM transition: 268.0/136.0
Retention time: 2.3 min
599950
399950
199950
-50
intensity, cps
time, min
Ribosyl-adenosine
MRM transition: 400.0/268.0
Retention time: 3.1 min
5999950
3999950
1999950
-50
intensity, cps
time, min
Ribosyl-adenosine
MRM transition: 400.0/136.0
Retention time: 3.1 min
7950
3950
-50
intensity, cps
time, min
Diribosyl-adenosine
MRM transition: 532.2/400.0
Retention time: 3.5 min
7950
3950
-50
intensity, cps
time, min
Diribosyl-adenosine
MRM transition: 532.2/268.0
Retention time: 3.5 min
1950
950
-50
intensity, cps
time, min
Diribosyl-adenosine
MRM transition:532.2/136.0
Retention time: 3.5 min

| ANALYTE | MRM TRANSITION (Parent/daughter, Da) | INTEGRATED PEAK AREA |
|---|---|---|
| Adenosine | 268.0/136.0 | 2160000 |
| Ribosyl-adenosine | 400.0/136.0 | 45800000 |
| Diribosyl-adenosine | 532.2/136.0 | 7870 |

$$\text{Branching frequency} = \frac{\text{(peak area of diribosyl-adenosine)}}{\text{(peak area of ribosyl-adenosine)}} = 1.7 \times 10^{-4} \text{ OR } 0.017\%$$

**c**

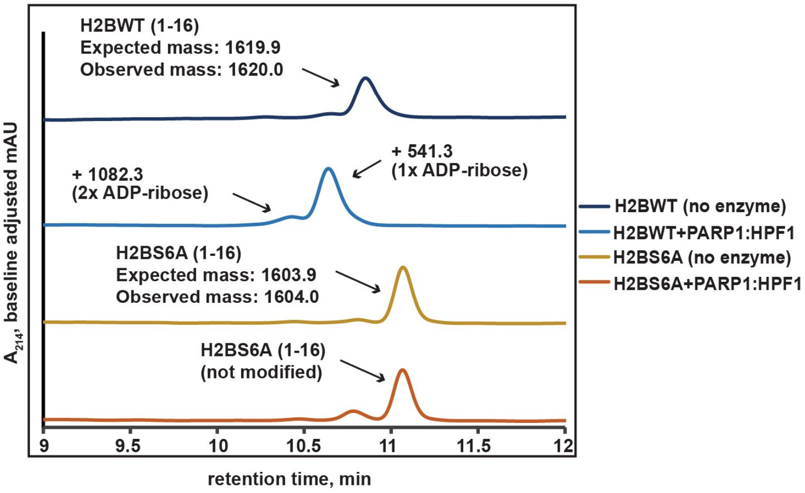


**d**

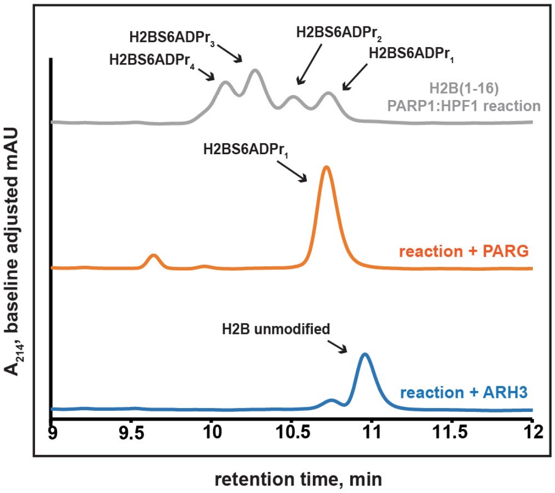


**e**

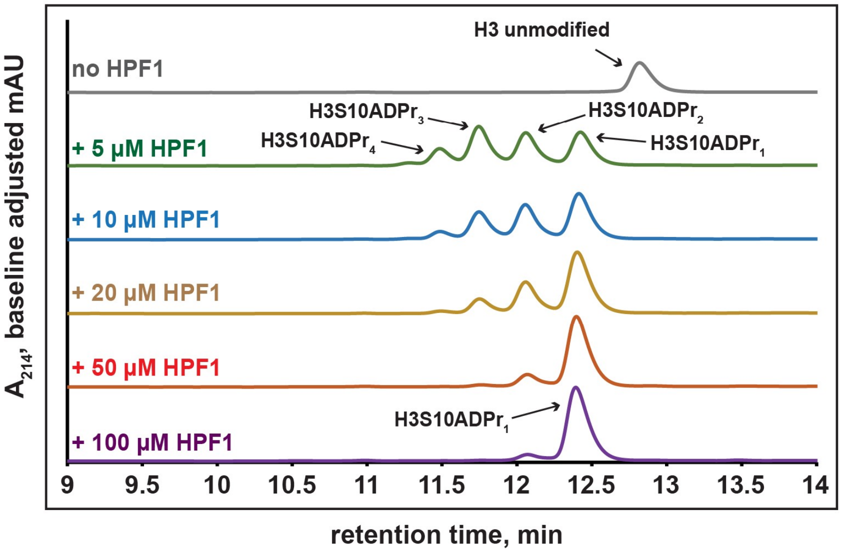

**Supplementary Fig. 1: Site-specific ADP-ribosylation of peptides from PARP1:HPF1 substrates and RP-HPLC-MS characterization of all peptides described here and in Fig. 1 and 2**
**a**, RP-HPLC and MS (raw and deconvoluted spectra) characterization of all purified, ADP-ribosylated H3 (amino acids 1-20), H2B (amino acids 1-16), PARP1 (amino acids 501-515), TMA16 (amino acids 2-19) and H3(amino acids 21-34, C-term thiazolidine, N-term SEA) peptides described here and in Fig. 1 and 2. All RP-HPLC gradients are from 0-35% Solvent B (2-22 min), except for those for H3S10A and H2BS6A, which are from 0-30% Solvent B (2-22 min). All H3 and H2B peptide MS spectra were produced on a Sciex X500B QTOF, while the rest were produced on an Agilent LC/MSD. **b**, LC-MS/MS analysis of enzymatic digestion products from PAR chains installed on a peptide using our technology. The spectra were produced on a Sciex QTRAP 6500+ mass spectrometer. **c,** RP-HPLC and MS analysis of substrate peptides (H2B wild-type or S6A mutant, amino acids 1-16) and corresponding PARP1:HPF1 reaction products. RP-HPLC gradients are from 0-35% Solvent B (2-22 min). **d**, RP-HPLC traces from PARG- or ARH3-treated H2B peptide ADPr reactions that were optimized for ADP-ribose chain elongation. The number of ADP-ribose units was verified by MS analysis. **e,** Representative RP-HPLC analyses of ADPr reactions from Fig. 1e containing PARP1 (1 $\mu$M), unmodified H3 substrate peptide (amino acids 1-20; 180 $\mu$M), $NAD^+$ (2 mM), and various HPF1 concentrations (indicated on corresponding trace). RP-HPLC gradients are from 0-35% Solvent B (2-22 min). For peak area integration values, see Supplementary Dataset.

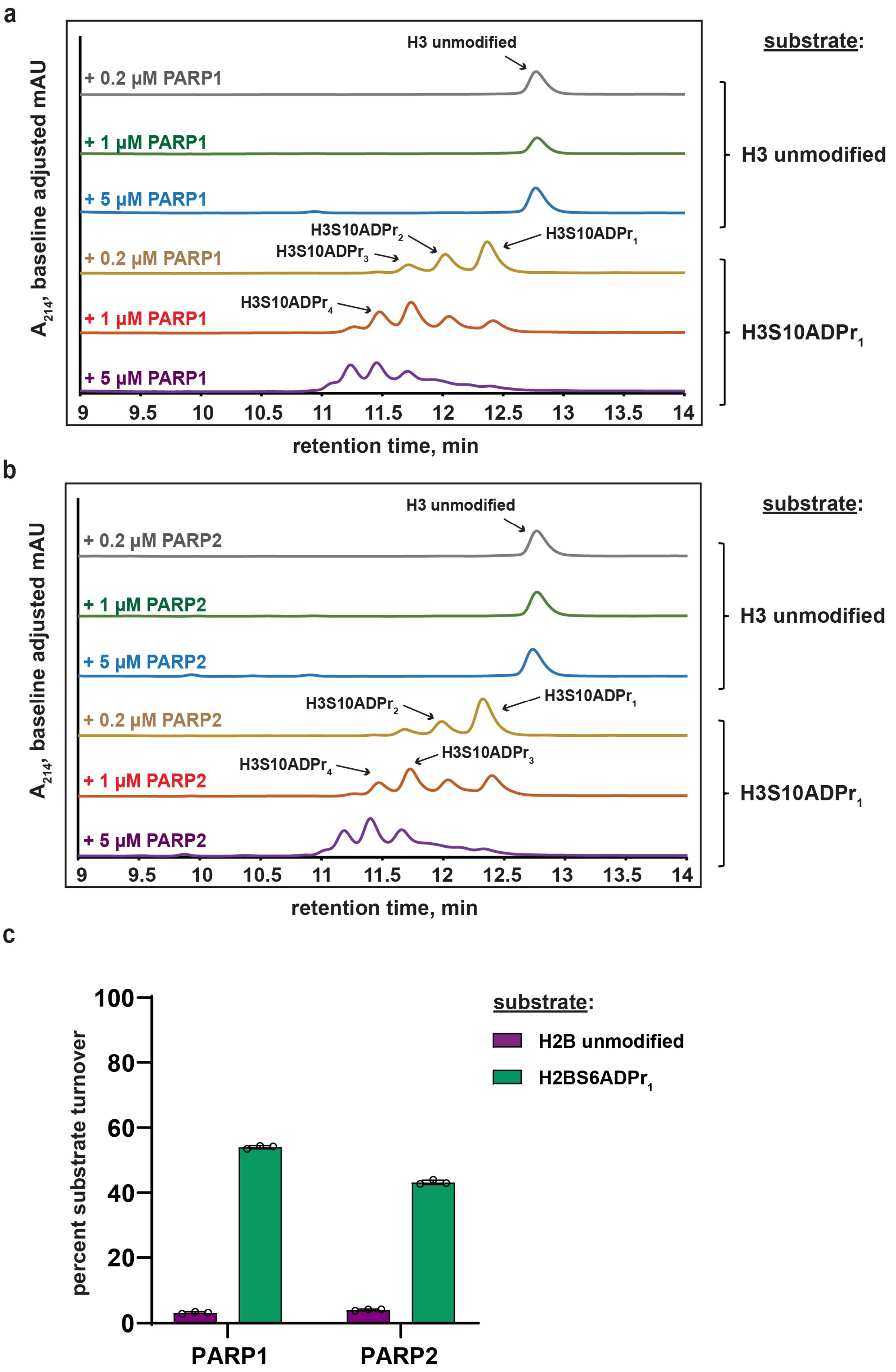
a
H3 unmodified
substrate:
+ 0.2 µM PARP1
+ 1 µM PARP1
+ 5 µM PARP1
H3 unmodified
H3S10ADPr2
H3S10ADPr3
H3S10ADPr1
+ 0.2 µM PARP1
H3S10ADPr4
+ 1 µM PARP1
H3S10ADPr1
+ 5 µM PARP1
A214, baseline adjusted mAU
9 9.5 10 10.5 11 11.5 12 12.5 13 13.5 14
retention time, min
b
H3 unmodified
substrate:
+ 0.2 µM PARP2
+ 1 µM PARP2
H3 unmodified
+ 5 µM PARP2
H3S10ADPr2
H3S10ADPr1
+ 0.2 µM PARP2
H3S10ADPr3
H3S10ADPr4
+ 1 µM PARP2
H3S10ADPr1
+ 5 µM PARP2
A214, baseline adjusted mAU
9 9.5 10 10.5 11 11.5 12 12.5 13 13.5 14
retention time, min
c
percent substrate turnover
100
80
60
40
20
0
substrate:
H2B unmodified
H2BS6ADPr1
PARP1
PARP2

d

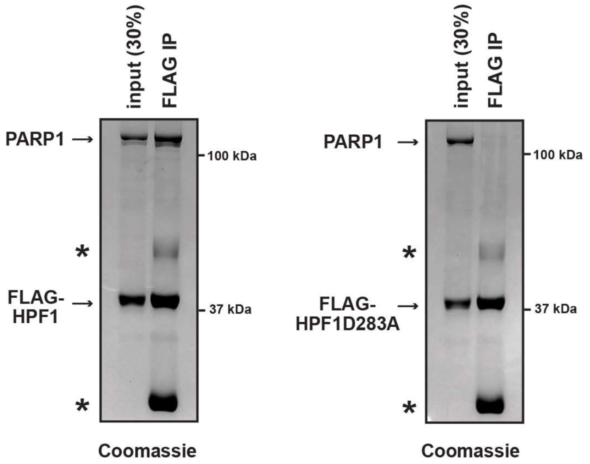
input (30%)
FLAG IP
PARP1
100 kDa
FLAG-HPF1
37 kDa
*
*
Coomassie
input (30%)
FLAG IP
PARP1
100 kDa
FLAG-HPF1D283A
37 kDa
*
*
Coomassie

e

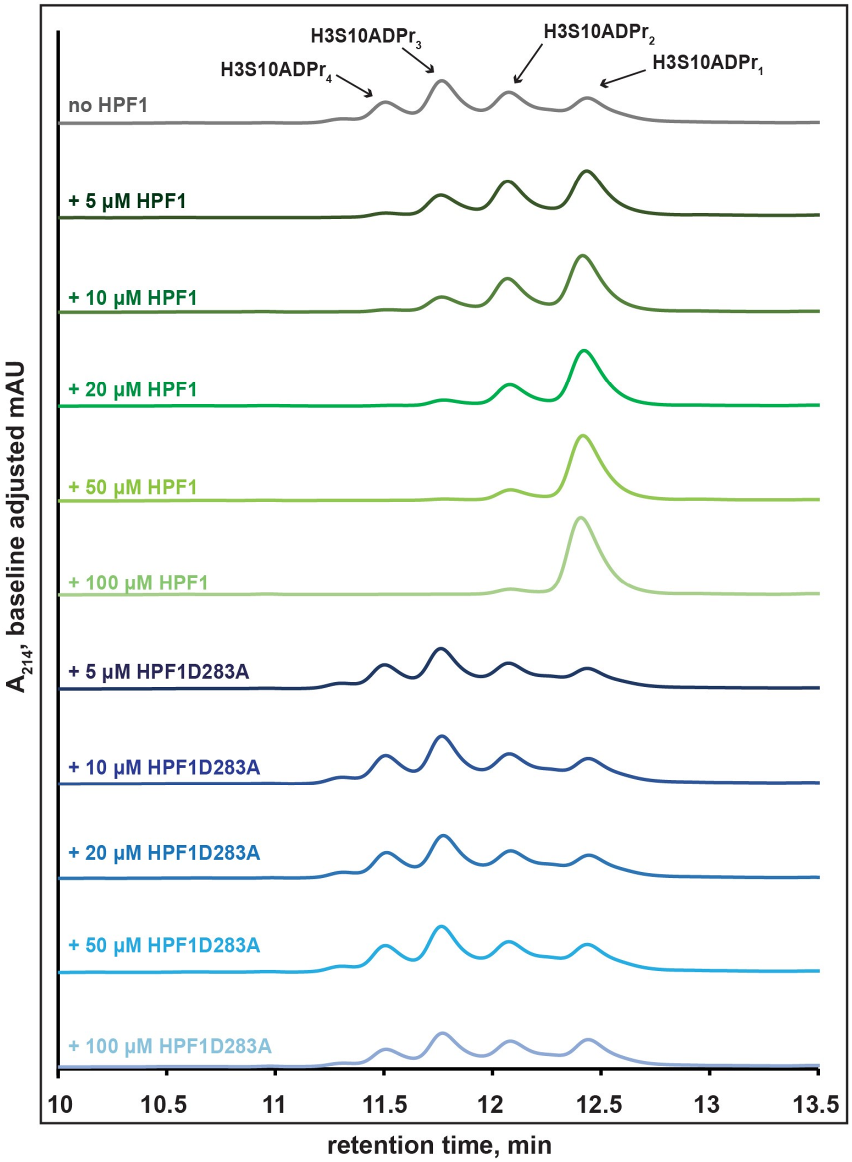
H3S10ADPr4
H3S10ADPr3
H3S10ADPr2
H3S10ADPr1
no HPF1
+ 5 µM HPF1
+ 10 µM HPF1
+ 20 µM HPF1
+ 50 µM HPF1
+ 100 µM HPF1
+ 5 µM HPF1D283A
+ 10 µM HPF1D283A
+ 20 µM HPF1D283A
+ 50 µM HPF1D283A
+ 100 µM HPF1D283A
A214, baseline adjusted mAU
10
10.5
11
11.5
12
12.5
13
13.5
retention time, min

f

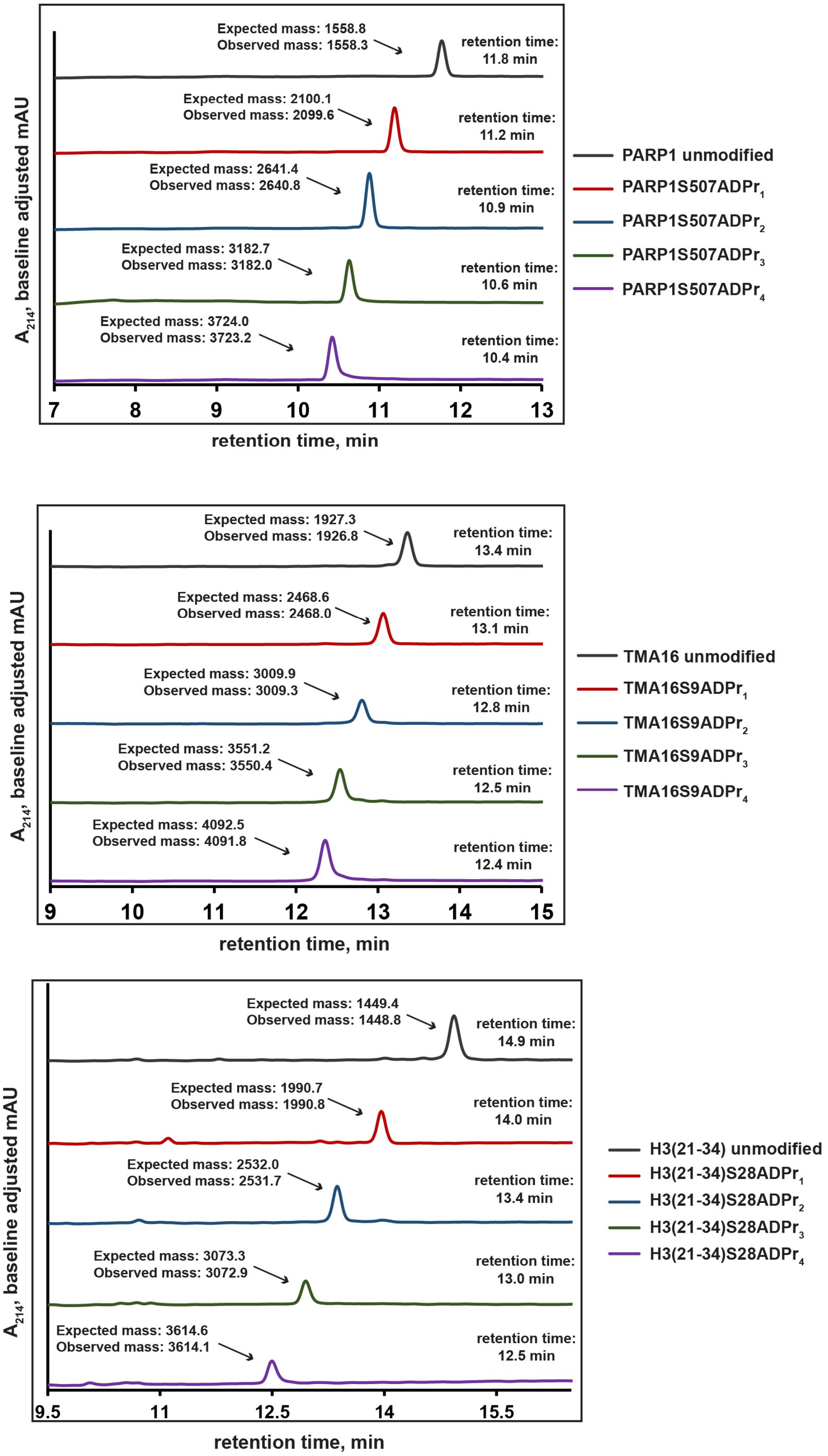
Expected mass: 1558.8
Observed mass: 1558.3
retention time: 11.8 min
Expected mass: 2100.1
Observed mass: 2099.6
retention time: 11.2 min
Expected mass: 2641.4
Observed mass: 2640.8
retention time: 10.9 min
Expected mass: 3182.7
Observed mass: 3182.0
retention time: 10.6 min
Expected mass: 3724.0
Observed mass: 3723.2
retention time: 10.4 min
$A_{214}$, baseline adjusted mAU
7 8 9 10 11 12 13
retention time, min
PARP1 unmodified
PARP1S507ADPr$_1$
PARP1S507ADPr$_2$
PARP1S507ADPr$_3$
PARP1S507ADPr$_4$
Expected mass: 1927.3
Observed mass: 1926.8
retention time: 13.4 min
Expected mass: 2468.6
Observed mass: 2468.0
retention time: 13.1 min
Expected mass: 3009.9
Observed mass: 3009.3
retention time: 12.8 min
Expected mass: 3551.2
Observed mass: 3550.4
retention time: 12.5 min
Expected mass: 4092.5
Observed mass: 4091.8
retention time: 12.4 min
$A_{214}$, baseline adjusted mAU
9 10 11 12 13 14 15
retention time, min
TMA16 unmodified
TMA16S9ADPr$_1$
TMA16S9ADPr$_2$
TMA16S9ADPr$_3$
TMA16S9ADPr$_4$
Expected mass: 1449.4
Observed mass: 1448.8
retention time: 14.9 min
Expected mass: 1990.7
Observed mass: 1990.8
retention time: 14.0 min
Expected mass: 2532.0
Observed mass: 2531.7
retention time: 13.4 min
Expected mass: 3073.3
Observed mass: 3072.9
retention time: 13.0 min
Expected mass: 3614.6
Observed mass: 3614.1
retention time: 12.5 min
$A_{214}$, baseline adjusted mAU
9.5 11 12.5 14 15.5
retention time, min
H3(21-34) unmodified
H3(21-34)S28ADPr$_1$
H3(21-34)S28ADPr$_2$
H3(21-34)S28ADPr$_3$
H3(21-34)S28ADPr$_4$

**g**

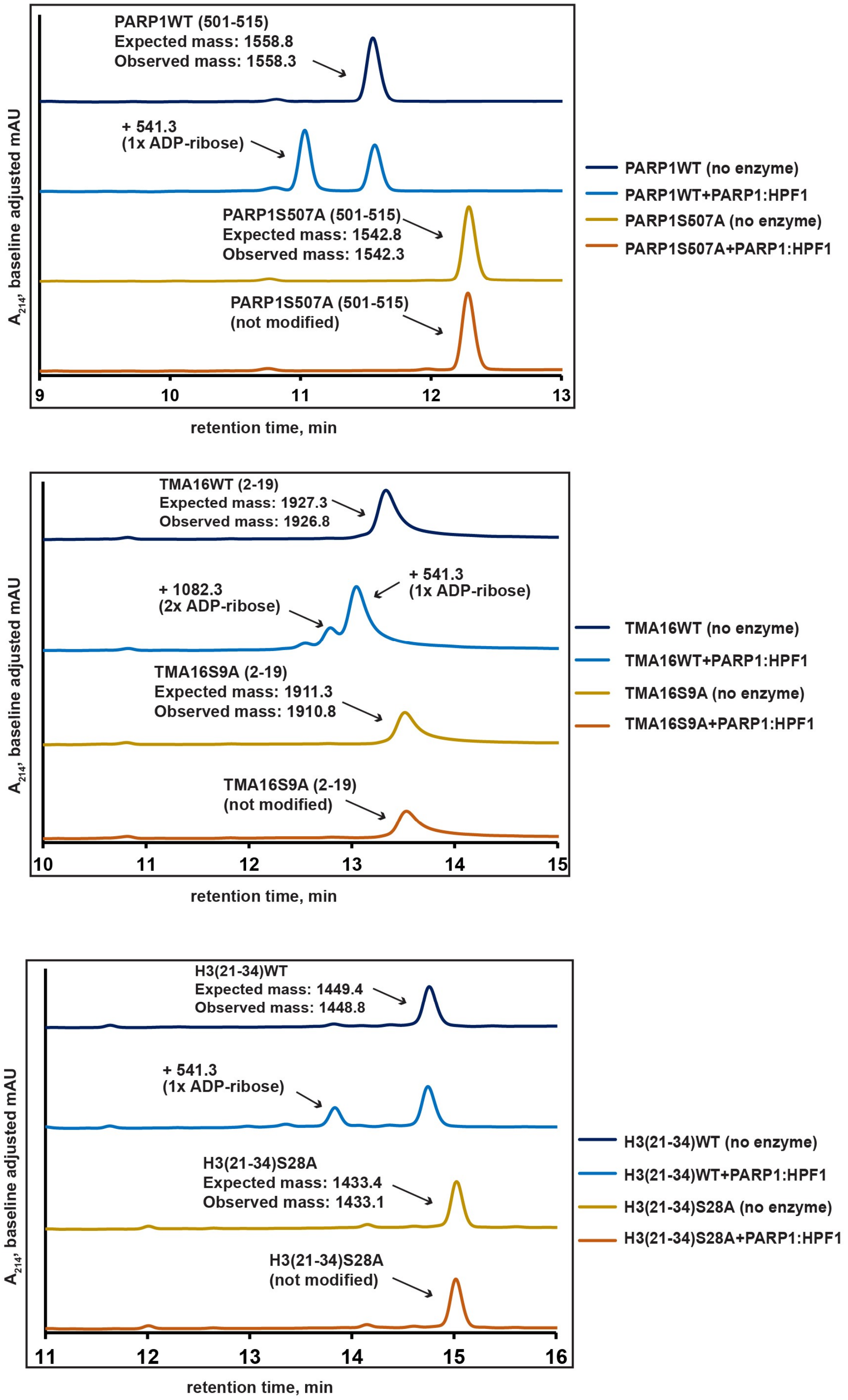
PARP1WT (501-515)
Expected mass: 1558.8
Observed mass: 1558.3
+ 541.3
(1x ADP-ribose)
PARP1S507A (501-515)
Expected mass: 1542.8
Observed mass: 1542.3
PARP1S507A (501-515)
(not modified)
A214, baseline adjusted mAU
9
10
11
12
13
retention time, min
PARP1WT (no enzyme)
PARP1WT+PARP1:HPF1
PARP1S507A (no enzyme)
PARP1S507A+PARP1:HPF1
TMA16WT (2-19)
Expected mass: 1927.3
Observed mass: 1926.8
+ 1082.3
(2x ADP-ribose)
+ 541.3
(1x ADP-ribose)
TMA16S9A (2-19)
Expected mass: 1911.3
Observed mass: 1910.8
TMA16S9A (2-19)
(not modified)
A214, baseline adjusted mAU
10
11
12
13
14
15
retention time, min
TMA16WT (no enzyme)
TMA16WT+PARP1:HPF1
TMA16S9A (no enzyme)
TMA16S9A+PARP1:HPF1
H3(21-34)WT
Expected mass: 1449.4
Observed mass: 1448.8
+ 541.3
(1x ADP-ribose)
H3(21-34)S28A
Expected mass: 1433.4
Observed mass: 1433.1
H3(21-34)S28A
(not modified)
A214, baseline adjusted mAU
11
12
13
14
15
16
retention time, min
H3(21-34)WT (no enzyme)
H3(21-34)WT+PARP1:HPF1
H3(21-34)S28A (no enzyme)
H3(21-34)S28A+PARP1:HPF1

**Supplementary Fig. 2: Employing a two-step enzymatic process to prepare poly-ADP-ribosylated peptides on a large scale**

**a**, Representative RP-HPLC analyses of ADPr reactions from Fig. 2b containing various PARP1 concentrations (indicated on corresponding trace), unmodified H3 or H3S10ADPr$_1$ substrate peptide (amino acids 1-20; 180 μM), $NAD^+$ (2 mM), and no HPF1. RP-HPLC gradients are from 0-35% Solvent B (2-22 min). For peak area integration values, see Supplementary Dataset. **b**, As in **a**, but with various PARP2 concentrations (indicated on corresponding trace). **c**, Substrate turnover analysis of ADPr reactions containing PARP1 (1 μM) or PARP2 (1 μM), the indicated H2B substrate peptide (amino acids 1-16; 40 μM), $NAD^+$ (2 mM), and no HPF1. Purple bars represent total percent turnover of an unmodified H2B peptide to mono- or poly-ADP-ribosylated products. Green bars represent total percent turnover of the H2BS6ADPr$_1$ peptide to poly-ADP-ribosylated products. For peak area integration values, see Supplementary Dataset. Data are represented as mean ± s.d. (n = 3). **d,** Coomassie blue-stained SDS-PAGE gel showing FLAG-HPF1:PARP1 or FLAG-HPF1D283A:PARP1 immunoprecipitation (IP) experiments. *Gel migration species corresponding to the immunoglobulin chains from the anti-FLAG M2 magnetic beads**.** **e**, Representative RP-HPLC analyses of ADPr reactions from Fig. 2c containing PARP1 (1 μM), H3S10ADPr$_1$ substrate peptide (amino acids 1-20; 180 μM), $NAD^+$ (2 mM), and various HPF1 or HPF1D283A concentrations (concentration indicated on corresponding trace). RP-HPLC gradients are from 0-35% Solvent B (2-22 min). For peak area integration values, see Supplementary Dataset. **f,** RP-HPLC and MS analysis of mono- and poly-ADP-ribosylated PARP1, TMA16 and H3(21-34) peptides that have been puridied to homogeneity using semi-preparative HPLC. RP-HPLC gradients are from 0-35% Solvent B (2-22 min) **g,** RP-HPLC and MS analysis of substrate peptides (PARP1 wild-type or S507A mutant, TMA16 wild-type or S9A mutant, H3(21-34) wild-type of S28A mutant) and corresponding PARP1:HPF1 reaction products. RP-HPLC gradients are from 0-35% Solvent B (2-22 min).

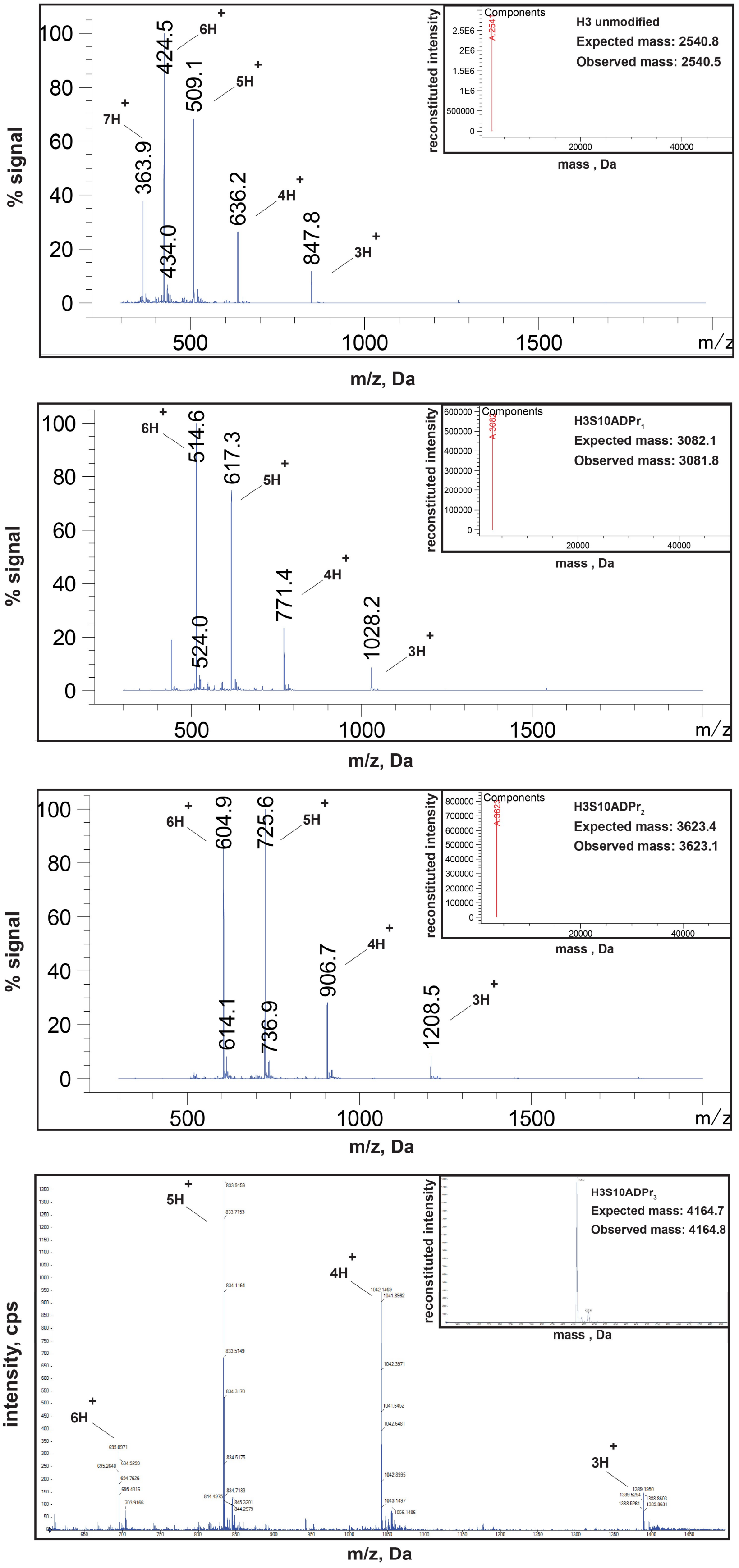
% signal
m/z, Da
6H+
424.5
5H+
509.1
7H+
363.9
4H+
636.2
3H+
847.8
434.0
reconstituted intensity
Components
H3 unmodified
Expected mass: 2540.8
Observed mass: 2540.5
mass , Da
6H+
514.6
617.3
5H+
4H+
771.4
3H+
1028.2
524.0
H3S10ADPr1
Expected mass: 3082.1
Observed mass: 3081.8
6H+
604.9
725.6
5H+
4H+
906.7
3H+
1208.5
614.1
736.9
H3S10ADPr2
Expected mass: 3623.4
Observed mass: 3623.1
intensity, cps
H3S10ADPr3
Expected mass: 4164.7
Observed mass: 4164.8

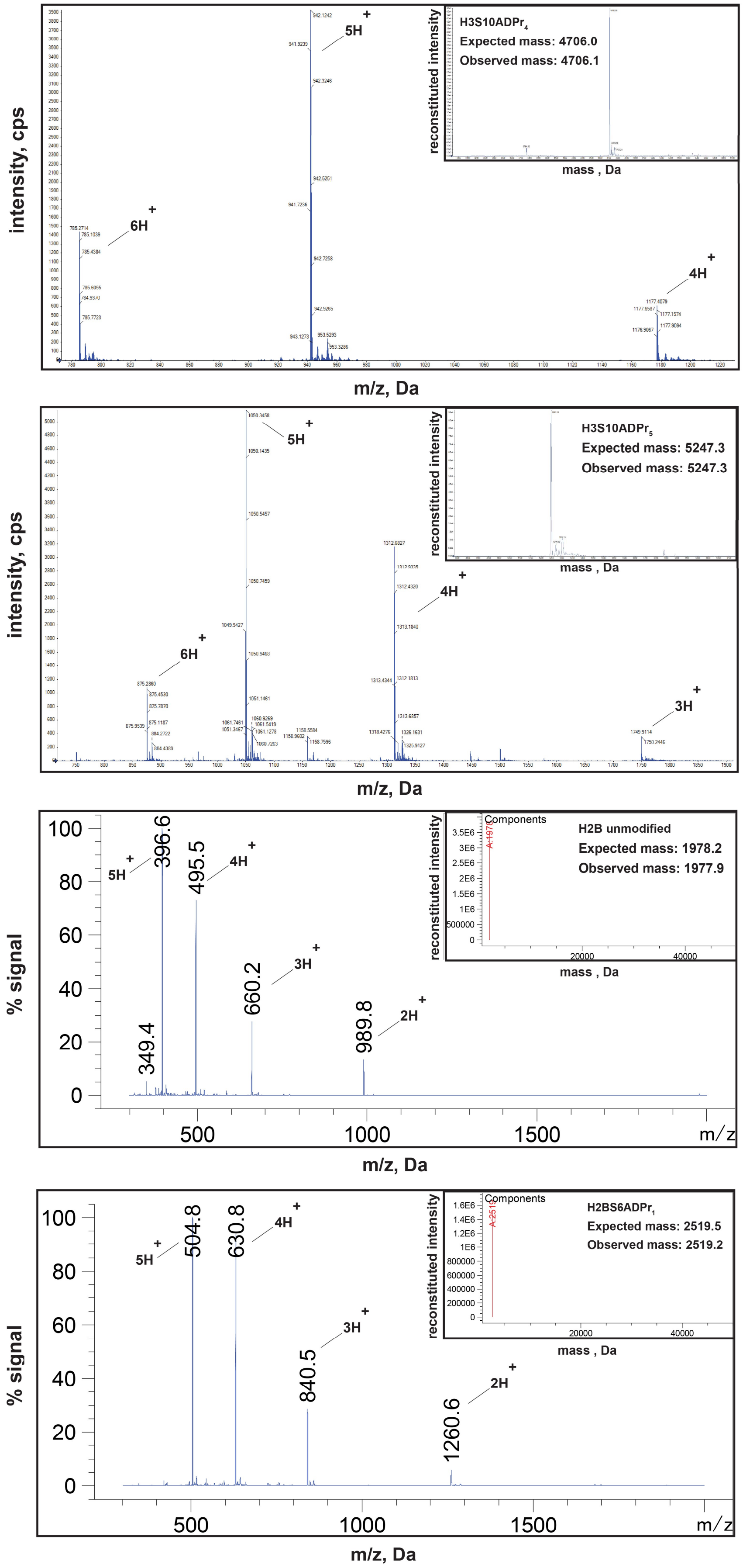
intensity, cps
m/z, Da
reconstituted intensity
mass , Da
H3S10ADPr4
Expected mass: 4706.0
Observed mass: 4706.1
5H+
6H+
4H+
H3S10ADPr5
Expected mass: 5247.3
Observed mass: 5247.3
3H+
% signal
Components
H2B unmodified
Expected mass: 1978.2
Observed mass: 1977.9
396.6
495.5
660.2
989.8
349.4
2H+
H2BS6ADPr1
Expected mass: 2519.5
Observed mass: 2519.2
504.8
630.8
840.5
1260.6
m/z

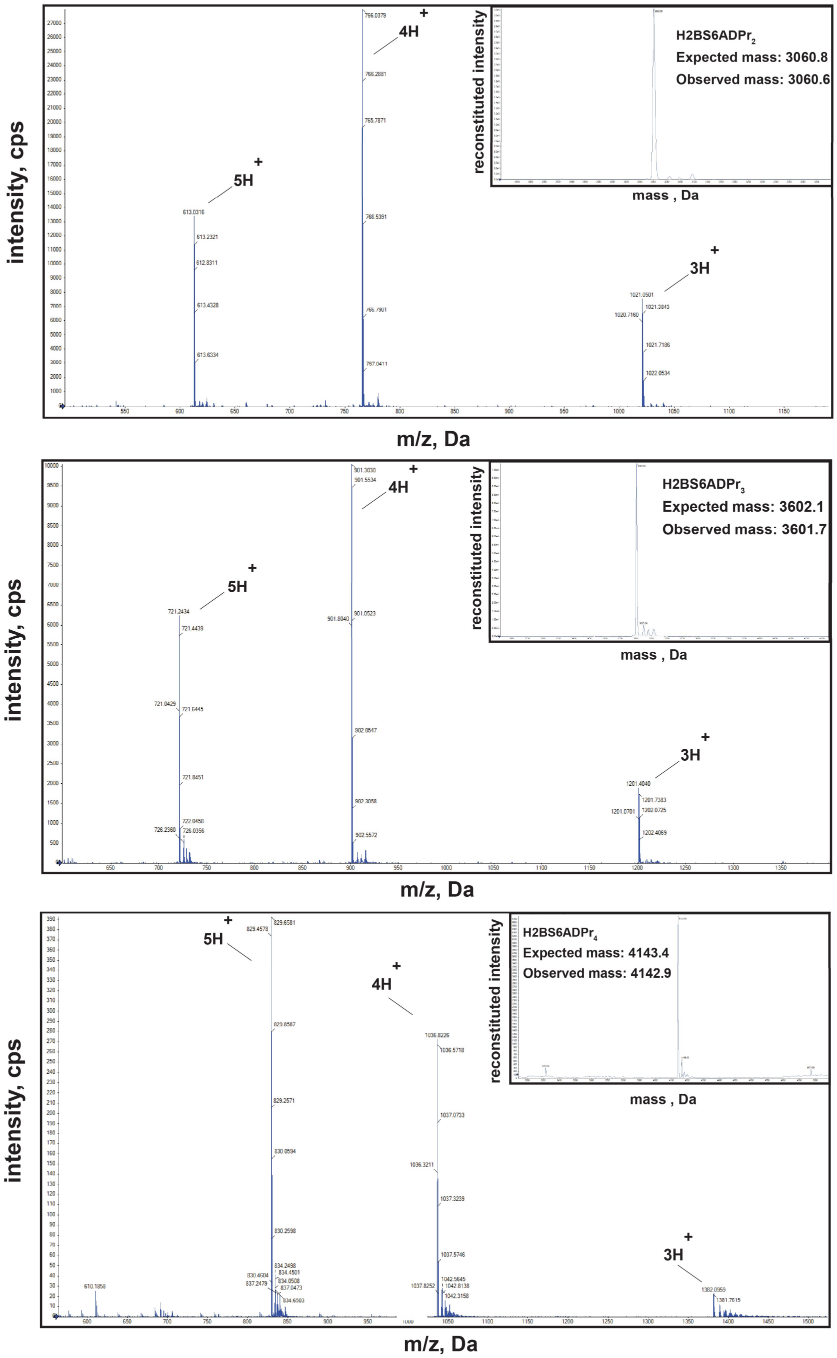
intensity, cps
4H+
5H+
3H+
reconstituted intensity
H2BS6ADPr$_2$
Expected mass: 3060.8
Observed mass: 3060.6
mass , Da
m/z, Da
intensity, cps
4H+
5H+
3H+
reconstituted intensity
H2BS6ADPr$_3$
Expected mass: 3602.1
Observed mass: 3601.7
mass , Da
m/z, Da
intensity, cps
5H+
4H+
3H+
reconstituted intensity
H2BS6ADPr$_4$
Expected mass: 4143.4
Observed mass: 4142.9
mass , Da
m/z, Da

**Supplementary Fig. 3: MS characterization of all fluorescein-labeled H3 (1-20) and H2B (1-16) peptides described in Fig. 3**
All MS spectra were produced on a Sciex X500B QTOF with the exception of H3 unmodified, H3S10ADPr$_1$, H3S10ADPr$_2$, H2B unmodified, and H2BS6ADPr$_1$, which were analyzed on an Agilent LC/MSD. Raw and deconvoluted spectra are shown.
*We note that peptides labeled with 5(6)-carboxyfluorescein typically exhibit peak splitting on RP-HPLC chromatograms due to the unique retention times of the separate enantiomers. We therefore relied upon whole-fraction MS analysis to characterize the identity and purity of purified, fluorescein-labeled molecules.

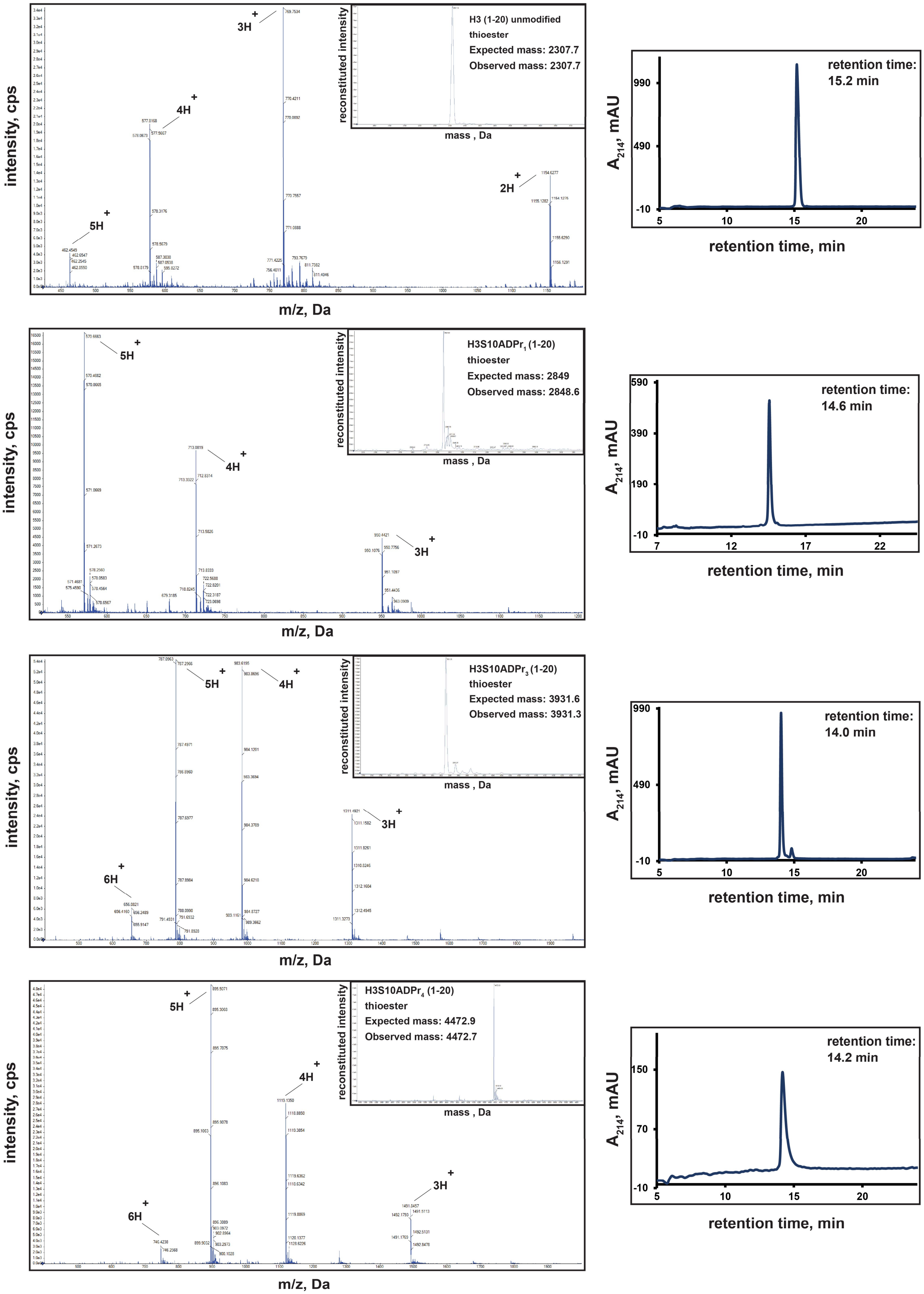
intensity, cps
m/z, Da
reconstituted intensity
mass , Da
H3 (1-20) unmodified
thioester
Expected mass: 2307.7
Observed mass: 2307.7
2H+
3H+
4H+
5H+
A214, mAU
retention time, min
retention time:
15.2 min
H3S10ADPr1 (1-20)
thioester
Expected mass: 2849
Observed mass: 2848.6
retention time:
14.6 min
H3S10ADPr3 (1-20)
thioester
Expected mass: 3931.6
Observed mass: 3931.3
6H+
retention time:
14.0 min
H3S10ADPr4 (1-20)
thioester
Expected mass: 4472.9
Observed mass: 4472.7
retention time:
14.2 min

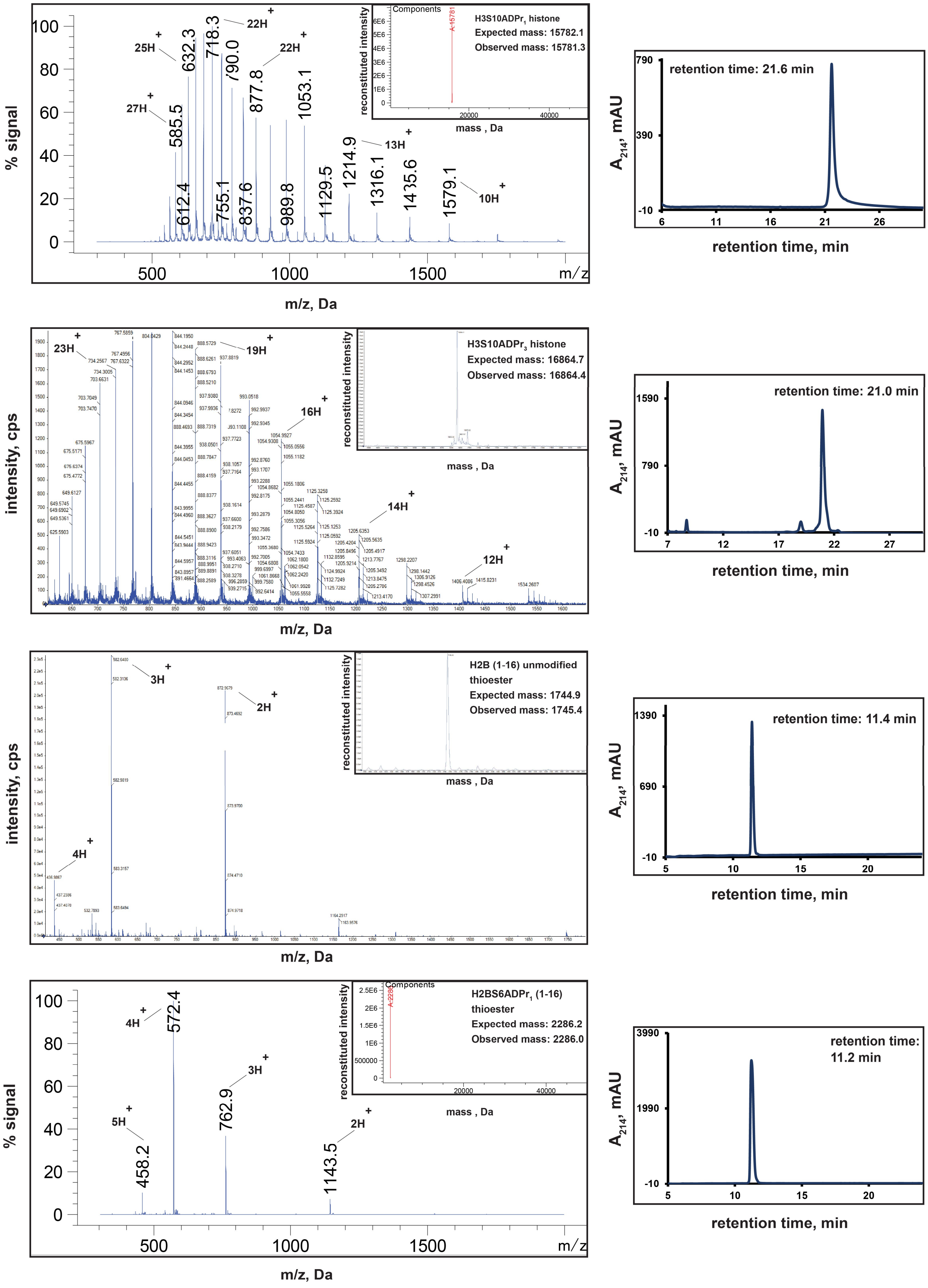
% signal
m/z, Da
25H+
27H+
22H+
13H+
10H+
585.5
612.4
632.3
718.3
755.1
790.0
837.6
877.8
989.8
1053.1
1129.5
1214.9
1316.1
1435.6
1579.1
Components
reconstituted intensity
mass , Da
H3S10ADPr1 histone
Expected mass: 15782.1
Observed mass: 15781.3
retention time: 21.6 min
A214, mAU
retention time, min
intensity, cps
23H+
19H+
16H+
14H+
12H+
H3S10ADPr3 histone
Expected mass: 16864.7
Observed mass: 16864.4
retention time: 21.0 min
3H+
2H+
4H+
H2B (1-16) unmodified thioester
Expected mass: 1744.9
Observed mass: 1745.4
retention time: 11.4 min
4H+
3H+
5H+
2H+
572.4
762.9
458.2
1143.5
H2BS6ADPr1 (1-16) thioester
Expected mass: 2286.2
Observed mass: 2286.0
retention time: 11.2 min

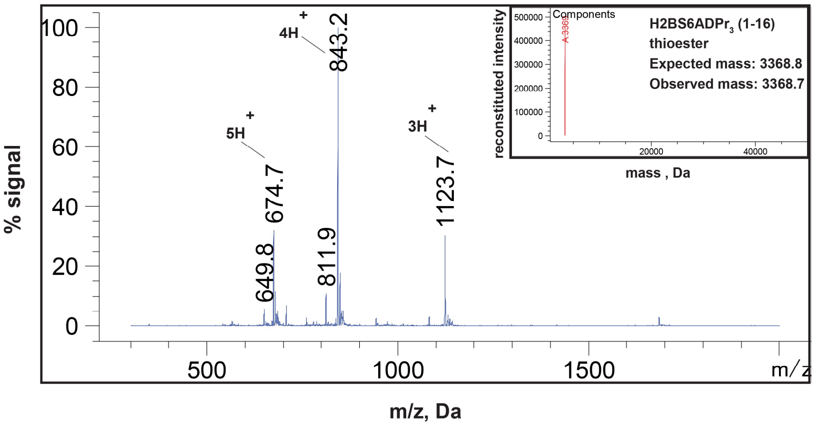

% signal
100
80
60
40
20
0
4H+
843.2
5H+
674.7
3H+
1123.7
649.8
811.9
500
1000
1500
m/z
m/z, Da
reconstituted intensity
Components
mass , Da
20000
40000
H2BS6ADPr$_3$ (1-16) thioester
Expected mass: 3368.8
Observed mass: 3368.7


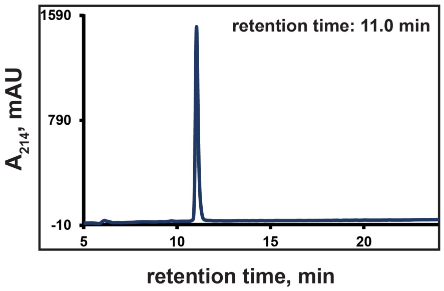

A$_{214}$, mAU
retention time: 11.0 min
1590
790
-10
5
10
15
20
retention time, min


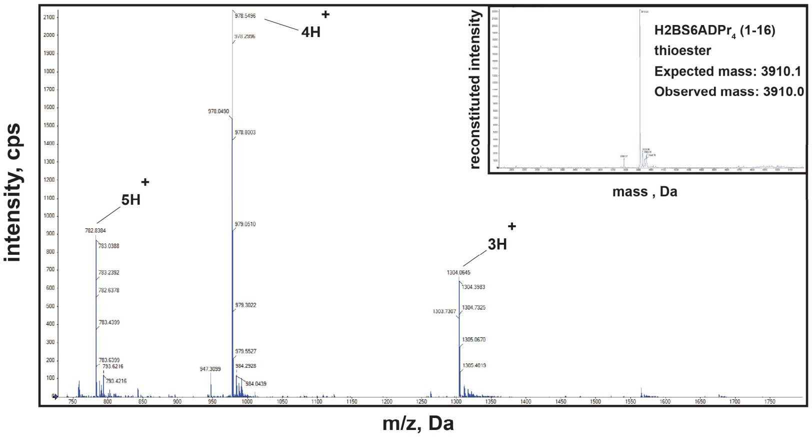

intensity, cps
4H+
5H+
3H+
m/z, Da
reconstituted intensity
mass , Da
H2BS6ADPr$_4$ (1-16) thioester
Expected mass: 3910.1
Observed mass: 3910.0


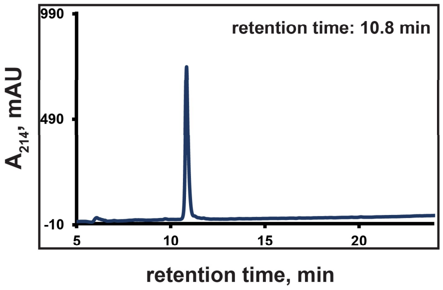

A$_{214}$, mAU
retention time: 10.8 min
990
490
-10
5
10
15
20
retention time, min


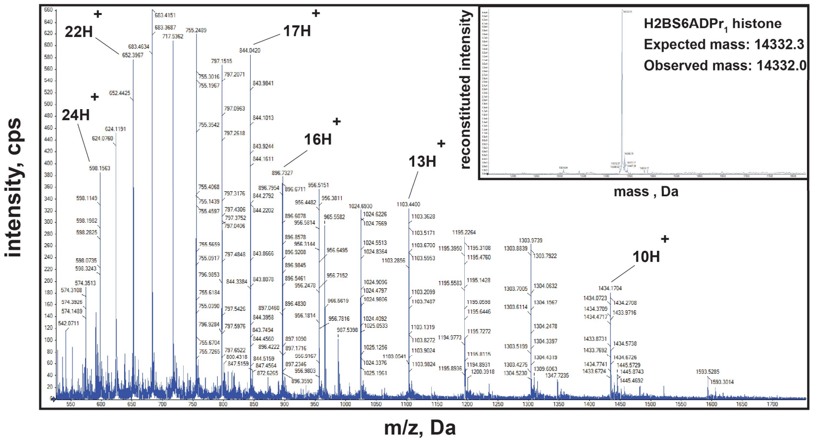

intensity, cps
22H+
24H+
17H+
16H+
13H+
10H+
m/z, Da
reconstituted intensity
mass , Da
H2BS6ADPr$_1$ histone
Expected mass: 14332.3
Observed mass: 14332.0


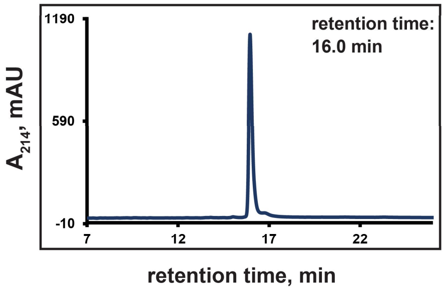

A$_{214}$, mAU
retention time: 16.0 min
1190
590
-10
7
12
17
22
retention time, min


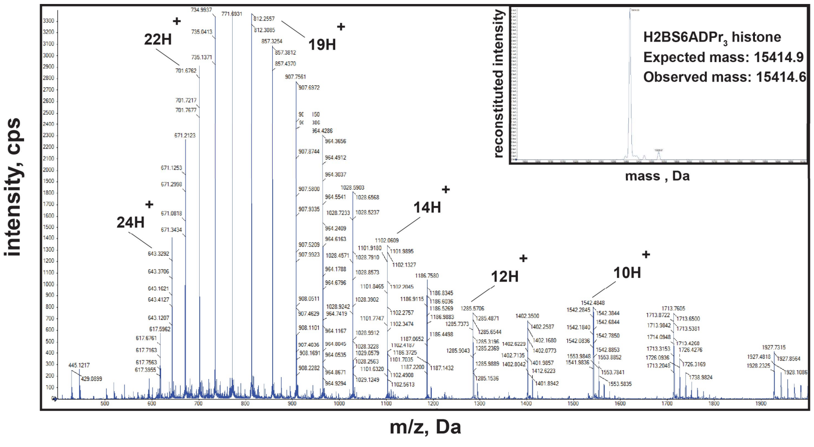

intensity, cps
22H+
19H+
24H+
14H+
12H+
10H+
m/z, Da
reconstituted intensity
mass , Da
H2BS6ADPr$_3$ histone
Expected mass: 15414.9
Observed mass: 15414.6


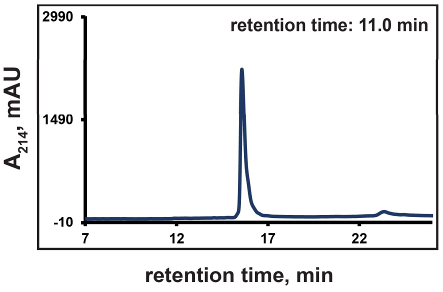

A$_{214}$, mAU
retention time: 11.0 min
2990
1490
-10
7
12
17
22
retention time, min

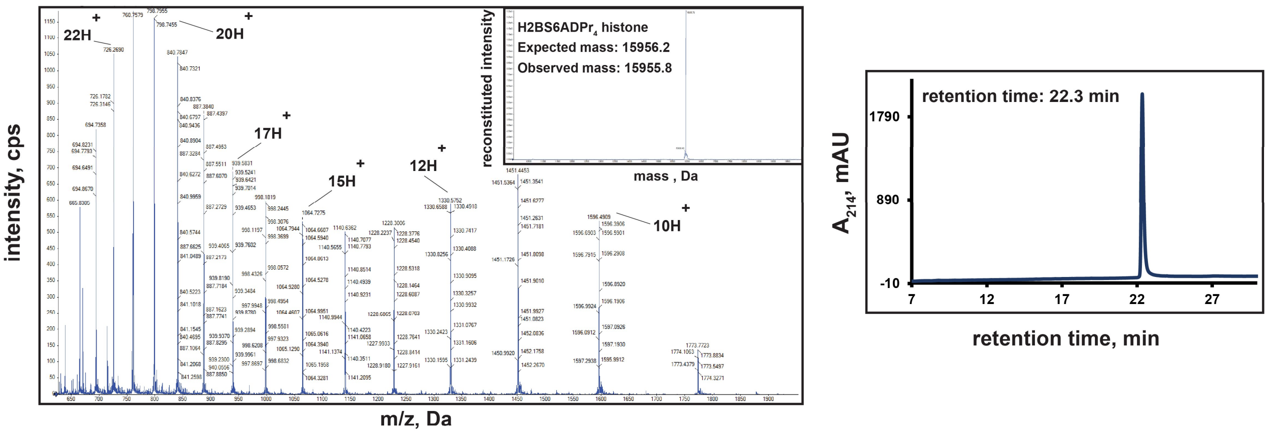


**Supplementary Fig. 4: RP-HPLC and MS characterization of full-length, ADP-ribosylated H3 and H2B histones and corresponding ADP-ribosylated thioester peptide building blocks**

For peptides: All MS spectra were produced on a Sciex X500B QTOF with the exception of $H2BS6ADPr_1$ and $H2BS6ADPr_3$, which were analyzed on an Agilent LC/MSD. Raw and deconvoluted spectra are shown. H2B (amino acids 1-16) and H3 (amino acids 1-20) were converted to thioesters via treatment with MESNa as described in Methods. RP-HPLC gradients are from 0-30% Solvent B (2-22 min).

For full-length proteins: All MS spectra were produced on a Sciex X500B QTOF with the exception of $H3S10ADPr_1$, which was analyzed on an Agilent LC/MSD. Raw and deconvoluted spectra are shown. All RP-HPLC gradients are from 30-80% Solvent B (3-33 min) except for that for $H2BS6ADPr_4$, which is from 0-80% Solvent B (2-22 min).

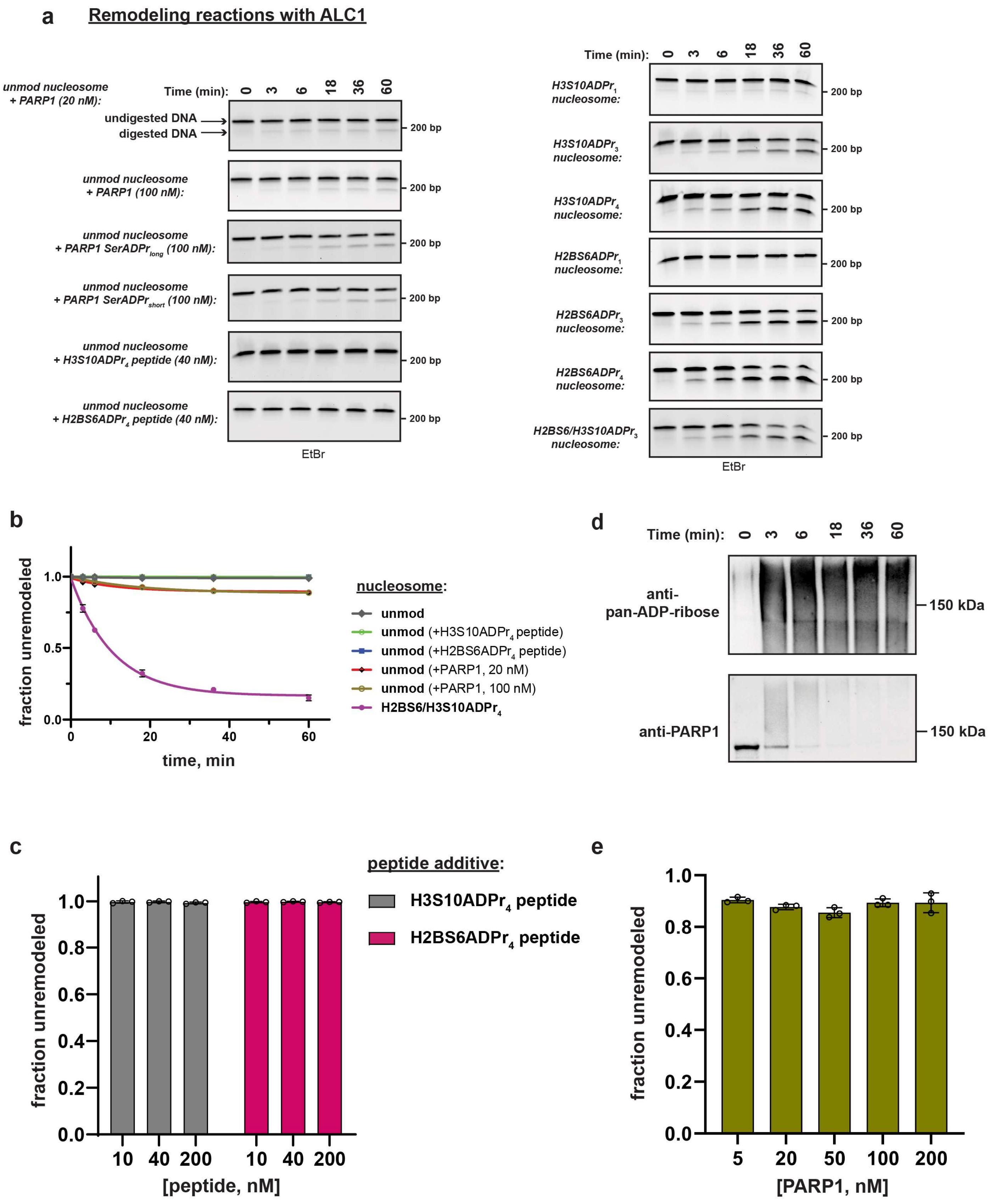
a
Remodeling reactions with ALC1
Time (min): 0 3 6 18 36 60
unmod nucleosome + PARP1 (20 nM):
undigested DNA
digested DNA
200 bp
unmod nucleosome + PARP1 (100 nM):
unmod nucleosome + PARP1 SerADPr$_{long}$ (100 nM):
unmod nucleosome + PARP1 SerADPr$_{short}$ (100 nM):
unmod nucleosome + H3S10ADPr$_4$ peptide (40 nM):
unmod nucleosome + H2BS6ADPr$_4$ peptide (40 nM):
EtBr
H3S10ADPr$_1$ nucleosome:
H3S10ADPr$_3$ nucleosome:
H3S10ADPr$_4$ nucleosome:
H2BS6ADPr$_1$ nucleosome:
H2BS6ADPr$_3$ nucleosome:
H2BS6ADPr$_4$ nucleosome:
H2BS6/H3S10ADPr$_3$ nucleosome:
b
fraction unremodeled
time, min
nucleosome:
unmod
unmod (+H3S10ADPr$_4$ peptide)
unmod (+H2BS6ADPr$_4$ peptide)
unmod (+PARP1, 20 nM)
unmod (+PARP1, 100 nM)
H2BS6/H3S10ADPr$_4$
d
Time (min): 0 3 6 18 36 60
anti-pan-ADP-ribose
anti-PARP1
150 kDa
c
peptide additive:
H3S10ADPr$_4$ peptide
H2BS6ADPr$_4$ peptide
fraction unremodeled
10 40 200 10 40 200
[peptide, nM]
e
fraction unremodeled
5 20 50 100 200
[PARP1, nM]

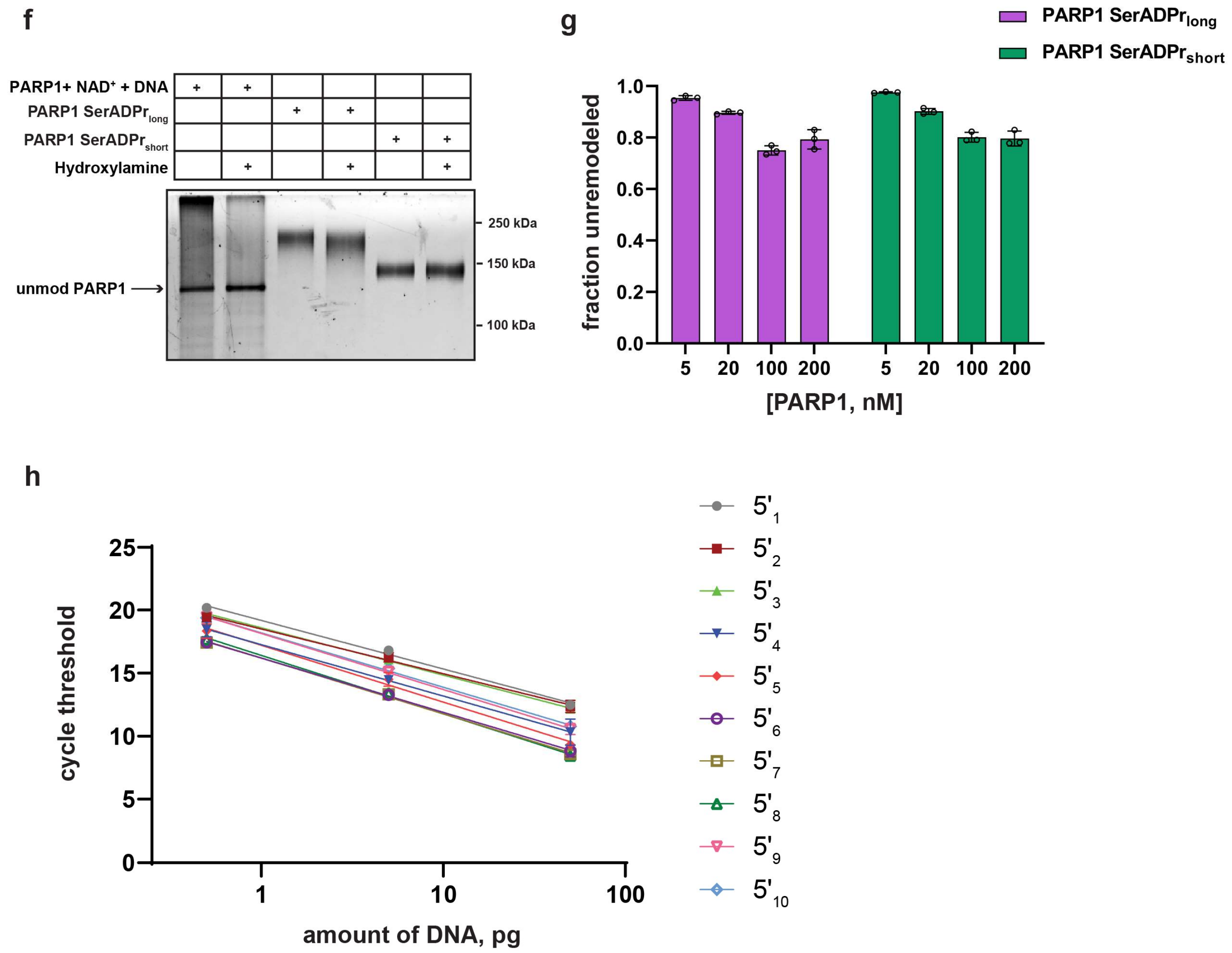


**Supplementary Fig. 5: Representative ALC1 remodeling time-course assay and additional controls to support that ALC1 directly remodels poly-ADP-ribosylated nucleosomes as demonstrated in Fig. 5**

**a**, Representative TBE gel analyses corresponding to recombinant ALC1 time course remodeling activity assay on the indicated nucleosome substrates and peptide additives where applicable. Densitometry values are included in Supplementary Dataset. EtBr = ethidium bromide stain. **b**, ALC1 nucleosome remodeling assay time-course wherein each reaction comprises ALC1 and the indicated nucleosome (‘unmod’= unmodified) (20 nM), and the indicated modified histone peptide or PARP1 and $NAD^+$ (2 mM). Modified histone peptide concentration is equal to the corresponding full-length histone concentration (40 nM). The H2BS6/H3S10ADPr4 nucleosome remodeling data is included for direct comparison. **c,** Single time-point (1 h) ALC1 nucleosome remodeling assay wherein each reaction comprises ALC1, an unmodified nucleosome substrate (20 nM), and the indicated ADP-ribosylated histone peptide concentration. Data are represented as mean ± s.d. (n = 3). **d,** PARP1 and ADP-ribose western blot analyses following an ALC1 remodeling reaction time-course that includes unmodified nucleosome, PARP1 (20 nM), and $NAD^+$. **e**, Single time-point (1 h) ALC1 nucleosome remodeling assay wherein each reaction comprises ALC1, an unmodified nucleosome substrate (20 nM), the indicated PARP1 concentration, and $NAD^+$ (2 mM). Data are represented as mean ± s.d. (n = 3). **f**, Silver-stained gel image depicting the effect of hydroxylamine treatment on ADP-ribosylated PARP1 modified under different conditions, with and without HPF1. **g,** Single time-point (1 h) ALC1 nucleosome remodeling assay wherein each reaction comprises ALC1, an unmodified nucleosome substrate (20 nM), and the indicated auto-modified PARP1 concentration. Data are represented as mean ± s.d. (n = 3). **h,** Primer efficiency curves corresponding to all primer:template pairs described in Fig. 5g,h.

**a**

| Identities: | Positives: | Gaps: |
|---|---|---|
| 214/470 (46%) | 298/470 (63%) | 37/470 (7%) |

```
ALC1   60    RFHCQNGC--ILGDEMGLGKTCQTIALFIY-LAGRLNDEGPFLILCPLSVLSNWKEEMQR  116
             RF    G   IL DEMGLGKT QT A+F+Y L    + +GPFL+  PLS + NW+ E +
CHD4   738   RFSWAQGTDTILADEMGLGKTVQT-AVFLYSLYKEGHSKGPFLVSAPLSTIINWEREFEM  796

ALC1   117   FAPGLSCVTYAGDKEERACLQQD-----------------LKQES--RFHVLLTTYEICL  157
             +AP +   VTY GDK+ RA ++++                 +K+E+  +FHVLLT+YE+
CHD4   797   WAPDMYVVTYVGDKDSRAIIRENEFSFEDNAIRGGKKASRMKKEASVKFHVLLTSYELIT  856

ALC1   158   KDASFLKSFPWSVLVVDEAHRLKNQSSLLHKTLSEFSVVFSLLLTGTPIQNSLQELYSLL  217
              D + L S  W+ L+VDEAHRLKN  S    + L+ +S+    LLLTGTP+QN+L+EL+ LL
CHD4   857   IDMAILGSIDWACLIVDEAHRLKNNQSKFFRVLNGYSLQHKLLLTGTPLQNNLEELFHLL  916

ALC1   218   SFVEPDLFSKEEVGDFIQRYQDIEKESESASELHKLLQPFLLRRVKAEVATELPKKTEVV  277
             +F+ P+ F   E   F++ + DI KE +    +LH +L P +LRR+KA+V    +P KTE++
CHD4   917   NFLTPERFHNLE--GFLEEFADIAKEDQ-IKKLHDMLGPHMLRRLKADVFKNMPSKTELI  973

ALC1   278   IYHGMSALQKKYYKAILMKDLDAFENETA-KKVKLQNILSQLRKCVDHPYLFDGVE----  332
             +   +S +QKKYYK IL ++ +A         +V L N++  L+KC +HPYLF
CHD4   974   VRVELSPMQKKYYKYILTRNFEALNARGGGNQVSLLNVVMDLKKCCNHPYLFPVAAMEAP  1033

ALC1   333   --PEPFEVGDHLTEASGKLHLLDKLLAFLYSGGHRVLLFSQMTQMLDILQDYMDYRGYSY  390
               P     G  L  ASGKL LL K+L  L  GGHRVL+FSQMT+MLD+L+D++++ GY Y
CHD4   1034  KMPNGMYDGSALIRASGKLLLLQKMLKNLKEGGHRVLIFSQMTKMLDLLEDFLEHEGYKY  1093

ALC1   391   ERVDGSVRGEERHLAIKNF---GQQPIFVFLLSTRAGGVGMNLTAADTVIFVDSDFNPQN  447
             ER+DG + G  R  AI  F   G Q  F FLLSTRAGG+G+NL  ADTVI  DSD+NP N
CHD4   1094  ERIDGGITGNMRQEAIDRFNAPGAQQ-FCFLLSTRAGGLGINLATADTVIIYDSDWNPHN  1152

ALC1   448   DLQAAARAHRIGQNKSVKVIRLIGRDTVEEIVYRKAASKLQLTNMIIEGG  497
             D+QA +RAHRIGQNK V + R + R +VEE + + A  K+ LT++++  G
CHD4   1153  DIQAFSRAHRIGQNKKVMIYRFVTRASVEERITQVAKKKMMLTHLVVRPG  1202
```

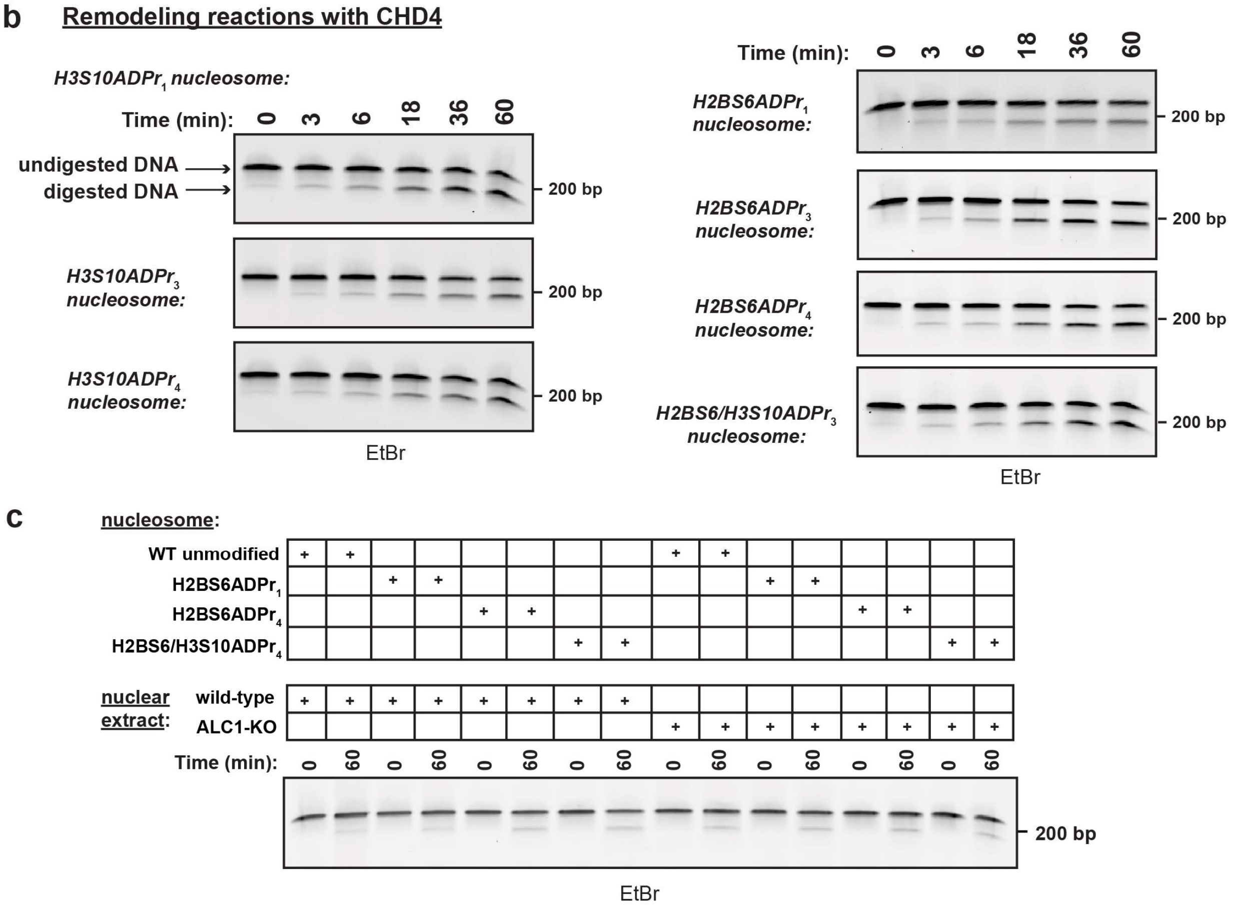

**d**

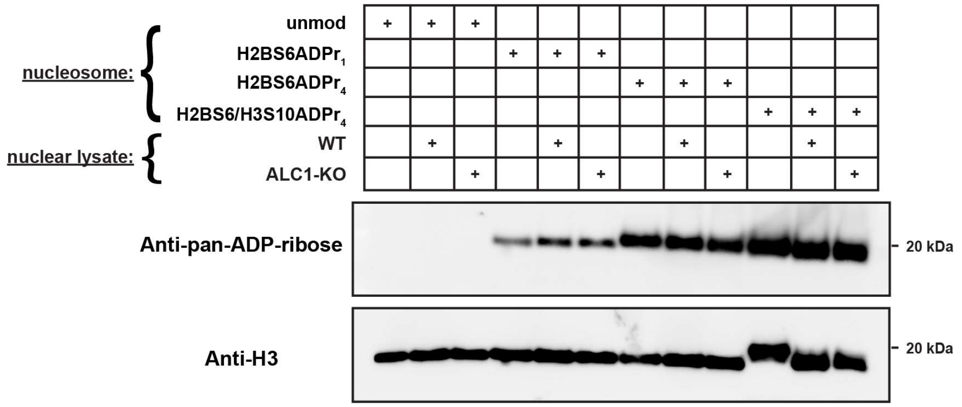


**Supplementary Fig. 6: H3S10ADPr and H2BS6ADPr specifically sensitize nucleosomes to chromatin remodeling by ALC1**

**a**, BLAST alignment of the ATPase domains from ALC1 (amino acids 60-497) and CHD4 (amino acids 738-1202). **b**, Representative TBE gel analyses corresponding to recombinant CHD4 time course remodeling activity assay on the indicated nucleosome substrates. Densitometry values are included in Supplementary Dataset. EtBr = ethidium bromide stain. **c**, Representative TBE gel analyses corresponding to wild-type or ALC1 knock-out (KO) HEK293T nuclear extract chromatin remodeling activity on the indicated nucleosome substrates. Densitometry values are included in Supplementary Dataset. EtBr = ethidium bromide stain. **d,** Western blot showing the effect of incubating nucleosomes with or without nuclear lysate on the ADPr status of the nucleosomes.

#### Supplementary Table 1

##### Affinities of Af1521 macrodomain* for ADP-ribosylated histone peptides

| peptide | $K_{d, app}$ (nM) | 95% CI | $R^2$ |
|---|---|---|---|
| H3 unmodified | n.d. | n.d. | 0.17 |
| $H3S10ADPr_1$ | 1301 | 987.7 to 1740 | 0.98 |
| $H3S10ADPr_2$ | 762.4 | 690.6 to 842.4 | 0.99 |
| $H3S10ADPr_3$ | 64.56 | 54.55 to 76.31 | 0.98 |
| $H3S10ADPr_4$ | 65.43 | 53.46 to 79.95 | 0.98 |
| $H3S10ADPr_5$ | 42.51 | 33.80 to 53.31 | 0.97 |
| H2B unmodified | n.d. | n.d. | 0.49 |
| $H2BS6ADPr_1$ | 1642 | 1411 to 1922 | 0.99 |
| $H2BS6ADPr_2$ | 526.5 | 484.5 to 572.4 | 0.99 |
| $H2BS6ADPr_3$ | 84.78 | 71.67 to 100.2 | 0.99 |
| $H2BS6ADPr_4$ | 96.35 | 77.05 to 120.2 | 0.98 |

##### Affinities of ALC1 macrodomain for ADP-ribosylated histone peptides

| peptide | $K_{d, app}$ (nM) | 95% CI | $R^2$ |
|---|---|---|---|
| H3 unmodified | n.d. | n.d. | 0.28 |
| $H3S10ADPr_1$ | n.d. | n.d. | 0.36 |
| $H3S10ADPr_2$ | n.d. | n.d. | 0.40 |
| $H3S10ADPr_3$ | 21.22 | 17.06 to 26.33 | 0.98 |
| $H3S10ADPr_4$ | 27.24 | 19.12 to 38.69 | 0.94 |
| $H3S10ADPr_5$ | 27.93 | 22.43 to 34.74 | 0.98 |
| H2B unmodified | n.d. | n.d. | 0.29 |
| $H2BS6ADPr_1$ | n.d. | n.d. | 0.11 |
| $H2BS6ADPr_2$ | 4909 | 2080 to 46956 # | 0.91 |
| $H2BS6ADPr_3$ | 30.64 | 23.17 to 40.38 | 0.96 |
| $H2BS6ADPr_4$ | 36.95 | 28.18 to 48.30 | 0.96 |

* : the Af1521 macrodomain is from the commercially available anti-pan-ADP-ribose binding reagent and has the immunoglobulin Fc fused to it.
n.d. : indicates that the value could not be determined with reliability because of a poor fit to the non-linear regression model being used.
### : indicates that the value is a 90% CI instead of 95%, because the 95% CI could not be assigned a numerical limit.

Abbreviations
CI : confidence interval
$R^2$ : r-squared value or the coefficient of determination
app : apparent

#### Supplementary Table 2

##### Chromatin remodeling rate constants for single-substrate assays with ALC1

| nucleosome | k ($min^{-1}$) | 95% CI | $R^2$ | |
|---|---|---|---|---|
| unmodified | 0.0001379 | 8.073e-005 to 0.0001952 | 0.62 | |
| $H3S10ADPr_1$ | 0.0003232 | 5.836e-005 to 0.0005893 | 0.30 | $ |
| $H3S10ADPr_3$ | 0.01002 | 0.009478 to 0.01057 | 0.99 | |
| $H3S10ADPr_4$ | 0.01037 | 0.008533 to 0.01231 | 0.92 | |
| $H2BS6ADPr_1$ | 0.0001646 | 0.0001004 to 0.0002290 | 0.65 | |
| $H2BS6ADPr_3$ | 0.01536 | 0.01312 to 0.01775 | 0.95 | |
| $H2BS6ADPr_4$ | 0.02776 | 0.02207 to 0.03464 | 0.93 | |
| $H2BS6/H3S10ADPr_3$ | 0.04024 | 0.03546 to 0.04566 | 0.98 | |
| $H2BS6/H3S10ADPr_4$ | 0.05152 | 0.04040 to 0.06597 | 0.95 | |
| unmodified+$H3S10ADPr_4$ | 8.323e-005 | 3.061e-005 to 0.0001359 | 0.41 | $ |
| unmodified+$H2BS6ADPr_4$ | 1.886e-005 | 0 to 8.611e-005 | 0.02 | $ |
| unmodified+PARP1 (20 nM) | 0.001667 | 0.001166 to 0.002173 | 0.76 | |
| unmodified+PARP1 (100 nM) | 0.001970 | 0.001518 to 0.002425 | 0.85 | |
| unmodified+PARP1 $SerADPr_{long}$ | 0.005027 | 0.004190 to 0.005880 | 0.92 | |
| unmodified+PARP1 $SerADPr_{short}$ | 0.003826 | 0.003256 to 0.004403 | 0.93 | |

##### Chromatin remodeling rate constants for single-substrate assays with CHD4

| nucleosome | k ($min^{-1}$) | 95% CI | $R^2$ |
|---|---|---|---|
| unmodified | 0.01010 | 0.008834 to 0.01142 | 0.96 |
| $H3S10ADPr_1$ | 0.01049 | 0.008671 to 0.01240 | 0.92 |
| $H3S10ADPr_3$ | 0.008974 | 0.007868 to 0.01011 | 0.96 |
| $H3S10ADPr_4$ | 0.009624 | 0.008744 to 0.01053 | 0.98 |
| $H2BS6ADPr_1$ | 0.007561 | 0.006142 to 0.009034 | 0.90 |
| $H2BS6ADPr_3$ | 0.01141 | 0.009590 to 0.01333 | 0.93 |
| $H2BS6ADPr_4$ | 0.01092 | 0.008884 to 0.01307 | 0.91 |
| $H2BS6/H3S10ADPr_3$ | 0.009619 | 0.008354 to 0.01093 | 0.95 |
| $H2BS6/H3S10ADPr_4$ | 0.007857 | 0.006470 to 0.009295 | 0.91 |

$ : indicates that the data did not fit well with the non-linear regression model being used but the values have been plotted in the associated figure (Figure 5E).

#### Supplementary Table 3

##### Chromatin remodeling rate constants for multi-substrate assays with ALC1

###### H3 substrate pool

| nucleosome | k ($min^{-1}$) | 95% CI | $R^2$ |
|---|---|---|---|
| unmodified ($5'_1$) | n.d. | n.d. | -0.11 |
| unmodified ($5'_9$) | n.d. | n.d. | 0.15 |
| $H3S10ADPr_1$ | n.d. | n.d. | -0.23 |
| $H3S10ADPr_3$ | 0.03784 | 0.02821 to 0.05097 | 0.91 |
| $H3S10ADPr_4$ | 0.03280 | 0.02194 to 0.04905 | 0.84 |

###### H2B substrate pool

| nucleosome | k ($min^{-1}$) | 95% CI | $R^2$ |
|---|---|---|---|
| unmodified ($5'_1$) | n.d. | n.d. | -0.35 |
| unmodified ($5'_9$) | n.d. | n.d. | 0.21 |
| $H2BS6ADPr_1$ | n.d. | n.d. | 0.06 |
| $H2BS6ADPr_3$ | 0.03637 | 0.02566 to 0.05170 | 0.88 |
| $H2BS6ADPr_4$ | 0.06686 | 0.04344 to 0.1042 | 0.90 |

n.d. : indicates that the value could not be determined with reliability because of a poor fit to the non-linear regression model being used.

#### Supplementary Table 4

##### DNA oligonucleotides described in this study

| Identifier | Sequence (5'→3') |
|---|---|
| ALC1-KO gRNA1 | GTCGCCTGCATATGTTACAC |
| ALC1-KO gRNA2 | GACCACCTGACTGAGGCTAG |
| PARP stimulating DNA | GCTGGTTCGCGAACCAGC |
| 601 amplification 45-Forward | GGCCGCTCTAGAACTAGTGG |
| 601 amplification 8-Reverse | CTCTGATGCTGGAGAATCC |
| 601 amplification 5′ 1 | GCCATCACGCCACAGTTTCGATCCGATATCGCTGTTCACC |
| 601 amplification 5′ 2 | CGCTGACGCACTCAAATGCGATCCGATATCGCTGTTCACC |
| 601 amplification 5′ 3 | CGGCTAAGAGTAGGTTGGGGATCCGATATCGCTGTTCACC |
| 601 amplification 5′ 4 | CTTACTAAGGCCATCGCGGGATCCGATATCGCTGTTCACC |
| 601 amplification 5′ 5 | CACGATTCAACTACGCCGCGATCCGATATCGCTGTTCACC |
| 601 amplification 5′ 6 | CGTGACGACGTTCCTGCTAGATCCGATATCGCTGTTCACC |
| 601 amplification 5′ 7 | CGCTGCAGACACTATACCGGATCCGATATCGCTGTTCACC |
| 601 amplification 5′ 8 | GCAGCGACATCCACTTGAGGATCCGATATCGCTGTTCACC |
| 601 amplification 5′ 9 | CAGCAGTTCGTCTCTGCTGGATCCGATATCGCTGTTCACC |
| 601 amplification 5′ 10 | CAAGACTCGCCTTACGGCTGATCCGATATCGCTGTTCACC |
| qPCR 5′ 1 | GCCATCACGCCACAGTTTC |
| qPCR 5′ 2 | CGCTGACGCACTCAAATGC |
| qPCR 5′ 3 | CGGCTAAGAGTAGGTTGGG |
| qPCR 5′ 4 | CTTACTAAGGCCATCGCGG |
| qPCR 5′ 5 | CACGATTCAACTACGCCGC |
| qPCR 5′ 6 | CGTGACGACGTTCCTGCTA |
| qPCR 5′ 7 | CGCTGCAGACACTATACCG |
| qPCR 5′ 8 | GCAGCGACATCCACTTGAG |
| qPCR 5′ 9 | CAGCAGTTCGTCTCTGCTG |
| qPCR 5′ 10 | CAAGACTCGCCTTACGGCT |
| Universal qPCR-Reverse | CTCTGATGCTGGAGAATCCCG |

*Note: the underlined sequence refers to the unique 5′ priming site included for each 601 DNA fragment prepared for the nucleosome remodeling competition assay.

**Supplementary Table 5**

**Antibodies used in this study**

| ANTIBODY | SOURCE | IDENTIFIER | DILUTION |
|---|---|---|---|
| IRDye 800CW Goat anti-Rabbit IgG (H+L) | Thermo-Fisher Scientific | NC9401842 | 1:5000 |
| IRDye 680RD Goat anti-Mouse IgG (H+L) | Thermo-Fisher Scientific | NC0252290 | 1:5000 |
| Anti-pan-ADP-ribose binding reagent | Millipore Sigma | MABE1016 | 1:1000 |
| Histone H3 (D1H2) XP® Rabbit mAb #4499 | Cell Signaling Technologies | 4499S | 1:8000 |
| Histone H2B (D2H6) Rabbit mAb #12364 | Cell Signaling Technologies | 12364S | 1:2000 |
| Histone H2A (L88A6) Mouse mAb #3636 | Cell Signaling Technologies | 3636A | 1:600 |
| PARP (46D11) Rabbit mAb #9532 | Cell Signaling Technologies | 9532S | 1:1000 |
| CHD1L (E1I8C) Rabbit mAb #13460 | Cell Signaling Technologies | 13460S | 1:1000 |
| CHD1 (D8C2) Rabbit mAb #4351 | Cell Signaling Technologies | 4351S | 1:1000 |
| SNF2H (D4W6N) Rabbit mAb #38410 | Cell Signaling Technologies | 38410S | 1:1000 |
| CHD4 (D8B12) Rabbit mAb #11912 | Cell Signaling Technologies | 11912S | 1:1000 |
| Brg1 (P680) Rabbit Antibody #3514 | Cell Signaling Technologies | 3514S | 1:1000 |
| Anti-histone H3 antibody mouse ab10799 | Cell Signaling Technologies | 3514S | 1:1000 |
